# Evolutionary genomics of the emergence of brown algae as key components of coastal ecosystems

**DOI:** 10.1101/2024.02.19.579948

**Authors:** France Denoeud, Olivier Godfroy, Corinne Cruaud, Svenja Heesch, Zofia Nehr, Nachida Tadrent, Arnaud Couloux, Loraine Brillet-Guéguen, Ludovic Delage, Dean Mckeown, Taizo Motomura, Duncan Sussfeld, Xiao Fan, Lisa Mazéas, Nicolas Terrapon, Josué Barrera-Redondo, Romy Petroll, Lauric Reynes, Seok-Wan Choi, Jihoon Jo, Kavitha Uthanumallian, Kenny Bogaert, Céline Duc, Pélagie Ratchinski, Agnieszka Lipinska, Benjamin Noel, Eleanor A. Murphy, Martin Lohr, Ananya Khatei, Pauline Hamon-Giraud, Christophe Vieira, Svea Sanja Akerfors, Shingo Akita, Komlan Avia, Yacine Badis, Tristan Barbeyron, Arnaud Belcour, Wahiba Berrabah, Samuel Blanquart, Ahlem Bouguerba-Collin, Trevor Bringloe, Rose Ann Cattolico, Alexandre Cormier, Helena Cruz de Carvalho, Romain Dallet, Olivier De Clerck, Ahmed Debit, Erwan Denis, Christophe Destombe, Erica Dinatale, Simon Dittami, Elodie Drula, Sylvain Faugeron, Jeanne Got, Louis Graf, Agnès Groisillier, Marie-Laure Guillemin, Lars Harms, William John Hatchett, Bernard Henrissat, Galice Hoarau, Chloé Jollivet, Alexander Jueterbock, Ehsan Kayal, Andrew H. Knoll, Kazuhiro Kogame, Arthur Le Bars, Catherine Leblanc, Line Le Gall, Ronja Ley, Xi Liu, Steven T. LoDuca, Pascal Jean Lopez, Philippe Lopez, Eric Manirakiza, Karine Massau, Stéphane Mauger, Laetitia Mest, Gurvan Michel, Catia Monteiro, Chikako Nagasato, Delphine Nègre, Eric Pelletier, Naomi Phillips, Philippe Potin, Stefan A. Rensing, Ellyn Rousselot, Sylvie Rousvoal, Declan Schroeder, Delphine Scornet, Anne Siegel, Leila Tirichine, Thierry Tonon, Klaus Valentin, Heroen Verbruggen, Florian Weinberger, Glen Wheeler, Hiroshi Kawai, Akira F. Peters, Hwan Su Yoon, Cécile Hervé, Naihao Ye, Eric Bapteste, Myriam Valero, Gabriel V. Markov, Erwan Corre, Susana M. Coelho, Patrick Wincker, Jean-Marc Aury, Mark Cock

## Abstract

Brown seaweeds are keystone species of coastal ecosystems, often forming extensive underwater forests, that are under considerable threat from climate change. Despite their ecological and evolutionary importance, this phylogenetic group, which is very distantly related to animals and land plants, is still poorly characterised at the genome level. Here we analyse 60 new genomes that include species from all the major brown algal orders. Comparative analysis of these genomes indicated the occurrence of several major events coinciding approximately with the emergence of the brown algal lineage. These included marked gain of new orthologous gene families, enhanced protein domain rearrangement, horizontal gene transfer events and the acquisition of novel signalling molecules and metabolic pathways. The latter include enzymes implicated in processes emblematic of the brown algae such as biosynthesis of the alginate-based extracellular matrix, and halogen and phlorotannin biosynthesis. These early genomic innovations enabled the adaptation of brown algae to their intertidal habitats. The subsequent diversification of the brown algal orders tended to involve loss of gene families, and genomic features were identified that correlated with the emergence of differences in life cycle strategy, flagellar structure and halogen metabolism. Analysis of microevolutionary patterns within the genus *Ectocarpus* indicated that deep gene flow between species may be an important factor in genome evolution on more recent timescales. Finally, we show that integration of large viral genomes has had a significant impact on brown algal genome content and propose that this process has persisted throughout the evolutionary history of the lineage.

## INTRODUCTION

The brown algae (Phaeophyceae) are a lineage of complex multicellular organisms that emerged about 450 Mya^1^ from within a group of photosynthetic stramenopile taxa (derived from a secondary endosymbiosis involving a red alga^2^) that are either unicellular or have very simple filamentous multicellular thalli (Figure 1). The emerging brown algae acquired a number of characteristic features that are thought to have contributed to the evolutionary success of this lineage, including complex polysaccharide-based cell walls that confer protection and flexibility in the highly dynamic intertidal environment^3^, complex halogen^4^ and phlorotannin^5^ metabolisms that are thought to play important roles in multiple processes including defence, adhesion and cell wall modification, and a remarkable diversity of life cycles and developmental body architectures adapted to diverse marine environments^6^. As a result of these attributes, many brown algae have become established as key components of extensive coastal ecosystems. These seaweed-based ecosystems provide high value Earth-system-scale services, including the sequestration of several megatons of carbon per year globally, comparable to values reported for terrestrial forests^7^, but this important role of seaweed ecosystems is threatened by climate-related declines in seaweed populations worldwide^8^. However, appropriate conservation measures, coupled with the development of seaweed mariculture as a highly sustainable and low impact approach to food and biomass production, could potentially reverse this trend, allowing seaweeds to play a significant role in mitigating the effects of climate change^9^. To attain this objective, it will be necessary to address important gaps in our knowledge of the biology and evolutionary history of the brown algal lineage. For example, these seaweeds remain poorly described in terms of genome sequencing due, in part, to difficulties with extracting nucleic acids. The Phaeoexplorer project (https://phaeoexplorer.sb-roscoff.fr/) has generated a large dataset of genome sequences, spanning all the major orders of the Phaeophyceae^10^. This extensive genomic dataset has been analysed here to study the origin and evolution of key genomic features during the emergence and diversification of this important group of marine organisms.

**Figure 1.**
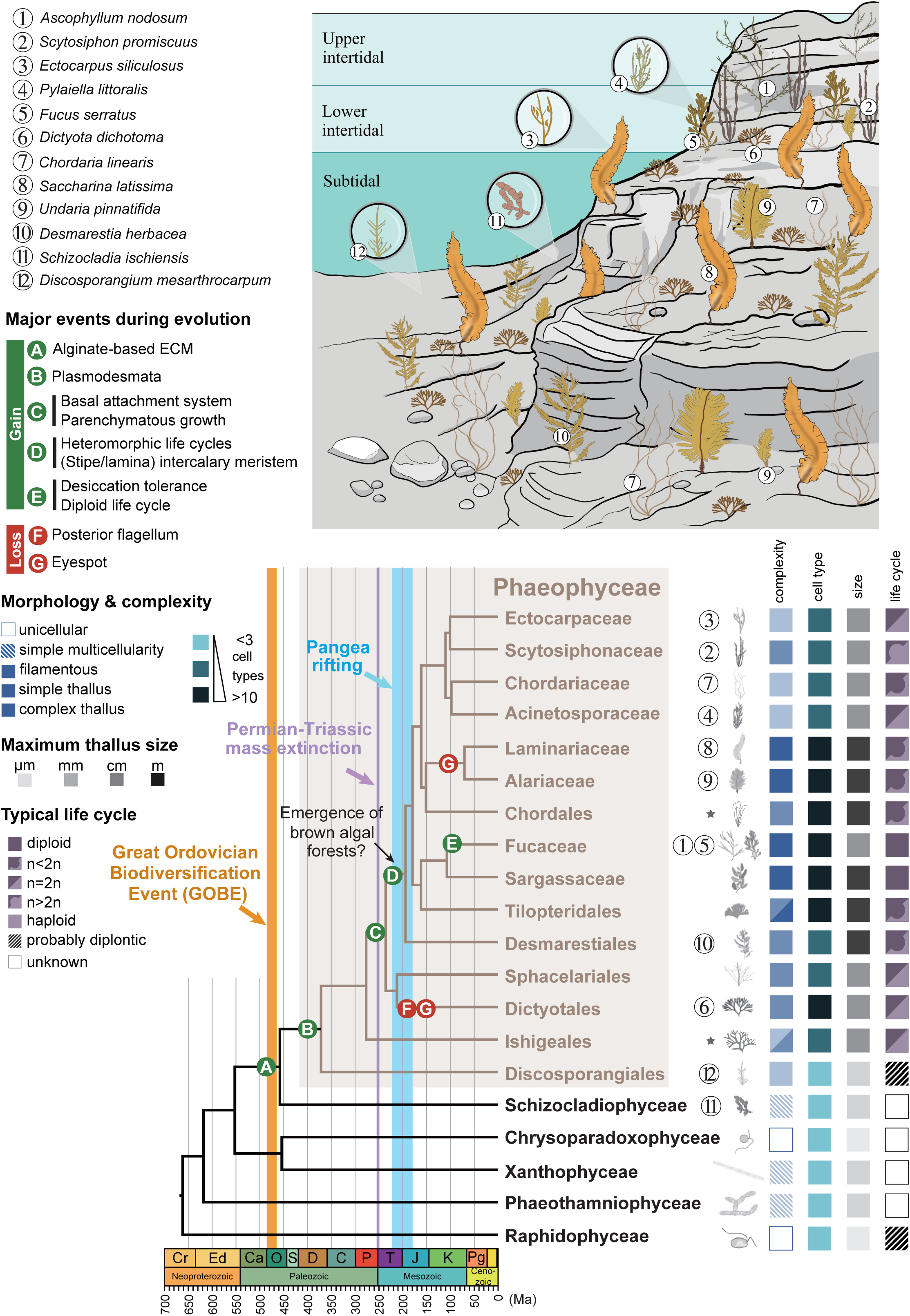
Ecology, diversity and evolutionary features of the brown algae. The upper panel indicates approximate positions in the intertidal of key species whose genomes have been sequenced by the Phaeoexplorer project. The lower panel illustrates the diversity of brown algae in terms of number of cell types, thallus size, morphological complexity and life cycle type (maximal values for each taxa). The panel also indicates a number of key evolutionary events that occurred during the emergence of the Phaeophyceae and diversification of the brown algal orders. Some lineages may have secondarily lost a characteristic after its acquisition. Note that members of the genus *Ishige* (Ishigeaceae) also exhibit desiccation tolerance (not shown). *, these orders were not analysed in this study; Cr, Cryogenian; Ed, Ediacaran; Ca, Cambrian; O, Ordovician; S, Silurian; D, Devonian; C, Carboniferous; P, Permian; T, Triassic; J, Jurassic; K, Cretaceous; Pg, Paleogene.

## RESULTS

### In-depth sequencing of brown algal genomes

Until now, good quality genome assemblies have been obtained for only five brown algal species^11–15^, together with about 46 draft genome assemblies^16–20^. Here we report work that has significantly expanded the genomic data available by sequencing and assembling 17 good quality genomes using long-read technology (Table S1), plus an additional 43 draft genome assemblies. These 60 genomes correspond to 40 brown algae and four closely related species, covering 16 Phaeophyceae families providing a dense coverage of this lineage (Figures S1A, S1B, Table S1). The sequenced species include brown algae that occur at different levels of the intertidal and subtidal, and are representative of the broad diversity of this group of seaweeds in terms of size, levels of multicellular complexity, biogeography and life cycle structure (Figures 1, S2). The analyses carried out in this study have focused principally on a set of 21 good quality reference genomes, which include four previously published genomes (Table S1).

### Marked changes in genome content and gene structure during the emergence of the Phaeophyceae lineage

Recent evidence indicates that the brown algae emerged about 450 Mya during the Great Ordovician Biodiversification Event (GOBE)^1^, a conclusion that is supported by a fossil-calibrated tree built with a nuclear-gene-based phylogeny constructed using the Phaeoexplorer genome data (Figures 1, S3). An increase in atmospheric oxygen at the time of the GOBE, associated with the emergence of herbivorous marine invertebrates^21^, is likely to have created conditions conducive to the observed transition towards increased multicellular complexity during early brown algal evolution.

To investigate genomic modifications associated with the emergence and diversification of the brown algae, we first carried out a series of genome-wide analyses aimed at identifying broad trends in genome evolution over evolutionary time (Figure 2). Dollo analysis of gain and loss of orthogroups (i.e. gene families) indicated marked gains during early brown algal evolution followed by a broad tendency to lose orthogroups later as the different brown algal orders diversified (Figures 2B, S4). Similarly, a phylostratigraphy analysis indicated that 29.6% of brown algal genes cannot be traced back to ancestors outside the Phaeophyceae, with the majority of gene founder events occurring early during the emergence of the brown algae (Figure 2E, S5A, Table S2), again indicating a burst of gene birth during the emergence of this lineage. Both the Dollo analysis and the phylostratigraphy approach indicated that the gene families acquired during early brown algal evolution were significantly enriched in genes that could not be assigned to a COG category, suggesting a burst in the acquisition of genetic novelty (Figure 2G).

**Figure 2.**
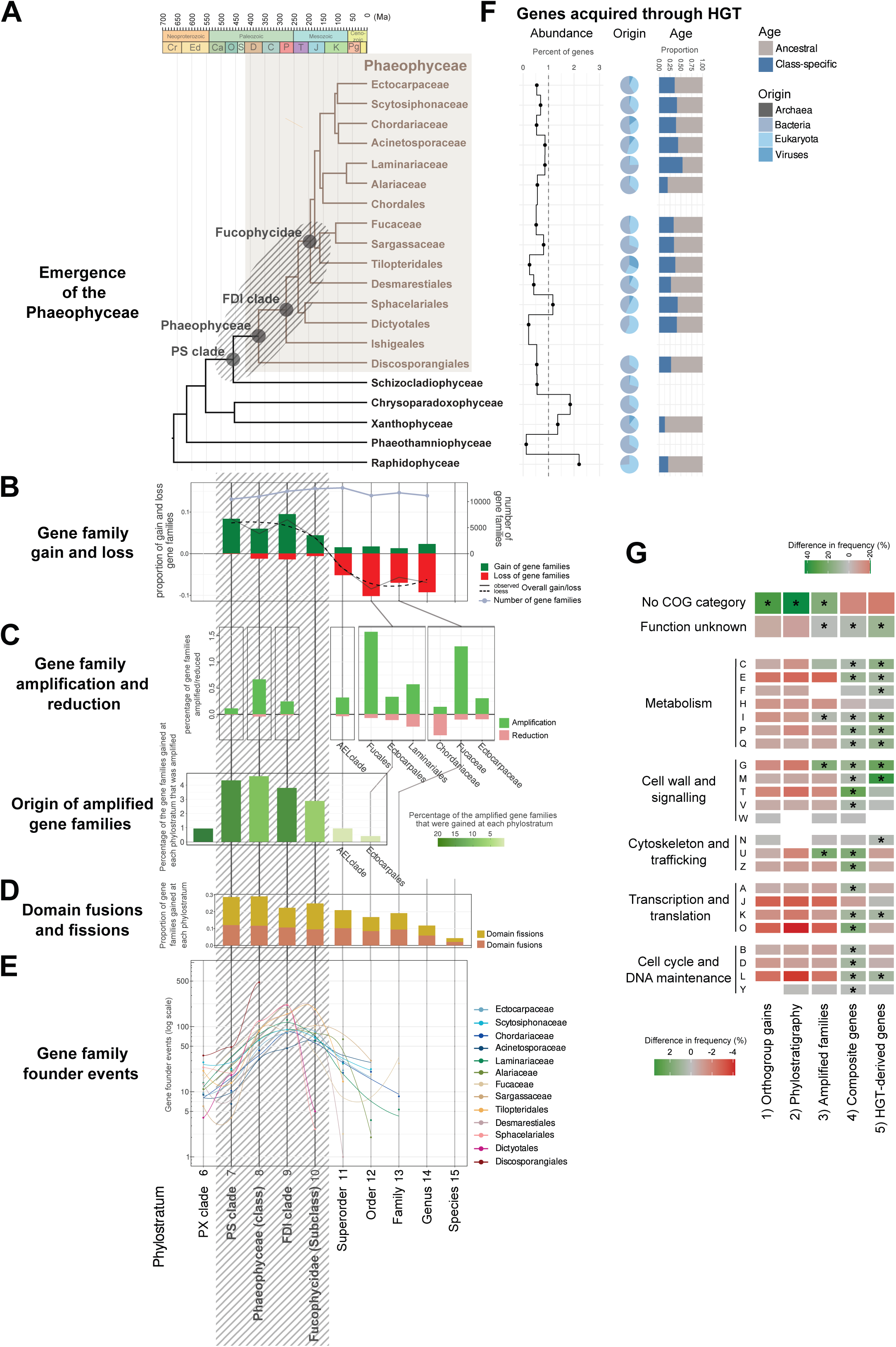
Genome-wide analyses of brown algal genome and gene content evolution. (A) Time-calibrated cladogram based on Figure S3 with Phaeophyceae in brown and outgroup species in black. The grey hatched area, which indicates key nodes corresponding to the origin and early emergence of the brown algae, is mirrored in (B) to (F). (B) Gene family (orthogroup) gain (green) and loss (red) during the emergence and diversification of the brown algae based on a Dollo parsimony reconstruction (Figure S4). (C) Upper panel: timing of gene family (orthogroup) amplification and reduction during the evolutionary history of the Phaeophyceae (CAFE5 analysis). Lower panel: time of origin (orthogroup gain, based on the Dollo parsimony reconstruction) of the 180 most strongly amplified gene families. (D) Composite gene analysis. Proportions of gene families showing domain fusion (orange) or domain fission (yellow) at different age strata. (E) Inferred gene family founder events after accounting for homology detection failure. (F) Horizontal-gene-transfer-derived genes in orthologous groups and across species. Taxa along the y-axis are as in (A). Grey bars indicate the number of HGT genes (light-grey) and the number of orthogroups containing HGT genes (dark-grey) for each species. The black trace represents the percentage of genes resulting from HGT events per species. Pie charts summarise the predicted origins (donor taxa) of the HGT genes. The right-hand bar graph indicates the proportions of ancestral (i.e. acquired before the root of the phylogenetic class, in grey) and class-specific (i.e. acquired within the phylogenetic class, in blue) HGT genes. (G) Enrichment of COG categories in sets of gene families identified as being 1) gained at the four indicated early nodes by the Dollo analysis, 2) gene founder events at the four indicated phylostrata, 3) amplified in the Phaeophyceae (180 most strongly amplified families), 4) domain fusions or fissions and 5) HGT-derived. Asterisks indicate significantly enriched categories compared to the entire set of gene families. FDI clade, Fucophycideae/Dictyotales/Ishigeales; PS clade, Phaeophyceae plus Schizocladiophyceae; PX clade, Phaeophyceae plus Xanthophyceae.

One of the factors underlying the marked burst of gene gain during the emergence of the brown algae was an increase in the rate of acquisition of new genes via horizontal gene transfer (HGT). A phylogeny-based search for genes potentially derived from HGT events indicated that they constitute about 1% of brown algal gene catalogues and that the novel genes were principally acquired from bacterial genomes (Figures 2F, S5B). The proportion of class-specific HGT events (i.e. events where a gene was acquired via HGT by an ancestral acceptor species after the emergence of a taxonomic class so that the gene is only found in members of that class) compared to more ancient HGT events was greater for the brown algae (33.5%) than for the closely-related taxa Xanthophyceae (*Tribonema minus*) and Raphidophyceae (*Heterosigma akashiwo*; mean of 17.1% for the two taxa, Wilcoxon *p*-value 0.021), indicating that higher levels of HGT occurred during the emergence of the brown algae than in closely-related taxa (Figure 2F).

The marked increase in the rate of gene gain appears to have been a key factor in the emergence of the brown algal lineage but this was not the only process that enriched brown algal genomes during this period. Domain fusions and fissions (composite genes) were prevalent during the early stages of brown algal emergence (with gene fusion events being slightly more common than gene fissions; Figures 2D, S5C), affecting about 7% of brown algal gene complements. In contrast, gene family amplifications were most prevalent at a later stage of brown algal evolution, corresponding to the diversification of the major brown algal orders during the Mesozoic (Figures 2C, S5D, S5E, Table S3), and therefore did not share the same pattern of increased occurrence during early evolution of the lineage. However, the amplified gene families were significantly enriched in genes that had been gained during the emergence of the brown algae (χ^2^ *p*-value 1.04e-15; Table S1C) indicating that gene gain during the early evolution of the lineage nonetheless played a crucial role by establishing the majority of the gene families that would later undergo amplifications.

Analysis of the predicted functions of the three sets of gene families identified as having been amplified, derived from domain fusions/fissions or derived from an HGT event (Figure 2G) indicated that they were enriched in several functional categories, notably carbohydrate metabolism, signal transduction and transcription. These functional categories may have been important in the emergence of the complex brown algal cell wall or correspond to a complexification of signalling pathways as multicellular complexity increased. Interestingly, many of the genes acquired at, or shortly after, the origin of the Phaeophyceae encode secreted or membrane proteins (Figure S5A), suggesting roles in cell-cell communication that may have been important for the emergence of complex multicellularity or as components of defence mechanisms. The acquisition of plasmodesmata by brown algae directly after their divergence from their sister taxon *Schizocladia ischiensis*^22^ (Figure 1) underlines the importance of cell-cell communication from the outset of brown algal evolution.

The emergence of the brown algae also corresponded with changes in gene structure. On average, brown algal genes tend to be more intron-rich than those of the other stramenopile groups^23^, including closely-related taxa (Figures S6A), with the notable exception of *Chrysoparadoxa australica*, whose genes contain many short, C/T-rich introns (Figure S6A). A comparison of orthologous genes indicated a phase of rapid intron acquisition just before the divergence of the Phaeophyceae and the Schizocladiophyceae, followed by a period of relative intron stability up to the present day (Figure S6B). This phase of accelerated intron acquisition coincided approximately with the periods of marked gene gain and domain reorganisation discussed above and may have been an indirect consequence of increased multicellular complexity (Figure 1) due to a concomitant decrease in effective population size^24^. Once established, increased intron density may have facilitated some of the genome-wide tendencies described above, such as increased reorganisation of composite genes for example, and thereby played an important role in a context of increasing developmental complexity^25–28^.

Taken together, these analyses suggest a chain of events in which GOBE increases in both atmospheric oxygen and herbivory drove the emergence of the brown algae, and the acquisition of multicellular complexity by this emerging lineage. The emergence of this lineage was associated with marked changes in the general structure and content of their genomes, including the acquisition of new genes (some via HGT) and reorganisation of protein domains within genes. The following section focuses on metabolic and signalling pathways that are predicted to have played key roles during this important period of emergence and early evolution of the brown algae.

### Acquisition of key metabolic and signalling pathways during the emergence of the Phaeophyceae

The success of the brown algae as an evolutionary lineage has been attributed, at least in part, to the acquisition of several key metabolic pathways, particularly those associated with cell wall biosynthesis, and both halogen and phlorotannin metabolism^3–5^. AuCoMe^29^ was used to provide an overview of metabolic pathways across the brown algae by inferring genome-scale metabolic networks for all the species in the Phaeoexplorer dataset (https://gem-aureme.genouest.org/phaeogem/; Figures S7A, S7B) and also for a subset of long-read sequenced species (https://gem-aureme.genouest.org/16bestgem/; Figures S7C, S7D). Comparison of the brown algal metabolic networks with those of closely-related taxa allowed the identification of enzymatic activities that have been gained or lost in the former compared with the latter (Figures S7E, S7F, S7G). The most robust set of core reactions found mainly in brown algae is a set of 24 reactions catalysed by polyamine oxidases, carotenoid and indole-3-pyruvate oxygenases, a methyladenine demethylase, a PQQ-dependent sugar dehydrogenase, a xylosyl transferase (CAZyme family GT14), a ribulosamine kinase, galactosylceramide sulfotransferases and phosphopentomutases (Table S4).

The identification of carbohydrate-active enzymes (CAZymes) among the AuCoMe core reaction set is of particular interest because many of these enzymes have been implicated in the biosynthesis of the characteristic cell walls of brown algae. Large complements of CAZYme genes (237 genes on average) were found in all brown algal orders and in their sister taxon *S. ischiensis* but this class of gene was less abundant in the more distantly related unicellular alga *H. akashiwo* (Figures 3A, S8A, S8B, Tables S5, S6). The evolutionary history of carbohydrate metabolism gene families was investigated by combining information from the genome-wide analyses of gene gain/loss, HGT and gene family amplification (Figure 3B). This analysis indicated that several key genes and gene families (ManC5-E, GT23 and PL41) were acquired by the common ancestor of brown algae and *S. ischiensis*, with strong evidence in some cases that this occurred via HGT (PL41). Moreover, marked amplifications were detected for several families (AA15, ManC5-E, GH114, GT23 and PL41), indicating that both gain and amplification of gene families played important roles in the emergence of the brown algal carbohydrate metabolism gene set. Alginate is a major component of brown algal cell walls and it plays an important role in conferring resistance to the biomechanical effect of wave action^3^. It is therefore interesting that mannuronan C5 epimerase (ManC5-E), an enzyme whose action modulates the rigidity of the alginate polymer, appears to have been acquired very early, by the common ancestor of brown algae and *S. ischiensis* (Figures 3B, 3C). The acquisition of ManC5-E, together with other alginate pathway enzymes such as PL41 (Figures 3A, 3B and 3D), was probably an important evolutionary step, enabling the emergence of large, resilient substrate-anchored multicellular organisms in the highly dynamic and stressful coastal environment (Figure 1).

**Figure 3.**
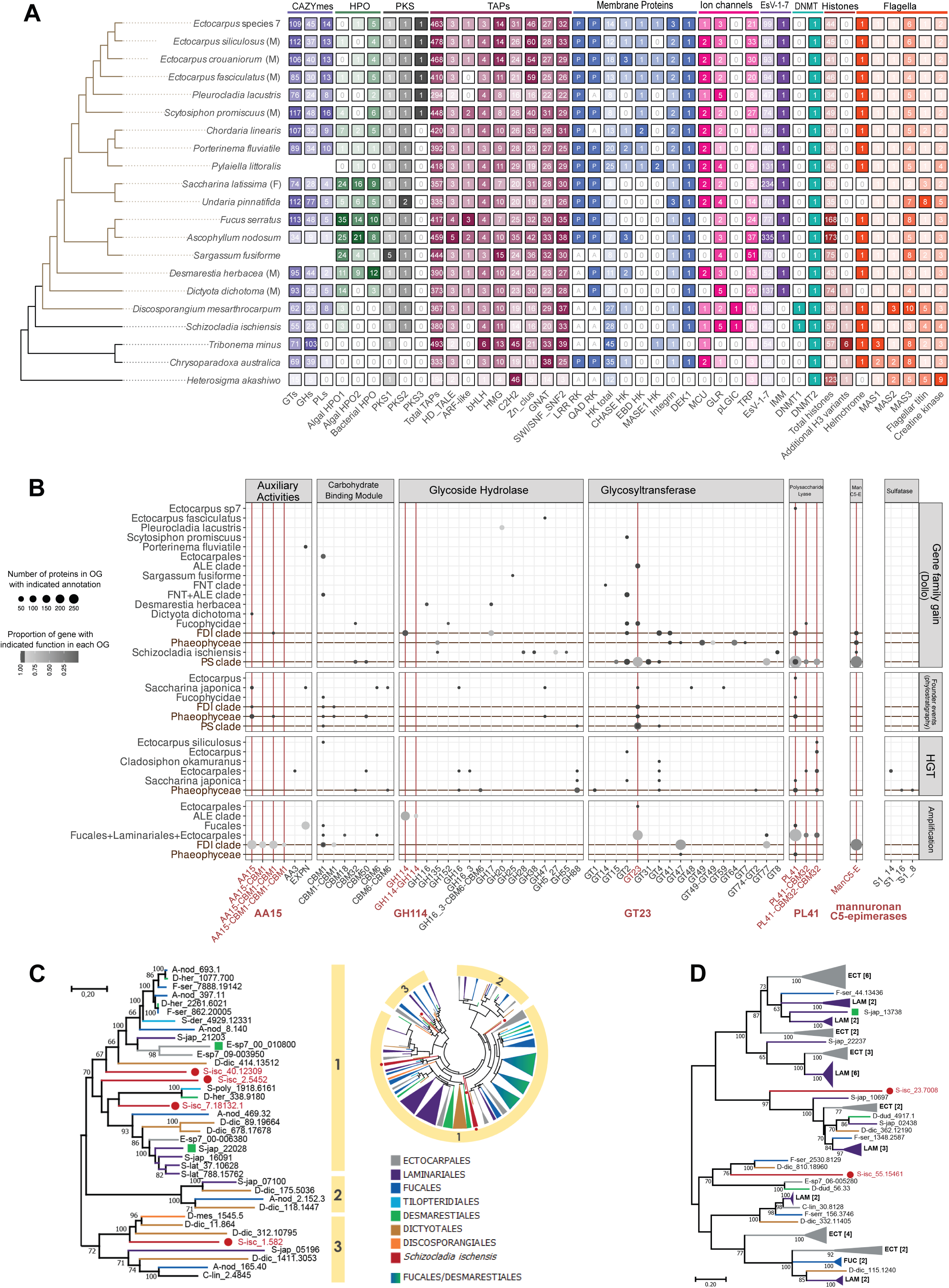
Gene family evolution during the emergence of the brown algal lineage and a focus on carbohydrate metabolism. (A) Variations in size for a broad range of key gene families in the brown algae and closely-related taxa. Numbers indicate the size of the gene family. Note that the *S. ischiensis* algal-type HPOs appear to be intermediate between classes I and II. Brown tree branches, Phaeophyceae. (B) Overview of information from the orthogroup Dollo analysis, the phylostratigraphy analysis, the horizontal gene transfer analysis and the gene family amplification analysis for a selection of cell-wall active protein (CWAP) families. Expert functional annotations been crossed with orthogroup (OG) composition, each dot represents a functional family / orthogroup couple. The size of the dot is proportional to the number of proteins annotated in the OG, and the colour represents the proportion of the functional annotation that falls into this OG. CWAP annotations are arranged by broad functional categories along the x-axis. Phylogenetic levels considered in the genome-wide analyses are indicated on the left. PS, Phaeophyceae and FDI clade, identified as gene innovation stages, are highlighted in brown. Functional categories with interesting evolutionary histories are highlighted in red. (C) Phylogenetic tree of mannuronan C5-epimerases (ManC5-E), a key enzyme in alginate biosynthesis. The inset circle represents a global ManC5-E phylogeny with three main clusters. The phylogeny shown on the left, with the three clusters indicated, is representative of the global view. (D) Phylogenetic tree of the polysaccharide lyase 41 (PL41) family, a key enzyme involved in alginate degradation in brown algae. The green squares indicate proteins that have been characterised biochemically. Brown algal sequences are colour-coded in relation to their taxonomy, as indicated in (C). Sequences belonging to the sister group Schizocladiaphyceae are shown in red and with a red circle. P, present; A, absent; CAZYmes, carbohydrate-active enzymes; HPO, vanadium haloperoxidase; PKS, type III polyketide synthase; TAPs, Transcription-associated proteins; EsV-1-7, EsV-1-7 domain proteins; DNMT, DNA methylase; GTs, glycosyltransferases; GHs, glycoside hydrolases; ARF, auxin response factor-related; bHLH, basic helix-loop-helix; HMG, high mobility group; Zn-clus, zinc cluster; C2H2, C2H2 zinc finger; GNAT, Gcn5-related N-acetyltransferase; SNF2, Sucrose nonfermenting 2; LRR, leucine-rich repeat; QAD, b-propeller domain; RK, membrane-localised receptor kinase; HK, histidine kinase; CHASE, cyclases/histidine kinases associated sensory extracellular domain; EBD, ethylene-binding-domain-like; MASE1, membrane-associated sensor1 domain; DEK1, defective kernal1; MCU, mitochondrial calcium uniporter; GLR, glutamate receptor; pLGIC, pentameric ligand-gated ion channel; TRP, transient receptor potential channel; IMM, IMMEDIATE UPRIGHT; H3, histone H3; MAS, mastigoneme proteins; AA, auxiliary activity; ECT, Ectocarpales; LAM, Laminariales; FUC, Fucales; DES, Desmarestiales.

Vanadium-dependent haloperoxidases (vHPOs) are a central component of brown algal halogen metabolism, which has been implicated in multiple biological processes including defence, adhesion, chemical signalling and the oxidative stress response. All three classes of brown algal vHPO (algal types I and II and bacterial-type^30–32)^ appear to have been acquired early during the emergence of the Phaeophyceae (Figures 3A, S8C, S8D, Tables S7, S8). Closely-related Stramenopile species do not possess any of these three types of haloperoxidase, with the exception of the sister taxon, *S. ischiensis*, which possesses three intermediate algal-type (i.e. equidistant phylogenetically from class I and class II algal-types) haloperoxidase genes (Figures 3A, S8C, S8D). Algal type I and II vHPO genes probably diverged from an intermediate-type ancestral gene similar to the *S. ischiensis* genes early during Phaeophyceae evolution. It is likely that the initial acquisitions of algal- and bacterial-type vHPOs represented independent events although the presence of probable vestiges of bacterial-type vHPO genes in *S. ischiensis* means that it is not possible to rule out acquisition of both types of vHPO through a single event. Clearly, however, gain of vHPO genes was an important event leading to the emergence of the emblematic halogen metabolism of brown algae early during their evolutionary history.

Gene gain may not, however, have been the proximal factor responsible for all the key metabolic innovations that occurred in the emerging brown algal lineage. Phlorotannins are characteristic brown algal polyphenolic compounds that occur in all Phaeophyceae species, with the exception of some members of the Sargassaceae. Phlorotannins are derived from phloroglucinol and brown algae possess three classes of type III polyketide synthase, two of which (PKS1 and PKS2) were acquired prior to the emergence of the Phaeophyceae and the third (PKS3) evolving much later within the Ectocarpales (Figures 3A, 4A, Table S9). Interestingly, PKS1 proteins from different brown algal species have been shown to have different activities leading to the production of distinct metabolites^33–35^, indicating that the acquisition of novel functions by this class of enzymes may have played an important role in the emergence of the brown algal capacity to produce phlorotannins. Moreover, many stramenopile PKS type III genes encode proteins with signal peptides or signal anchors (Figure S8E). For the brown algae, this feature is consistent with the cellular production site of phlorotannins and the observed transport of these compounds by physodes, secretory vesicles characteristic of brown algae^36^. Cross-linking of phlorotannins, embedded within other brown algal cell wall compounds such as alginates, has been demonstrated *in vitro* through the action of vHPOs^37–39^ and indirectly suggested by *in vivo* observations colocalising vHPOs with physode fusions at the cell periphery^40,41^. Consequently, vHPOs are good candidates for the enzymes that cross-link phlorotannins and other compounds, perhaps even for the formation of covalent bonds between phloroglucinol monomers and oligomers, which could occur via activation of aromatic rings through halogenation. These observations suggest that the acquisition of vHPOs by the common ancestor of brown algae and *S. ischiensis*, together perhaps with modifications of the existing PKS enzymes, triggered the emergence of new metabolic pathways leading to the production of the phlorotannin molecules characteristic of the Phaeophyceae lineage.

**Figure 4.**
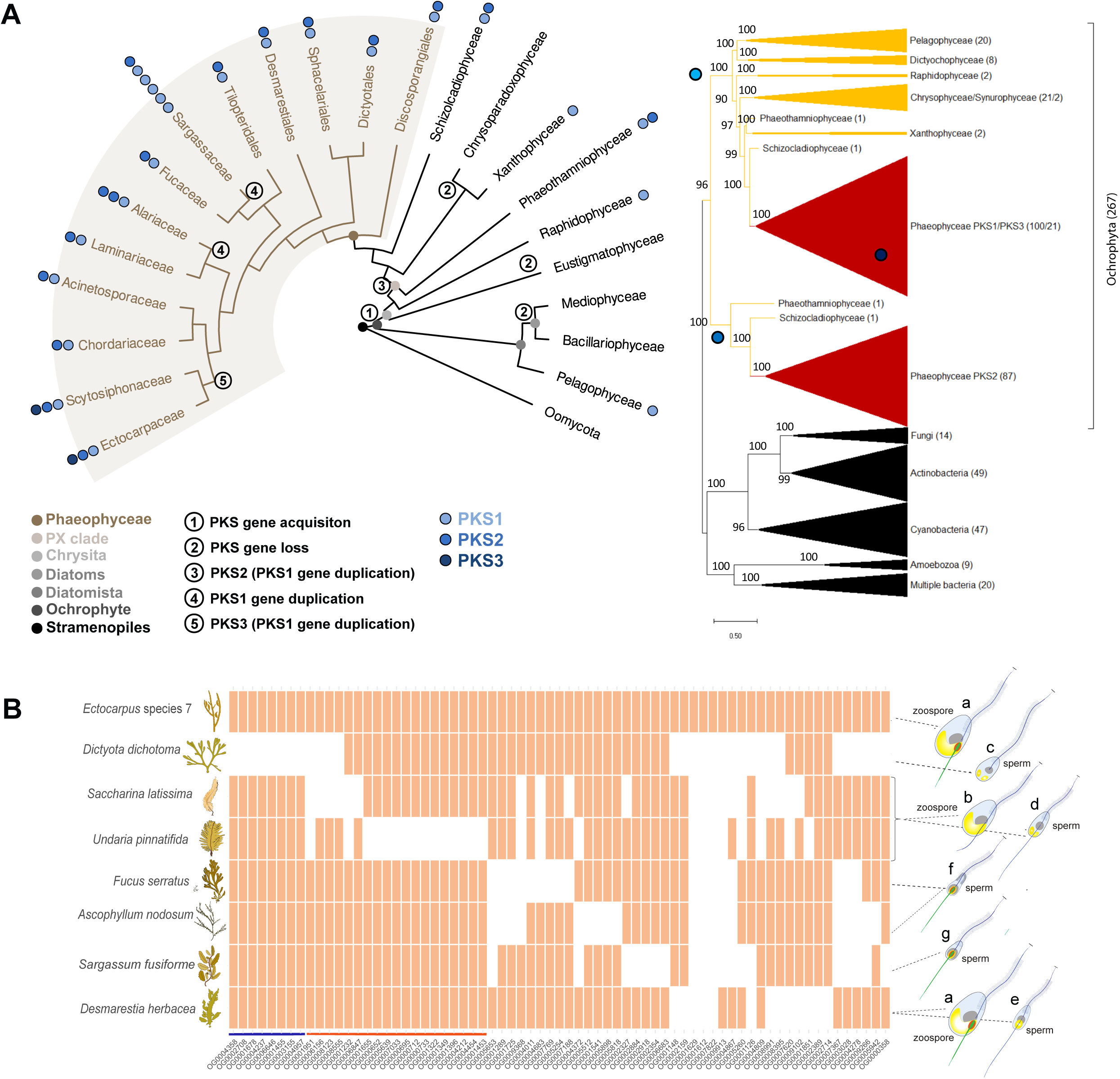
Evolution of key gene families during the emergence of the brown algal lineage. (A) Evolution of type III polyketide synthase (PKS) genes in the stramenopiles (left panel). Light blue, blue and dark blue circles correspond to the three classes of type III PKS (i.e. PKS1, PKS2 and PKS3), respectively. The number of circles indicates the number of gene copies and absence of a circle indicates absence of PKS genes. Coloured dots on the tree nodes indicate phylogenetic levels. Numbers indicate predicted evolutionary events. Right panel: condensed view of a phylogenetic reconstruction tree of stramenopile PKS III and closely-linked sequences. Red, orange and black branches correspond to brown algae, other stramenopiles and bacterial/eukaryotic outgroups, respectively. In brackets: number of sequences identified in each phylogenetic group. Branch bootstrap support is indicated. (B) Loss of orthogroups corresponding to flagellar proteome components^51^ in eight brown algal species from five orders. The drawings on the right indicate the cellular structures of zoids from the eight species. Grey, nucleus; yellow, chloroplast; blue, anterior flagella with mastigonemes; red, eyespot. The posterior flagellum is shown either in green to indicate the presence of green autofluorescence correlated with the presence of the eyespot or in blue in species without an eyespot. Note the loss of the eyespot in the Laminariales species *S. latissima* and *U. pinnatifida* and the loss of the entire posterior flagellum in *D. dichotoma*. Bars below the heatmap indicate gene losses associated with loss of just the eyespot (orange) or of the entire posterior flagellum (blue).

The acquisition of increased multicellular complexity and adaptation to new ecological niches during the early stages of brown algal evolution (i.e. during and immediately following the GOBE) is expected to have required modification and elaboration of signalling pathways. Membrane-localised signalling proteins (Figures 3A, S9) are of particular interest in this context not only as potential mediators of intercellular signalling in a multicellular organism but also because of potential interactions with the elaborate brown algal extracellular matrices (cell walls)^3,42^. A detailed analysis of the brown algal receptor kinase (RK) gene family, revealed that it actually includes two types of receptor, the previously reported leucine-rich repeat (LRR) RKs^11^ and a newly-discovered class of receptors with a beta-propeller extracellular domain (Figures 3A, S9A, Table S10). The phylogenetic distribution of RKs (Figures 3A, S9A) suggests that beta-propeller-type RKs may have originated before LRR RKs in the brown algae, although the presence of both beta-propeller and LRR RKs in *C. australica* (but none of the other closely-related outgroup species) raises questions about the timing of the emergence of these gene families. Note that oomycetes (distantly related stramenopiles) possess a phylogenetically distinct family of LRR RKs^11^ but not beta-propeller RKs. Diversification of receptor kinase families by the acquisition of different extracellular domains is therefore a characteristic that is so far unique to the three most complex multicellular lineages: animals, land plants and brown algae.

Major changes in epigenetic regulation also appear to have occurred during the emergence of the brown algae (see also supplemental results). *DNMT1* genes were identified in *Discosporangium mesarthrocarpum* and two closely-related outgroup species (*S. ischiensis* and *C. australica*) but not in other brown algal genomes, indicating that the common ancestor of brown algae probably possessed *DNMT1* but that this gene was lost after divergence of the Discosporangiales from other brown algal taxa (Figure 3A, Table S11). This is consistent with the reported absence of DNA methylation in the filamentous brown alga *Ectocarpus*^11^ and a very low level of DNA methylation in the kelp *Saccharina japonica*^43^ (which is thought to be mediated by DNMT2). Our analysis indicates that most brown algae either lack DNA methylation or exhibit very low levels of methylation and that this feature was acquired early during brown algal diversification.

### Impact of morphological, life cycle and reproductive diversification during the Mesozoic on brown algal genome evolution

A second major step in the evolutionary history of the Phaeophyceae was the rapid diversification of the major brown algal orders, which began after the origin of the FDI (Fucophycidae/Dictyotales/Ishigeales) clade, here estimated at 235.97 Ma (95% HPD: 158.88-312.48 Mya, an age that is broadly consistent with previous work^1^; Figures 1, S3). This diversification closely followed the Permian–Triassic mass extinction event (which dramatically impacted marine ecosystems in which red and green algae played dominant roles) and was facilitated by Triassic marine environments that favoured chlorophyll-c containing algae (e.g. high phosphate and low iron concentration), along with the appearance of new coastal niches created by Pangea rifting (Figure 1). This context would have facilitated the diversification of the brown algal lineage^44,45^, resulting in organisms that now exhibit a broad range of morphological complexity (ranging from filamentous to complex parenchymatous thalli), different types of life cycle and diverse reproductive strategies and metabolic capacities^3,6,29,46^ (Figures 1, S10A). The Phaeoexplorer dataset was analysed to identify genomic features associated with this diversification of phenotypic characteristics and to evaluate the impact on genome evolution and function.

We found indications that the diversification of life cycles, in some cases linked with the emergence of large, complex body architectures, impacted genome evolution through population genetic effects. Most brown algae have haploid-diploid life cycles involving alternation between sporophyte and gametophyte generations, the only exception being the Fucales, which have diploid life cycles. The theoretical advantages of different types of life cycle have been discussed in detail^47^ and one proposed advantage of a life cycle with a haploid phase is that this allows effective purifying counter-selection of deleterious alleles. When the brown algae with haploid-diploid life cycles were compared with species from the Fucales, increased rates of both synonymous and non-synonymous mutation rates were detected in the latter, consistent with the hypothesis that deleterious alleles are phenotypically masked in species where most genes function in a diploid context (Figure S10B). Comparison of non-synonymous substitution rates (dN) for genes in brown algae with different levels of morphological complexity, ranging from simple filamentous thalli though parenchymatous to morphologically complex, indicated significantly lower values of dN for filamentous species (Figure S10B). This observation suggests that the emergence of larger, more complex brown algae may have resulted in reduced effective population sizes and consequently weaker counter-selection of non-synonymous substitutions^24^.

For brown algae with haploid-diploid life cycles, the relative sizes of the two generations vary markedly across species with, for example, the sporophyte being much larger and more developmentally complex than the gametophyte in groups such as the kelps (Laminariales). To evaluate the effect of this diversity on life-cycle-related gene expression, generation-biased gene (GBG) expression was analysed in ten species whose haploid-diploid life cycles ranged from sporophyte dominance, through sporophyte-gametophyte isomorphy to gametophyte dominance. This analysis indicated, as expected^48^, that the number of GBGs was inversely correlated with the degree of intergenerational isomorphy (Figure S10C, Table S12). However, species with strongly heteromorphic life cycles had greater numbers of both sporophyte- and gametophyte-biased genes, indicating either that gene downregulation plays an important role in the establishment of heteromorphic generations or that there is also cryptic complexification of the more morphologically simple generation.

The diversification of the brown algae in terms of developmental complexity and life cycle structure was associated with modifications to reproductive systems, including, for example, partial or complete loss of flagella from female gametes in oogamous species and more subtle modifications such as loss of the eyespot in several kelps or of the entire posterior flagella in *D. dichotoma*^49,50^. Interestingly, these latter modifications are correlated with loss of the *HELMCHROME* gene, which is thought to be involved in light reception and zoid phototaxis^51^, from these species (Figures 3A, 4B). In addition, an analysis of the presence of genes for 70 high-confidence flagellar proteins^51^ across eight species with different flagellar characteristics identified proteins that correlate with presence or absence of the eyespot or of the posterior flagellum (Figure 4B, Table S13).

### Brown algal diversification and the emergence of marine forests was also associated with genomic changes affecting metabolic and signalling pathways

Forests of brown algae (i.e., Laminariales, Desmarestiales, Tilopteridales and Fucales^52^) are a key aspect of the modern marine biosphere. One of the pivotal innovations related to their emergence was a new developmental tissue, an intercalary meristem situated in the zone between the stipe and the lamina. The presence of this tissue is an ancestral state of the brown algal crown radiation (BACR) clade, and the current study indicates that the intercalary meristem was acquired as early as 190 Mya (Figure 1). This type of intercalary meristem would have facilitated the transition from annual to perennial life history and would, therefore, have been important for the establishment and maintenance of marine forests, particularly when upper parts of thalli are subjected to heavy grazing pressure^10^. Our results indicate that the Desmarestiales, Tilopteridales, and Fucales were all present by the early Cretaceous (Figure 1). Thus, it is possible that brown algal forests, at least at a small scale, provided both nutrients and shelter for the marine herbivorous animals that became common during the Cretaceous Period (e.g. algae-eating echinoids, sea turtles and euteleostean fish^53,54^).

While our estimates of kelp antiquity are earlier than those of Starko *et al*.^55^, they are consistent with their suggestion that Cenozoic cooling facilitated the geographic expansion of the kelp forest ecosystem. Indeed, many of the animals found today in kelp forest ecosystems originated toward the end of the Cretaceous Period, or later^56^. Currently, our understanding of Mesozoic marine noncalcified macroalgae on the basis of fossils^57–59^ is too poor to provide much guidance in this regard, but documentation by Kiel *et al*.^56^ of fossil holdfasts indicates that kelp forests were present by the late Paleogene Period (∼32 Mya). The highly complex, multi-layered and canopy-forming kelp forests of today, however, seem to have emerged only relatively recently, during the mid-Neogene, following the expansion of cooler water shelf environments^55,56^.

Comparative analysis of the Phaeoexplorer genome dataset identified a number of gene family expansions that potentially played important roles in the adaptation of the brown algae to their diverse niches and, more particularly, in the emergence of large, forest-forming species such as the kelps. For example, the ManC5-E family expanded markedly in the Laminariales and Fucales (Figure 3C), the two main orders that constitute extant phaeophycean forests. The capacity of ManC5-E to modify organ flexibility^3^ may therefore have been an important factor for large organisms coping with the harsh hydrodynamic conditions of coastal environments^60^. In addition, five different orthogroups containing proteins with the mechanosensor wall stress-responsive component (WSC) domain were identified as having increased in size during the diversification of the brown algal lineage (Table S3), indicating that metabolic innovations affecting cell walls may have been concomitant with a complexification of associated signalling pathways.

The haloperoxidase gene families, discussed above, expanded independently in several brown algal orders, again with expansions being particularly marked in the Fucales and the Laminariales (Figures 3A, S8C). In contrast, with some rare exceptions, the Ectocarpales, which are mostly small, filamentous algae, do not possess expanded vHPO gene families and there is evidence for multiple, independent gene losses resulting in loss of one or two of the three vHPO classes in many of the species in this order. In the Laminariales, the algal type I family are specialised for iodine rather than bromine^61^ and this may have been an innovation that occurred specifically within the Laminariales, resulting in a halogen metabolism with an additional layer of complexity. However, further analysis of the enzymatic activities of proteins from other orders will be necessary to determine if substrate specialisation was concomitant with this specific gene family expansion.

One of the proposed roles of halogenated molecules in brown algae is in biotic defence^4^ and, clearly, an effective defence system would have been an important prerequisite for the emergence of the large, perennial organisms that constitute marine forests. Additional immunity-related families^62^ that expanded during the diversification of the brown algae include five orthogroups that contain either GTPases with a central Ras of complex proteins/C-terminal of Roc RocCOR domain tandem (ROCO GTPases) or nucleotide-binding adaptor shared by apoptotic protease-activating factor 1, R proteins and CED-4 tetratricopeptide repeat (NB-ARC-TPR) genes (Table S3). In addition, analysis of gene family expansions indicated that modifications to additional biological processes such as photosynthesis, carbon metabolism and signalling may have been important (Figures S11, S12, Table S3, S14, S15, S16).

Finally, one of the most remarkable gene family amplifications detected in this study was for proteins containing the EsV-1-7 domain, a short, cysteine-rich motif that may represent a novel class of zinc finger^63^. EsV-1-7 domain proteins are completely absent from animal and land plant genomes and most stramenopiles either have just one member (oomycetes and eustigmatophytes) or entirely lack this gene family^63^. Analysis of the Phaeoexplorer data (Figure 3A, Table S17) indicated that the EsV-1-7 gene family started to expand in the common ancestor of the brown algae and the raphidophyte *H. akashiwo*, with 31-54 members in the non-Phaeophyceae taxa that shared this ancestor. Further expansion of the family then occurred in most brown algal orders, particularly in some members of the Laminariales (234 members in *Saccharina latissima*) and the Fucales (335 members in *Ascophyllum nodosum*), with the genes tending to be clustered in tandem arrays (Tables S3, S17). These observations are consistent with the previous description of a large EsV-1-7 domain family (95 genes) in *Ectocarpus* species 7^63^ and with recent observations by Nelson *et al*.^20^. One member of this family, IMMEDIATE UPRIGHT (IMM), has been shown to play a key role in the establishment of the elaborate basal filament system of *Ectocarpus* sporophytes^63^, suggesting that EsV-1-7 domain proteins may be novel developmental regulators in brown algae. Orthologues of the *IMM* gene were found in brown algal crown group taxa and in *Dictyota dichotoma* but not in *D. mesarthrocarpum* (Figure 3A, Table S17), indicating that this gene originated within the EsV-1-7 gene family as the first brown algal orders started to diverge. *IMM* is therefore conserved in most brown algal taxa but its function in species other than *Ectocarpus* remains to be determined.

### Recent evolutionary events within the genus *Ectocarpus*

The above analyses focused on deep-time evolutionary events related to the emergence of the Phaeophyceae and the later diversification of the brown algal orders during the Mesozoic. To complement these analyses an evaluation of relatively recent and ongoing evolutionary events in the brown algae was conducted by sequencing 22 new strains from the genus *Ectocarpus*, which originated about 19 Mya (Figure S13A).

A phylogenetic tree was constructed for 11 selected *Ectocarpus* species based on 261 high quality alignments of 1:1 orthologs (Figure 5A). The tree indicates substantial divergence between *E. fasciculatus* and two well-supported main clades, designated clade 1 and clade 2. Rates of synonymous and non-synonymous substitutions, based on alignments of genes in syntenic blocks between *Ectocarpus* species 7 and four other species, correlated with relative divergence times as estimated from the phylogenetic tree (Table S18), supporting the tree topology. Incongruencies between the species tree and trees for individual genes indicated introgression events and/or incomplete lineage sorting across the *Ectocarpus* genus. This conclusion was supported by phylogenetic tree comparisons, Bayesian hierarchical clustering, phylogenetic network reconstructions and PCoA analysis (Figures 5A, S13B, S13C). D-statistic analysis, specifically ABBA-BABA tests, detected incongruities among species quartets, indicating potential gene flow at various times during the evolution of the *Ectocarpus* genus. Evidence for gene flow was particularly strong for clade 2 and there was also evidence for marked exchanges between the two main clades (Figure 5B), suggesting that gene flow has not been limited to recently-diverged species pairs. These findings suggest a complex evolutionary history involving rapid divergence, hybridization and introgression among species within the *Ectocarpus* genus, with evidence for hybridisation occurring between 10.5 Mya (for clades 1 and 2) and 3.3 Mya (for *Ectocarpus* species 5 and 7) based on the fossil-calibrated tree (Figure S13A). A similar scenario has been reported for the genus *Drosophila*^64^, suggesting that recurrent hybridization and introgression among species may be a common feature associated with rapid species radiations. Major environmental changes such as the expansion of cold-water coastal areas following the green-house / cold-house Eocene–Oligocene transition (∼30 Mya^68^), and particularly the rapid climate destabilisation and temperature drop associated with the end of the mid-Miocene thermal maximum (∼15 Mya^68^), may have created many new opportunities for the rapid expansion and diversification of the *Ectocarpus* genus.

**Figure 5.**
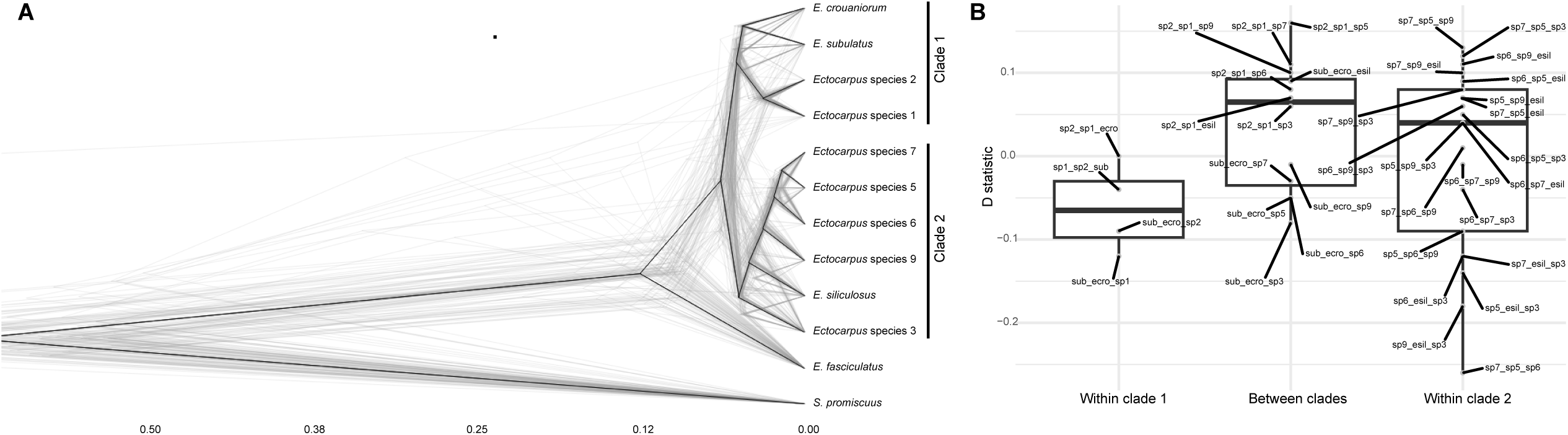
Evidence for gene flow within the genus *Ectocarpus*. (A) DensiTree visualisation of gene trees (grey lines) for 261 orthologues shared by 11 *Ectocarpus* species and the outgroup species *S. promiscuus,* together with the consensus species tree (black lines). The species tree is strongly supported, with all nodes displaying high posterior probabilities consistently reaching 0.99 or 1. (B) Box plot reporting the range of D-statistic (Patterson’s D) values between P2 and P3 species. Within-lineage comparisons (i.e. within clades 1 and 2) and between-lineage comparisons are distinguished on the x-axis. The annotation of each dot indicates species that were designated as P2 and P3. For this test, *Ectocarpus fasciculatus* was defined as the outgroup.

### Brown algal genomes contain large amounts of inserted viral sequences

A particularly striking result of this study was the identification of extensive amounts of integrated DNA sequence corresponding to large DNA viruses of the *Phaeovirus* family (Figure 6A, Table S19), which integrate into brown algal genomes as part of their lysogenic life cycles^69^. Analysis of 72 genomes in the Phaeoexplorer and associated public genome dataset identified a total of 792 viral regions (VRs) of *Nucleocytoviricota* (NCV) origin in 743 contigs, with a combined length of 32.3 Mbp. Individual VRs ranged in size from two to 705 kbp, but the majority (81.3 %) were between two and 50 kbp, whilst only 9% were longer than the expected minimum size (100 kbp) for an NCV genome. On average, VRs comprised 65% of their contig by length, and 72% of contigs with VRs were shorter than 100 kbp. Therefore, for most VRs, their true size and genomic context is unknown due to assembly gaps and short contig lengths. However, 40.8% of VRs had at least one flanking region providing direct evidence for insertion of the sequence in the algal genome (Table S19). Figure 6B shows three examples of long VRs. Most genes in VRs are monoexonic and transcriptionally silent, as previously observed for the 310 kbp VR in the *Ectocarpus* species 7 strain Ec32 genome^11^.

**Figure 6.**
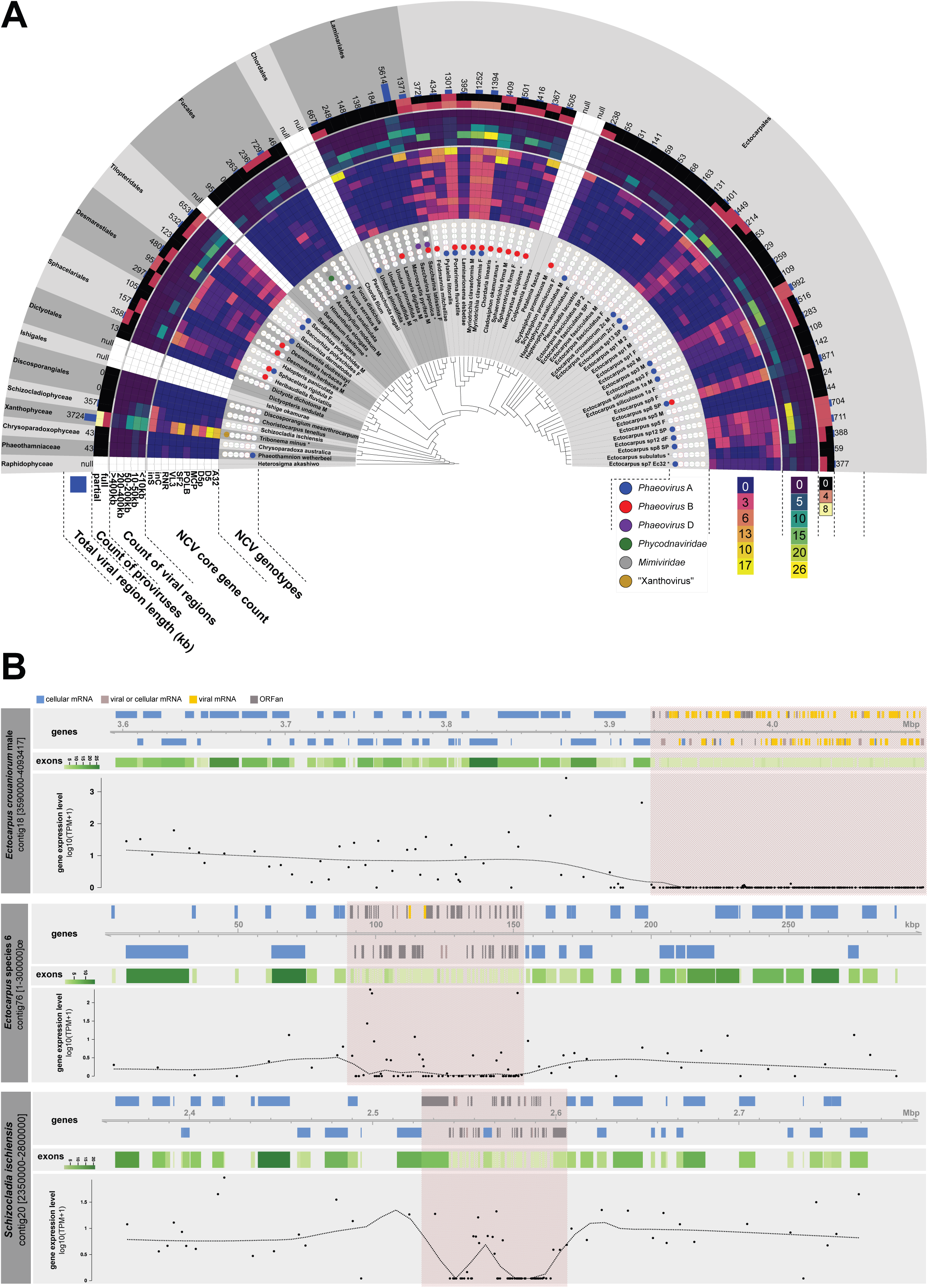
Inserted viral regions in brown algal genomes. (A) Annotated phylogeny summarising key statistics of the presence of nucleocytoviricota (NCV) sequences in the genomes of brown algae and closely-related taxa. Eight genomes that were sourced from public databases are labelled with an asterisk. Outer layers around tree are as follows: 1) NCV genotypes in each genome, 2) NCV core gene count indicates the number of copies of each viral core gene (A32, A32 packaging ATPase; D5/D5p, D5 helicase/primase; MCP, major capsid protein; POLB, DNA polymerase B; SF2, superfamily 2 helicase; VL3, very late transcription factor 3; RNR, ribonucleotide reductase; inC, integrase recombinase; inS, integrase resolvase), 3) Count of viral regions is the number of viral regions within each size range category as indicated, 4) Count of proviruses is the estimated number of complete or partial integrated viral genomes in a genome, 5) Total viral region length is the sum of the lengths in kbp of all viral regions within a genome. The outermost layer indicates the taxonomic class or order of the host clades. (B) Three examples of contigs containing large viral insertions (pink shading). Genes (coloured boxes) were classified as viral, cellular (i.e. cellular organism), known proteins of unclear origin (viral or cellular) or unknown (ORFan) based on comparisons with viral and cellular protein databases (see STAR Methods). Transcript abundances are shown with a locally estimated scatterplot smoothing (LOESS) plot. Exons, exons per gene.

On average, each of the 72 analysed genomes contained 469 kbp of VR (with a maximum of 5,614 kbp) and only two genomes contained no VRs (both from the Discosporangiales). There were a number of outlier genomes that contained more than 1 Mbp of VRs (*T. minus*, *S. latissima*, *S. japonica*, *P. fluviatile* and *Myriotrichia clavaeformis* male and female). The presence of key NCV marker genes was used to assess the completeness of inserted viral genomes. At least one partial provirus (a VR possessing several key NCV marker genes) was present in 39 genomes, 29 of which had at least one full provirus with a complete set of seven key NCV marker genes (Figure 6A, Table S19). In addition to the previously known infections in Ectocarpales^69^ and Laminariales^70^, integrated NCV proviruses were found in all Phaeophyceae orders screened, except the Discosporangiales and Dictyotales, and were also detected in *T. minus* (Xanthophyceae). Moreover, NCV marker gene composition indicated that multiple integrated proviruses were present in 16 genomes from multiple Phaeophyceae orders (Ectocarpales, Desmarestiales, Sphacelariales, Tilopteridales, Laminariales), and the Xanthophyceae (Figure 6A, Table S19). Phylogenetic analysis of the major capsid protein (MCP) and DNA polymerase genes indicated that the majority of the integrated NCVs belonged to the genus *Phaeovirus*, the sole viral group known to infect brown algae (Figure S14A,B). However, this analysis also revealed integrated sequences corresponding to other viral groups. Viral sequences in *T. minus* belonged to a putative novel genus closely related to *Phaeovirus*, for which we propose the name *Xanthovirus*. Finally, mimiviridae-related VRs were identified in *S. latissima* and *Pelvetia canaliculata*, but since they are partial proviruses and do not appear to possess integrase genes, they may have originated from ancient endogenization events, similar to those described in chlorophytes^71^.

The identification of integrated NCVs across almost all brown algal orders and in closely related outgroup taxa suggests that the lysogenic life cycle strategy of phaeoviruses is ancient and that giant viral genomes have been integrating into the genomes of brown algae throughout the latters’ evolutionary history. This conclusion was supported by the phylostratigraphic analysis, which detected the appearance of many novel virus-related genes dating back to the origin of the Phaeophyceae (Figure S5A). Marked differences were detected in total VR size and NCV marker gene presence across the brown algal genome set, and large differences were even detected between strains from the same genus (between 24 and 992 kbp of VR in different *Ectocarpus* spp. for example; Figure 6A, Table S19). These differences indicate dynamic changes in VR content over evolutionary time, presumably due, at least in part, to differences in rates of viral genome integration, a process that can involve multiple, separate insertion events^72^, and rates of VR loss due to meiotic segregation^73^. In addition, the abundant presence of partial proviruses and NCV fragments in brown algal genomes indicates that inserted VRs can degenerate and fragment, probably also leading to VR loss over time. The identification of large-scale viral genome insertion events over such a long timescale (at least 450 My^1^) suggests that NCVs may have had a major impact on the evolution of brown algal genomes throughout the emergence of the lineage, making this a truly unique model system to investigate the impact of genomic exchanges on genome evolution.

The widespread presence of large quantities of viral genes in brown algal genomes creates a favourable situation for recruitment of this genetic information by the algal host via HGT (provided the acquired genes confer a selective advantage^74^) but clear evidence of this type of HGT event can be difficult to obtain. However, phylogenetic evidence indicates that several *Ectocarpus* species 7 histidine kinases were derived by HGT from viral insertions^75^ and analysis of the Phaeoexplorer genomes supported this hypothesis. Histidine kinases (HKs) are widespread in the stramenopiles but several classes of membrane-localised HK were either only found in brown algae (CHASE domain HKs and HKs with an extracellular domain resembling an ethylene binding motif^75^) or only in brown algae and closely-related taxa (MASE1 domain HKs^75^) and appear to be absent from other stramenopile lineages (Figures 3A and S9A, Table S20). These classes of HK all exhibit a patchy pattern of distribution across the brown algae and are often monoexonic suggesting possible multiple acquisitions from viruses via HGT following integration of viral genomes into algal genomes (Figures 3A, S9A, S14C). Phylogenetic analysis provided further support for a HGT origin for these classes of HK (Figure S14C).

## DISCUSSION

The genome resource presented and analysed in this study provides an unprecedented overview of genomic diversity across the full scope of the brown algal lineage. Comparative analysis of these genomes has allowed insights into brown algal genome evolution across the entire evolutionary history of the group, ranging from emergence of the lineage during the GOBE through diversification of the brown algal orders during the Mesozoic and up until relatively recent evolutionary events at the genus level during the Miocene.

These analyses identified a period of marked genome evolution concomitant with the emergence of the brown algal lineage that correlates with an increase in multicellular complexity possibly driven, at least in part, by increases in atmospheric oxygen and herbivory. These early genome evolution events included enhanced levels of gene gain (including via HGT) and the formation of genes with novel domain combinations. These genomic changes were important in equipping the brown algal ancestor with key components of several metabolic pathways, notably cell wall polysaccharide, phlorotannin and halogen metabolisms, that were essential for their colonisation of intertidal and subtidal environments. The capacity to synthesise flexible and resilient alginate-based cell walls allows these organisms to resist the hydrodynamic forces of wave action^60^, whereas phlorotannins and halogen derivatives are thought to play important roles in defence^76^. There is also evidence that cell wall cross-linking by phlorotannins may be important for strong adhesion to substrata, another important characteristic in the dynamic intertidal and subtidal coastal environments^77^. The capacity to adhere strongly and resist both biotic and abiotic stress factors would prove essential for the success of large, sedentary multicellular organisms in these intertidal niches over evolutionary time. Interestingly, the early genome evolution events described above also set the stage for later genome evolution events in the sense that the gene families that have amplified in the brown algae are significantly enriched in genes that were gained during the emergence of the lineage.

The period of increased gene gain during the emergence of the brown algae was followed by a period of overall gene loss that extended up until the present day (Figure S4B). Interestingly, similar periods of ancestral gene gain followed by gene loss have also been observed for both the animal and land plant lineages^78^, indicating that this may be a common feature of multicellular eukaryotic lineages.

About 220 MY after the emergence of the brown algae, the aftermath of the Permian–Triassic mass extinction event and the initiation of Pangea rifting appear to have created favourable environments for rapid diversification of the main brown algal orders^44,45^, resulting in the emergence of a diversity of developmental, life cycle and reproductive strategies, with correlated effects on genome evolution. In some orders, such as the Laminariales and Fucales, certain characteristics acquired during this period of diversification are predicted to have been important prerequisites for the emergence of marine forests. These characteristics include developmental features such as an intercalary meristem but also modification of metabolic, defence and developmental processes, novelties which seem to have arisen, as indicated by the present study, through order-specific amplifications of key gene families.

Analysis of the genomes of multiple *Ectocarpus* species demonstrated that genomic modifications, including gene gain and gene loss have continued to occur up until the present time and indicated that these modifications can potentially be transmitted between species as a result of gene flow occurring within a genus due to incomplete reproductive boundaries and introgression.

Finally, one of the most surprising observations was that brown algal genomes contain many inserted viral sequences corresponding to large DNA viruses of the *Phaeovirus* family. Inserted viral sequences are widespread in eukaryotic genomes^79,80^ and insertions corresponding to nucleocytoplasmic large DNA viruses have been found in green algal genomes^71,81^ but the brown algal *Phaeovirus* VRs are remarkable because they are nearly ubiquitous in this lineage (being present in 67 of 69 brown algal genomes analysed) and because individual genomes can contain several phylogenetically diverse *Phaeovirus* insertions and insertions of a broad range of different sizes. The near ubiquitous occurrence of these elements may be attributed to the capacity of phaeoviruses to insert into their hosts’ genomes as part of their life cycle, whereas the broad range of brown algal species that harbour these elements indicates that *Phaeovirus* insertions have probably been occurring since the emergence of this lineage.

The above observations illustrate how the Phaeoexplorer genome dataset, along with the various analyses carried out in this study, can be used to link the gene content of brown algal genomes to biological processes and characteristics that have played key roles during the evolution of this lineage. The establishment of this genome resource represents an important step forward for a key lineage that has remained poorly characterised at the genome level. The Phaeoexplorer dataset not only provides good quality genome assemblies for many, previously uncharacterised brown algal species but also represents a tool to explore genome function via comparative genomics approaches, adding an important evolutionary dimension to efforts to understand gene function in this lineage. The identification and analysis of key metabolic and signalling genes implicated in a broad range of brown algal biological functions (growth and development, adhesion, reproduction and resilience to abiotic and biotic stresses) represents an important resource for future research programs aimed at optimising brown seaweed production in a mariculture context or at preserving and protecting natural seaweed populations in the context of climate change. Both of these approaches could potentially contribute to mitigation of the effects of climate change via multiple positive effects in terms of carbon capture, ecosystem services and by the promotion of highly sustainable cultivation practises.

To facilitate future use of this genome dataset, the annotated genomes have been made available through a website portal (https://phaeoexplorer.sb-roscoff.fr), which provides multiple additional resources including genome browsers, BLAST interfaces, transcriptomic data and metabolic networks. The existing genome dataset provides very good coverage of the phylogenetic diversity of the Phaeophyceae and reasonably complete gene catalogues for each species but future work is needed to improve further the quality of the genome assemblies described here and to add genomes for additional species, particularly members of the minor brown algal orders that are not represented in the dataset. The large proportion of genes with no predicted function in brown algal genomes is also a limitation that needs to be addressed. The recent development of CRISPR-Cas9 methodology for brown algae^82,83^, together with the other tools and resources currently available for the model brown alga *Ectocarpus*^84^, provide the means to deploy the functional genomics approaches necessary to address this question.

## ACKNOWLEDGMENTS

This work was supported by the France Génomique National infrastructure project Phaeoexplorer (ANR-10-INBS-09), the European Research Council project Sexsea (638240), the Investissements d’Avenir project Idealg (ANR-10-BTBR-04-01), the European BG-01 BlueGrowth H2020 project Genialg (727892), Laoshan Laboratory grants (LSKJ202203801, LSKJ202203204), the Taishan Scholars Program and Talent Projects of Distinguished Scientific Scholars in Agriculture, the CNRS international research network DABMA (00022), the ANR projects Epicycle (ANR-19-CE20-0028-01), BrownSugar (ANR-20-CE44-0011), HaloGene (ANR-22-CE20-0025), Seabioz (ANR-20-CE43-0013) and BrownLincs (ANR-23-CE20-0048-01), the National Research Foundation of Korea (2022R1A2B5B03002312, 2022R1A5A1031361) granted to H.S.Y., the projects Connect Talent EpiAlg Région Pays de la Loire-Nantes Métropole and Etoiles Montantes M-EpiCC Région Pays de la Loire, the MITI-funded project Algometabionte, Dr. Karl Feldbausch-Stiftung, the CNRS and Sorbonne University. We are grateful to the Roscoff Bioinformatics platform ABiMS (http://abims.sb-roscoff.fr), which is part of the Institut Français de Bioinformatique (ANR-11-INBS-0013) and BioGenouest network, for providing both help and computing and storage resources.

## AUTHOR CONTRIBUTIONS

Software: L.B.G., R.D., A.L.B., X.L., D.N. and E.C.; Formal analysis: F.D., O.G., L.D., D.M., T.M., D.S., X.F., L.Ma., N.Te., J.B.R., R.P., L.R., S.W.C., J.J., K.U., K.B., C.Du., P.R., A.L., A.E.M., M.L., A.K., P.H.G., C.V., S.S.A., S.A., K.A., Y.B., T.Ba., A.Be., S.B., A.Bo., A.Cor., H.C.C., A.D., E.Di., S.D., E.Dr., J.G., L.G., A.G., M.L.G., L.H., B.H., A.J., E.K., A.H.K., C.B., L.L., R.L., S.T.L., P.J.L., E.M., S.M., G.M., C.N., S.A.R., E.R., D.Sch., A.S., L.T., T.T., K.V., H.V., G.W., H.K., A.F.P., H.S.Y., C.H., N.Y., E.B., M.V., G.V.M., E.C., S.M.C., J.M.A. and J.M.C; Investigation: C.C., S.H., Z.N., N.Ta., A.Cou., B.N., W.B., E.De., C.J., L.Me., S.R. and D.Sco.; Resources: O.G., A.Cou., L.B.G., T.Br., R.A.C., C.De., S.F., W.J.H., G.H., K.K., A.L.B., K.M., C.M., N.P., P.P., S.R., D.Sco., H.V., F.W., H.K., A.F.P., M.V., E.C. and J.M.A.; Data Curation: O.G., C.C., A.Cou., L.B.G., J.M.A. and J.M.C.; Writing-Original Draft: F.D., O.G., J.M.A. and J.M.C.; Writing-Review & Editing: all authors; Visualization: F.D., O.G., L.B.G., L.D., D.M., T.M., D.S., X.F., L.Ma., J.B.R., R.P., L.R., K.U., K.B., P.R., A.L., M.L., P.H.G., C.V., A.D., A.L.B., H.K., E.C. and J.M.C.; Supervision: O.G., N.Te., M.L., R.A.C., H.C.C., O.D.C., S.D., G.H., A.J., C.B., E.P., P.L., S.A.R., A.S., L.T., H.V., G.W., H.K., H.S.Y., C.H., N.Y., E.B., M.V., G.V.M., E.C., S.M.C., J.M.A. and J.M.C.; Project administration: F.D., E.B., M.V., G.V.M., E.C., S.M.C., P.W., J.M.A. and J.M.C.; Funding acquisition: H.C.C., C.B., P.P., C.H., N.Y., S.M.C. and J.M.C.; Conceptualization: J.M.C.

## DECLARATION OF INTERESTS

The authors declare no competing interests.

## Supplementary figure legends

Figure S1. Taxonomic diversity and assembly quality of the Phaeoexplorer genomes, related to Figure 1.

(A) Taxonomic distribution and assembly quality (contig N50) of the Phaeoexplorer genome dataset (blue) and previously published brown algal genomes (brown). “Reference” quality Phaeoexplorer genomes are circled in black.

(B) Number of genes annotated in each genome (upper panel). BUSCO scores for the predicted proteome of each genome (middle panel). Correlation of CDS lengths for each species with the corresponding sequences from the *Ectocarpus* species 7 reference genome (lower panel). Previously published genomes are indicated in grey. longreads, genomes assembled using long reads.

F, female; M, male.

Figure S2. Geographic localisation of species and strains, related to Figure 1.

(A) World map indicating the positions of the sampling sites for the strains sequenced in this study.

(B) North-south distributions of the species analysed in this study. The data shown were obtained from the Global Biodiversity Information Facility (gbif.org). Asterisks indicate the sampling latitude for the sequenced strains of each species.

Figure S3. Fossil-calibrated maximum-likelihood phylogenetic tree of the brown algal and closely-related taxa analysed in this study, related to Figure 1.

Bayesian divergence time estimation using 32 nuclear protein sequences, together with two fossil calibrations and a calibration for the root based on Choi *et al*.^1^ (numbered blue circles). Grey bars on the nodes show 95% highest density region (HPD) intervals of the node ages. The Great Ordovician Biodiversification Event (GOBE) is shown as an orange box, the Permian-Triassic mass extinction as a purple band and Pangea rifting as a blue box.

Figure S4. Dollo-logic-based analysis of gene family gain and loss during the emergence of the brown algae, related to Figure 2.

(A) Cladogram indicating orthogroup (OG) gain and loss during the emergence of the Phaeophyceae based on Dollo parsimony analysis. Taxonomic classes, orders and families are indicated in brown (brown algae) or grey (outgroup species) on the right. The nodes of the tree (n0-n22) are numbered in red, and listed on the left with the corresponding name, if one exists. The number of OGs predicted to be present at each node is indicated by the circles and the branches are coloured according to overall gene family gain.

(B) Heat map of COG functional categories associated with OGs gained at each node of the cladogram.

Figure S5. Genome-wide analyses of gene family evolution, related to Figure 2.

(A) Functional and structural features of founder genes from three taxonomic levels. FDI, Fucophycideae/Dictyotales/Ishigeales; PS, Phaeophyceae plus Schizocladiophyceae; PX, Phaeophyceae plus Xanthophyceae.

(B) Inference of HGT origins from 74 species based on monophyletic most similar homologue (MMSH) analysis. The tree on the left, which is derived from the NCBI taxonomy tree, indicates the source taxa for HGT-derived genes. The tree at the bottom of the figure, which was constructed using single-copy genes, indicates the species that have received genes by HGT. The middle part of the figure indicates the number of HGT genes transferred from each source to the receiver genomes. The panel on the right illustrates the number of HGTs from each phylum. The legend in the lower left corner provides a reference for the circle size, which corresponds to 10, 100 or 300 HGT gene counts.

(C) Composite gene analysis. Phylogenetic distribution of fused (blue), split (green) and non-remodelled (grey) gene family originations across the evolution of brown algae (middle panel). Pie charts on each branch of the phylogeny indicate the relative contribution of gene fusion and fission to the overall emergence of novel gene families, quantified by the area of the circle. Brown algal species are indicated in brown and other stramenopiles in black. Note that only the topology of the species tree is displayed here, without specific branch lengths. Right panel: barplot indicating the percentage of gene families retained in extant genomes among all gene families that emerged during the evolution of the species set. Left panel: barplot representing the distribution of gene families in COG functional categories for functionally annotated fused, split, and non-remodelled orthologous groups. The functional annotation assigned to an orthogroup corresponded to the most frequent functional category annotated for the members of each orthogroup. Asterisks next to the bars indicate statistically significant differences between remodelled and non-remodelled gene families (*p*-value <0.05, two-sided Chi^2^ test with Yates correction).

(D) Left panel: gene families (orthogroups) significantly amplified (binomial test) in the brown algae compared to outgroup taxa. Orthogroups amplified in specific groups of species are indicated by green rectangles. Right panel: pie charts representing the proportions of manually-determined functional categories for each group of amplified gene families highlighted in the left panel.

(E) Plot showing the timing (phylogenetic clade) of the amplification of gene families (Y axis) and the timing of the appearance (gain) of the amplified gene family (X axis, based on the Dollo analysis of orthogroups). Nodes (e.g. n2) are as indicated in Figure S4.

Figure S6. Structural features of the predicted genes in the Phaeoexplorer genomes, related to Figure 1.

(A) Genome and gene statistics for the 21 reference genomes (Figure S1A and Table S1). Violin plots display size distributions. Cross or triangle, mean; diamond or square, median. For intergenic regions, half violins on the left correspond to intergenic distances between adjacent genes on opposite strands, and half violins on the right to intergenic distances between adjacent genes on the same strand.

(B) Intron acquisition during the emergence of the brown algal lineage (left panel). The analysis examined 949 introns in 235 conserved orthologues. Colours correspond to the species distributions of introns. Hatched blue indicates introns that are shared with at least two outgroups (ancestral introns). The right panel shows an example of an intron conservation profile for the orthogroup OG0004854. Colour code and species numbering as for the left panel. F, female; M, male.

Figure S7. Genome-scale metabolic network analyses, related to Figure 2.

(A-D). Multidimensional-scaling (MDS) plots of GSMNs reconstructed with AuCoMe using the presence and absence of biochemical reactions in these networks to compute the distance matrix (Jaccard index) used by the MDS.

(A) MDS computed using the draft metabolic networks created by the first step of AuCoMe with only the annotation from the genomes on all species from the Phaeoexplorer dataset.

(B) MDS computed using metabolic networks created after the last step of AuCoMe, i.e. after reaction propagation using orthologous genes and structural verification on genome sequence, on all genomes from the Phaeoexplorer dataset.

(C) MDS computed on draft metabolic networks from a subset of high-quality assemblies.

(D) MDS computed after reaction propagation on a subset of high-quality assemblies. Dim., dimension.

(E-G). Overview of losses and gains of metabolic components for different sets of brown algal species. Each row corresponds to a species and columns correspond to sets of reactions. For each row, there is a coloured block if the species contains the reactions present in this block and shared with the other species that also have this block in the same column. Numbers at the top, along the bottom and to the right indicate the number of species that possess each reaction, the number of reactions with this profile and the number of reactions per species, respectively.

(A) (E) All brown algae and stramenopile outgroups.

(B) (F) Selected reference quality genome assemblies.

(C) (G) Comparison of freshwater and marine species with reference quality genomes.

Figure S8. Analysis of metabolism gene families, related to Figure 3.

(A) Counts of numbers of genes predicted to encode glycosyltransferases (GTs), glycoside hydrolases (GHs), polysaccharide lyases (PLs) and all CAZymes (GTs, GHs, PLs, Carbohydrate Esterases CEs, Auxiliary Activities AAs, Carbohydrate Binding Modules CBMs), showing numbers for both full-length proteins (dark colours) and fragments (light colours). The data is averaged by order with the number of species analysed per order in brackets.

(B) Number of genes for selected CAZyme families in brown algae. Only CAZyme families with at least three members per genome on average, are shown. The species analysed are the same as in (A). Counts include full-length proteins and fragments.

(C) vHPO genes identified in the 21 reference genomes based on sequence homology and active site conservation. vHPO genes are indicated by a green cross or a number and absence by a red cross.

(D) Maximum likelihood phylogenetic trees for 259 algal-type vHPOs (left) and for bacterial-type vHPOs (right). Algal-type I vHPOs are coloured in blue and algal-type II vHPOs in violet. The clades that have structurally or biochemically characterised enzymes are highlighted in red for vBPOs and in yellow for vIPOs. Strongly supported representative branches have been collapsed. The clade names correspond either to taxa or to individual gene names. For the bacterial vHPOs, the FastTree reconstruction tool with 1000 bootstraps was drawn as a circular representation taking the gammaproteobacterial group as the starting point to arbitrary root the tree. Brown algal branches are coloured in brown. Green dots indicate bootstrap values of between 0.7 and 1.0 (1000 replicates).

(E) PKS III domain structures indicating amino-(IPR001099; green) and carboxy-terminal (IPR012328; brown) chalcone/stilbene synthase domains and amino-terminal signal peptide (SP; violet), signal anchor (SA; light blue), SP/SA hybrid (pink) or all three (SP, SA or SP/SA; black). The number of proteins in each group is indicated at the carboxy end of the protein.

Figure S9. Evolution of membrane-localised signalling proteins in the brown algae, related to Figure 3.

(A) Presence or absence of different membrane-localised signalling proteins in brown algae and other stramenopiles. (B) Diverse domain structures of stramenopile FG-GAP domain proteins showing possible evolutionary relationships. LRR, leucine-rich repeat; QAD, b-propeller domain; EBD, ethylene-binding domain-like; CHASE, cyclases/histidine kinases associated sensory extracellular domain; MASE1, membrane-associated sensor1 domain; HK, histidine kinase; FG-GAP, phenylalanyl-glycyl– glycyl-alanyl-prolyl domain; IPT, Ig-like plexins transcription factors domain; ITGA, α-integrin domain; Cad, cadherin domain; DEK1, defective kernal1; FAS, fasciculin; SP, signal peptide; TM, transmembrane domain.

Figure S10. Evolution associated with the diversification of biological traits across the brown algae, related to Figure 1.

(A) Ancestral state reconstructions for number of cell types and for morphological complexity in the diploid or haploid phase of the life cycle. Left, number of different cell types in the diploid phase (DP). Centre, morphological complexity in the diploid phase. Right, morphological complexity in the haploid phase (HP). The colours indicate the different states and the pie charts the likelihood of these states at each node.

(B) Rates of gene evolution in relation to life cycle structure and developmental complexity. Left, violin plots showing the distribution of omega (dN/dS), rates of non-synonymous substitution (dN) and rates of synonymous substitution (dS) for brown algae with haplodiplontic or diplontic life cycles. Right, violin plots showing the distribution of rates of non-synonymous substitution (dN) for brown algal species with filamentous, simple parenchymatous or elaborate parenchymatous thalli. The *p*-values are for pairwise (gene-by-gene) Wilcoxon tests and the percentage of genes that exhibited the same patterns of differences in dN/dS, dN or dS values as the median values are indicated. Significant differences are indicated by an asterisk.

(C) Generation-biased gene expression in relation to life cycle dimorphism. Pie diagrams indicating the proportions of sporophyte-specific (dark red), sporophyte-biased (light red), gametophyte-specific (dark blue), gametophyte-biased (light blue) and unbiased (grey) genes in species with different haploid-diploid life cycles ranging from sporophyte-dominant through isomorphic to gametophyte-dominant. sp., species.

Figure S11. Amplification of transcription factor gene families, related to Figure 3.

(A) Ancestral state reconstruction of the sizes of three TAP families: left, C2h2 zinc finger, centre, three-amino-acid loop extension homeodomain (HD_TALE) and, right, high mobility group (HMG). The colour gradient indicates the predicted number of genes in each family across the tree.

(B) Alignment of the homeodomain regions of three-amino-acid loop extension homeodomain transcription factors (TALE HD TFs). Sequences in clusters 1 and 3 are orthologous to the ORO and SAM proteins of *Ectocarpus* species 7, respectively. Cluster 2 corresponds to sequences similar to the third TALE HD TF gene of *Ectocarpus* species 7. Cluster 4 contains the sequences from closely-related outgroup taxa that could not be clearly assigned orthology to any of the *Ectocarpus* species 7 proteins.

Figure S12. Evolution of LHCX genes, related to Figure 3.

(A) LHCX cluster genomic context in ten brown algae and *Schizocladia ischiensis*. Text above and below the line indicates the chromosome (chr) or contig (ctg) and the gene number, respectively.

(B) Maximum likelihood (RAxML) phylogenetic tree of all LHCX and selected FCP protein sequences from the algal species in panel (A). Sequences are labelled with the first letters of the genus and species name, contig and gene number. Clustered LHCX genes are additionally numbered as in panel (A). Three groups of conserved LHCX paralogs encoded by unclustered, unique genes are highlighted by blue background.

(C) Examples of LHCX gene clusters (LHCX genes in red) in four brown algal species where the LHCX genes are located on opposite strands of the chromosome and overlap, either partially, with some of their exons being located in the intron of the gene on the opposite strand (*C. linearis, S. latissima*), or completely, with the gene being located entirely within the intron of the gene on the opposite strand (*E. crouaniorum, S. promiscuus*).

Figure S13. Microevolution of genomes within the genus *Ectocarpus,* related to Figure 5.

(A) Fossil-calibrated phylogenetic tree for 11 *Ectocarpus* species. Extracted from the fossil-calibrated tree shown in Figure S3. Numbered grey bars on the nodes show 95% HPD intervals of the node ages.

(B) Unrooted parsimony splits network of 11 *Ectocarpus* species. The tree was generated based on 261 concatenated orthologous genes using the ParsimonySplits method in Splitstree^85^. ParsimonySplits networks accommodate phylogenetic incongruities in the data by incorporating alternative branch splits. Conflicting splits are displayed as box-like structures.

(C) Principal Correspondence Analysis (PCoA) of distance branching patterns (Robinson-Foulds metric) emphasizing the node for *Ectocarpus* species 5, species 6 and species 7 in the 261 gene trees. The plot displays the first two principal axes, capturing the major patterns of variation in tree topology at this node among the gene trees. Each dot represents a single gene tree, with the distance between points corresponding to the dissimilarity in their branching patterns. Dot colour indicates the cluster membership of gene trees based on the clustering analysis.

Figure S14. Phylogenetic trees of viral and putatively virus-derived genes, related to figure 6.

(A) and (B) *Nucleocytoviricota* major capsid protein and DNA polymerase phylogenies, respectively. The trees were both generated from aligned amino acid sequences with the Q.yeast+F+R6 model and 1000 bootstraps in IQ-TREE. Subgroup labels refer to *Phaeovirus* genotypes. All sequences identified by this study are in bold, and highlighted in yellow if they clustered outside the *Phaeovirus*. Phy., *Phycodnaviridae*.

(B) Phylogenetic trees for three classes of histidine kinase. Membrane-associated sensor1 domain (MASE1) class: brown algal genes are more closely related to viral than to closely-related outgroup species genes, EsuBft1789.4 is located within a viral clade. Ethylene-binding-domain-like (EBD) class: viral-related clade limited to Ectocarpales. Cyclases/histidine kinases associated sensory extracellular domain (CHASE) class: complex pattern suggesting possible multiple HGTs. Genes are classed as algal (green or light green background label, strong or weak prediction) or inserted viral sequences (blue or light blue background label, strong or weak prediction) based on exon number, expression level and genomic context (neighbouring monoexonic or multiexonic genes). See Table S20 for the gene name abbreviations. Branch colours signify brown algae (green), closely-related outgroup taxa (violet) or EsV-1 genes (blue). TPM, maximum TPM; asterisks, gene located in an identified VR; nd, not determined.

Figure S15. Genome assembly procedures, related to Figure 1.

(A) Short read assembly procedure.

(B) Long read assembly procedure.

Figure S16. Genome structural characteristics, related to Figure 1.

(A) Ancestral state reconstruction of genome size (left in bp) and GC content (right as percent). The colour gradients indicate the genome size and GC content across the tree.

(B) Cumulative sequence covered by different features for each genome assembly: masked (repeated sequence) and unmasked regions (upper graph), intergenic and gene regions (middle graph) and intron UTR and CDS (lower graph).

Figure S17. Genome-wide analyses of composite and HGT-derived genes, related to Figure 2.

(A) Functional annotation status of fused, split and non-remodelled gene families (orthogroups).

(B) Example of the emergence of a gene family with a novel domain structure just prior to the emergence of the Phaeophyceae lineage. In brown algae and *S. ischiensis* orthogroup OG0007889 contains members with a novel domain structure combining domains (Interpro domains IPR041337 and IPR001660) found independently in orthogroups OG0006687 and OG0001104.

(C) Comparison of GC content and gene structure for different components of HGT-derived (HGT) and non-HGT-derived (core) genes. tssup, 5’ untranslated region; tssdown, 3’ untranslated region.

(D) Proportions of COG functional categories for HGT-derived and all genes. Numbers represent the percentage of the COG category for the considered gene set.

Figure S18. Intron conservation and spliceosome components, related to Figure 2.

(A) Intron conservation across a set of 235 conserved single-copy orthologous genes. The upper histogram shows the numbers of introns shared by the groups of species linked by black dots in the lower panel.

(B) Conservation of intron phase and size of the introns from each species compared with the *Ectocarpus* species 7 reference genome.

(C) Sequence logos of the donor and acceptor sites of introns from six selected species.

(D) Conservation of intron positions in *Ectocarpus* species and the outgroup *S. promiscuus* (right panel and insert).

(E) Phylogenetic tree of eukaryotic Lsm/Sm proteins. The tree was rooted using midpoint rooting. The colour code for the branches correspond to SmD3B (light green), SmD3A (dark green) and Lsm4 (grey).

(F) Phylogenetic tree of Lsm13 and Lsm14 proteins. The tree was rooted using midpoint rooting.

Figure S19. Analysis of long non-coding RNA genes in nine brown algal genomes and two closely-related taxa, related to Figure 1.

(A) Relative proportions of long non-coding RNA (lncRNA-like) and protein-coding (Coding-like) genes. M, male; F, female.

(B) Distributions of transcript length (left), length of longest ORF (middle) and percent GC (right) for lncRNA and protein-coding (mRNA) genes.

Figure S20. Analysis of organellar genomes, related to Figure 4.

(A) Loss of plastid genes during the evolution of the brown algae. Gene loss events were identified for five genes, with two of the genes (rbcR and rpl32) being lost more than once.

(B) Maximum likelihood phylogenetic trees for the brown algae based on genes from different genomic compartments. Phylogenetic tree based on plastid genes (top), phylogenetic tree based on mitochondrial genes (middle), discordance between the nuclear and plastid phylogenies and the mitochondrial phylogeny for the Ectocarpales (bottom). Ultrafast bootstrap values are indicated for 1000 replicates, with an asterisk indicating 100%.

Figure S21. Evolution of transcription-associated protein (TAP) families in the brown algae, related to Figure 3.

(A) Expansions, contractions, gains and losses of TAP families during the emergence of the brown algae. The number of gains (dark green), losses (dark red), expansions (light green) and contractions (light red) at each node is shown and the families involved are indicated using the same colour code. The light brown box indicates the Phaeophyceae.

(B) Distribution of transcription factors (TFs) across the brown algae and closely-related taxa. The bubbles indicate the relative abundance of 32 different TF families.

(C) Distribution of transcriptional regulators (TRs) across the brown algae and closely-related taxa. The bubbles indicate the relative abundance of 40 different TR families.

Figure S22. Histone structure and evolution, related to Figure 4

(A) Histone H3.1 and H3.3 isoforms differ at two residues located in the amino-terminal tail (31-32 AT for H3.1 and TA for H3.3, in green), three residues in the α2 helix of the histone fold domain (86-90 GSAVL for H3.1 and STAIL for H3.3) and one residue at the carboxy-terminal position (S or A, respectively in H3.1 and H3.3). Positions refer to the mature protein without the initial methionine.

(B) CenH3 proteins differ in their length and the amino acid composition of their amino-terminal tails. Strictly and highly conserved residues are depicted in black and red, respectively. The characteristic CATD (CENP-A targeting domain) is indicated by a red line. Positions refer to the mature protein without the initial methionine.

(C) Phylogenetic tree of histone H3.1 and H3.3 proteins of brown algae and other eukaryotes. Brown algae, diatoms, red algae, green algae, plant, animal and additional unicellular organisms (the myxomycetes *Physarum polycephalum* and *Dictyostellium discoideum*, the ciliate *Tetrahymena thermopila* and the yeast *Saccharomyces cerevisiae*) are depicted in brown, red, pink, light green, dark green, blue and black, respectively. See Methods section for species abbreviations.

(D) Schematic presentation of brown algal histone H1 linker proteins. The H1 variants are highly divergent across brown algae species. Strictly and highly conserved residues are depicted in black and red, respectively. Positions refer to the mature protein without the initial methionine.

Figure S23. Conservation and microevolution of genomes within the genus *Ectocarpus,* related to Figure 5.

(A) Dotplot illustrating genome-wide synteny between the *Ectocarpus* species 7 and *E. crouaniorum* genome assemblies. Horizontal lines delimit *E. crouaniorum* contigs.

(B) Gene family (orthogroup) gain and loss for the *Ectocarpus* genus based on a Dollo parsimony analysis. Cladogram branches are coloured according to the overall gain (+) or loss (-) of gene families. Grey circles on the nodes indicate the number of gene families predicted to be present.

(C) Predicted functions of genes in orthogroups gained within the *Ectocarpus* genus.

(D) Proportions of genes of different evolutionary ages, as estimated by phylostratigraphy analysis, on each chromosome of the *Ectocarpus* species 7 genome. Colour-coded groups of genes correspond to the following phylostratigraphy categories: 1, cellular organisms to phylum (ranks 1 to 5; orange); 2, PX clade to Phaeophyceae/*Schizocladia ischiensis* (ranks 6 to 7; yellow); 3, Phaeophyceae to subclass (ranks 8 to 10; green); 4, superorder to family (ranks 11 to 13; blue); 5, genus to species (ranks 14 to 15; pink).

(E) Distribution of synonymous (dS) and non-synonymous (dN) substitution rates for syntenic genes between *Ectocarpus* species 7 and *E. siliculosus*, *E. crouaniorum* or *E. fasciculatus* in relation to gene age based on the phylostratigraphy analysis. Correspondence between the colour-coded gene groups and age ranks are indicated on the left and are the same as in (D).

Figure S24. Evolution of signalling-related genes in the brown algae, related to Figure 3.

Overview of phytohormone biosynthesis pathways and distribution of gene families associated with biosynthesis across the brown algae and other stramenopile taxa. Left, biosynthesis pathways for the phytohormones abscisic acid (ABA), brassinosteroids (BR), cytokinins (CK; including iP, isopentenyladenine; tZ, trans-zeatin; DZ, dihydrozeatin; and cZ, cis-zeatin), ethylene (Eth), gibberellins (GA), auxin (IAA), jasmonic acid (JA), salicylic acid (SA) and strigolactones (SL). Right, Presence of putative homologs of phytohormone biosynthethic enzymes in brown algae based on EggNOG v5.0 gene families (root node). Colours denote z-score of the number of eggNOG family (left of heatmap) members. White denotes absence, (+) and (?) indicate that phytohormones have been reported to be present or have not been reported for brown algae (reviewed in^83,84^). ZEP, zeaxanthin epoxidase; NCED, nine-cis-epoxycarotenoid deoxygenate; SDR, short-chain dehydrogenase reductase; AAO, aldehyde oxidase; DET2, deetiolated2 (steroid 5alfa reductase); DWF4, dwarf4 (CYP90B); CPD, constitutive photomorphogenic dwarf (CYP90A); IPT, isopentenyl transferase; LOG, lonely guy (lysine de-carboxylase); SAMS, s-adenosyl methionine synthetase; ACS, ACC synthase; ACO, ACC oxidase; CPS, CDP/ent-kaurene synthase; KO, ent-kaurene oxidase; KAO, ent-kaurenoic acid oxidase; TAA, tryptophan aminotransferase; YUC, YUCCA (flavin monooxygenase); AMI, amidase; NIT, nitrilase; LOX, lipoxygenase; AOS, allene oxide synthase; AOC, allene oxide cyclase; OPR, oxo-phytodienoate reductase; ICS, isochorismate synthase; PAL, phenylalanine ammonia-lyase; D27, dwarf27 (all-trans/9-cis-B-carotene isomerase); CCD, carotenoid cleavage dioxygenase; MAX1, more axillary branches1 (CYP711A).

## Supplementary tables

Table S1. List of strains used for the project, the genome assemblies generated and accession numbers for sequence data.

(A) Strains used in the project.

(B) Genome assemblies.

(C) Statistical tests of correlations between genome assembly size and various genome features.

(D) Genomic data used in the project including both genomic DNA and RNA-seq data. For the latter, DE RNA-seq indicates replicated samples that were generated by experiments aimed at measuring differential gene expression.

(E) Genome assemblies and annotations generated by the project.

Table S2. Phylostratigraphy analysis.

(A) Gene ages estimated by phylostratigraphy. (B) Gene family founder events. (C) Gene ages after the homology detection failure correction. (D) Counts of founder events after the homology detection failure correction. (E) Statistics for *Ectocarpus* species 7.

Table S3. Gene families (orthogroups) significantly amplified in Phaeophyceae genomes compared to closely-related taxa.

OG, orthogroup; PHAEO, Phaeophyceae; FDI, Fucophycidae/Dictyotales/Ishigeales clade; LAMIN, Laminariales; ECTO, Ectocarpales.

Table S4. Core metabolic reactions most abundant in brown algae.

Metacyc IDs and reaction names (with EC numbers) for 24 genes present in all brown algae and less than 70% of outgroup species. Gene IDs can be retrieved from the GSMN Wikis using reaction IDs.

Table S5. Counts of CAZYme gene family members in brown algal and closely-related taxa genomes. Asterisks indicate species that were included in Figures S8A and S8B.

Table S6. CAZYmes identified in brown algal and closely-related taxa genomes. ProteinID, corresponds to the locusID; Description, CAZYme gene family.

Table S7. Algal-type vanadium haloperoxidase proteins encoded by brown algal and closely-related taxa genomes plus representative sequences from more distant taxa.

Table S8. Bacterial-type vanadium haloperoxidase proteins encoded by brown algal and closely-related taxa genomes plus representative sequences from more distant taxa.

Table S9. Type III polyketide synthase proteins encoded by brown algal and closely-related taxa genomes plus representative sequences from more distant taxa.

Table S10. Receptor kinase proteins encoded by brown algal and closely-related taxa genomes. mod, modified gene model; new, new gene model.

Table S11. DNA methyltransferase proteins encoded by brown algal and closely-related taxa genomes. Genome coordinates and protein sequences are provided for modified (mod) gene models.

Table S12. Predicted reference proteomes of the ten species analysed for generation-biased gene expression indicating sporophyte- and gametophyte-biased genes.

Table S13. Presence of flagellar protein genes in the genomes of nine brown algal species. P, present; A, absent.

Table S14. Mannitol cycle enzymes encoded by brown algal and closely-related taxa genomes. M1PDH, mannitol 1-phosphate dehydrogenase; M1Pase, mannitol 1-phosphate phosphatase; HK, hexokinase; M2DH, mannitol 2-dehydrogenase.

Table S15. Summary of the TAPscan output with lists of annotated transcription-associated proteins (TAPs) for brown algal and closely-related taxa genomes.

Table S16. Ion channels encoded by brown algae and other stramenopiles.

(A) Query sequences used for the screens. (B) Ion channel proteins detected. (C) Number of ion channels of each class per species. F, female; M, male; MCU, mitochondrial calcium uniporter; GLR, glutamate receptor; pLGIC, pentameric ligand-gated ion channel; TRP, transient receptor potential channel

Table S17. EsV-1-7 domain proteins encoded by brown algal and closely-related taxa genomes. Genome coordinates and protein sequences are provided for modified (mod) or newly created (new) gene models.

Table S18. Rates of synonymous and non-synonymous substitutions based on alignments of genes in syntenic blocks between *Ectocarpus* species 7 and four other *Ectocarpus* species.

Table S19. Inserted viral sequences in brown algal and closely-related taxa genomes.

(A) Summary data for figure 5. (B) List of viral genes. (C) List of viral regions. (D) Number of viral regions detected in each genome.

Table S20.Histidine kinase proteins encoded by brown algal and closely-related taxa genomes.

TM, transmembrane domain; CHASE, cyclases/histidine kinases associated sensory extracellular domain; EBD, ethylene-binding-domain-like; MASE1, membrane-associated sensor1 domain.

Table S21. Genomes studied for the various analyses carried out within the project. Y, genome analysed.

Table S22. Metabolism genes lost from either *Pleurocladia lacustris* or *Porterinema fluviatile* or from both, based on GSMN analysis.

Table S23.Histone proteins encoded by brown algae and other stramenopiles.

(A) Histone protein sequences. (B) Counts of histone proteins per genome. (C) Counts of histone genes per genome.

Table S24. *Porterinema fluviatile* genes differentially expressed under freshwater compared to seawater culture conditions.

Table S25. Tests for signatures of introgression among *Ectocarpus* species. Patterson’s D-statistic (ABBA-BABA tests) was calculated for concatenated alignments of 261 ortholog genes and significance was detected using a block-jackknifing approach with a block size of 5 Kbp. *Ectocarpus fasciculatus* was used as the outgroup taxon for all ABBA-BABA tests (noted as O in the four-taxon fixed phylogeny scheme: (((P1,P2)P3)O)). All values are significant.

Table S26. Spliceosome components identified in brown algal and closely-related taxa genomes. M, male; F, female.

Table S27. Statistics for the long non-coding RNA content of 11 genomes. M, male; F, female.

Table S28. Characteristics of plastid and mitochondrial genomes for 33 brown algae and *Chrysoparadoxa australica*.

Table S29. Presence or absence of 141 shared plastid genes across the brown algae. O, present; X, absent.

Table S30. Sulphatase proteins encoded by brown algal and closely-related taxa genomes.

Table S31. Integrin proteins encoded by brown algal and closely-related taxa genomes.

Genome coordinates and protein sequences are provided for modified (mod) or newly created (new) gene models. SP, signal peptide; TM, transmembrane domain; INTA, integrin alpha domain

Table S32. DEK1-like proteins encoded by brown algal and closely-related taxa genomes.

Table S33. Fasciclin proteins encoded by brown algal and closely-related taxa genomes.

Genome coordinates and protein sequences are provided for modified (mod) gene models. FAS1, fasciclin domain.

Table S34. Tetraspanin and tetraspanin-like proteins encoded by brown algal and closely-related taxa genomes.

Table S35. Comparative analysis of genes that were differentially expressed in freshwater compared with seawater in *E. subulatus* and *P. fluviatile* based on whether the genes are shared orthologues or lineage-specific.

Table S36. Blocks of syntenic genes shared between *Ectocarpus* species 7 and four other *Ectocarpus* species.

Table S37. Putative components of phytohormone biosynthetic pathways in brown algae.

## METHODS

## KEY RESOURCES TABLE

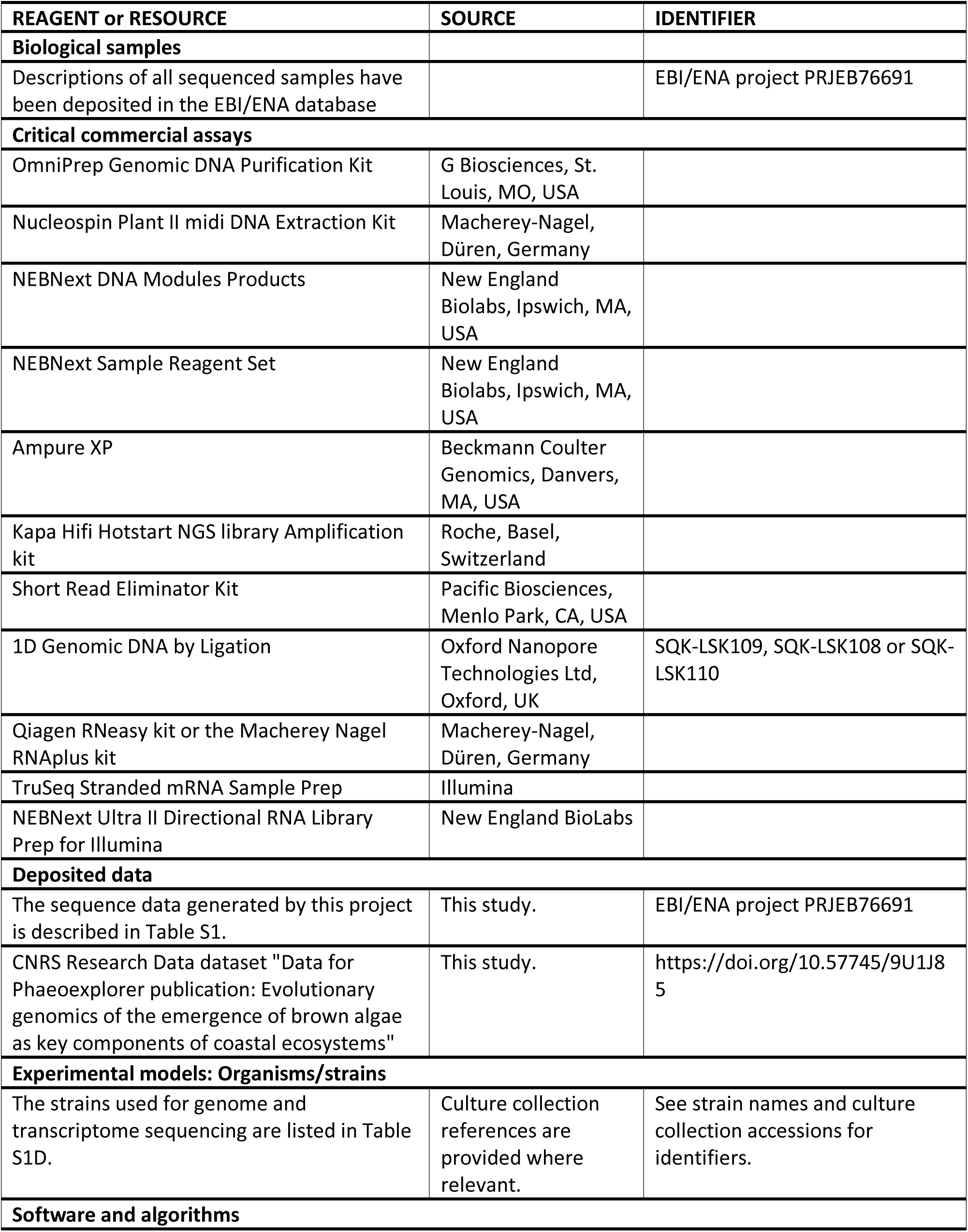
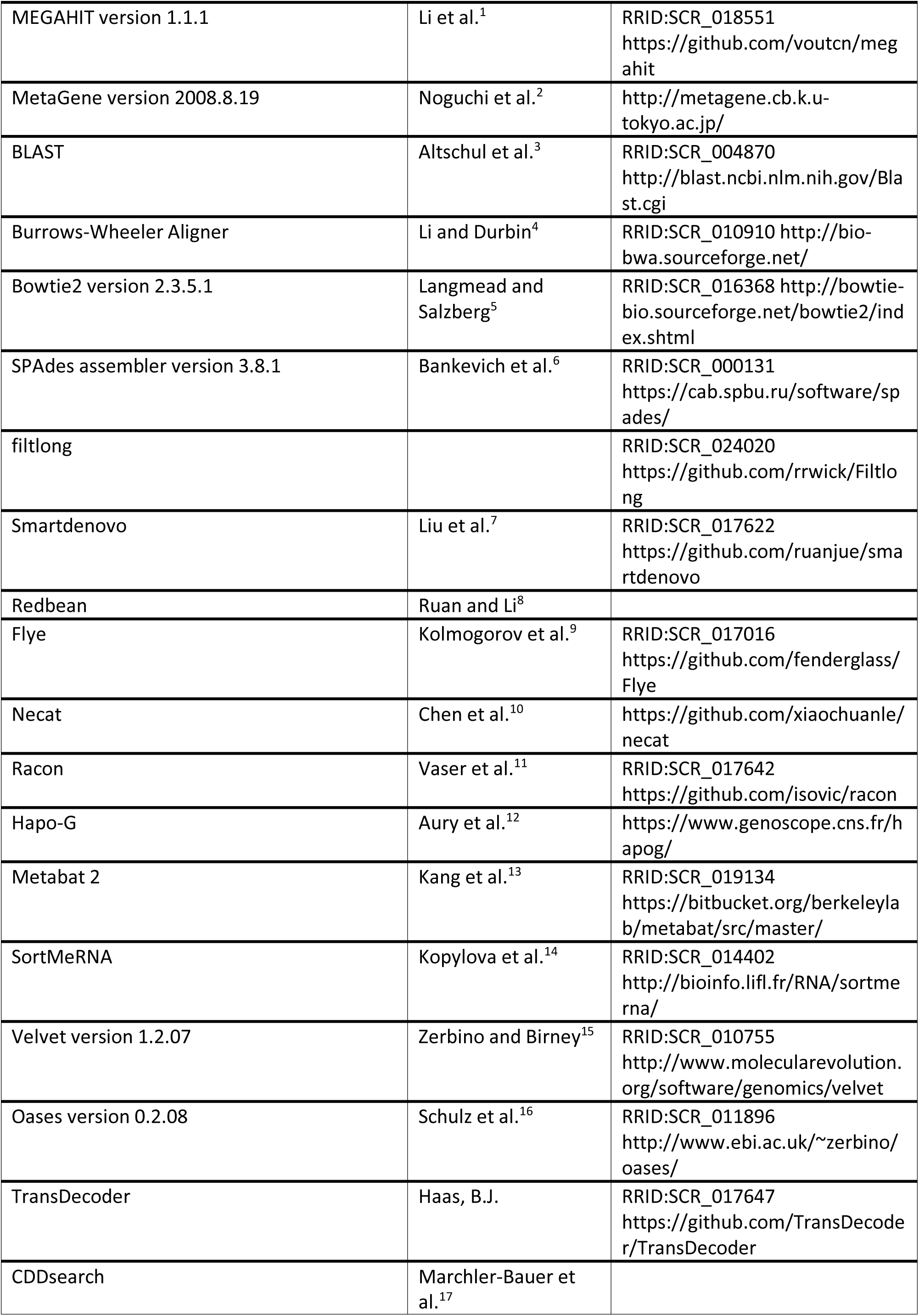
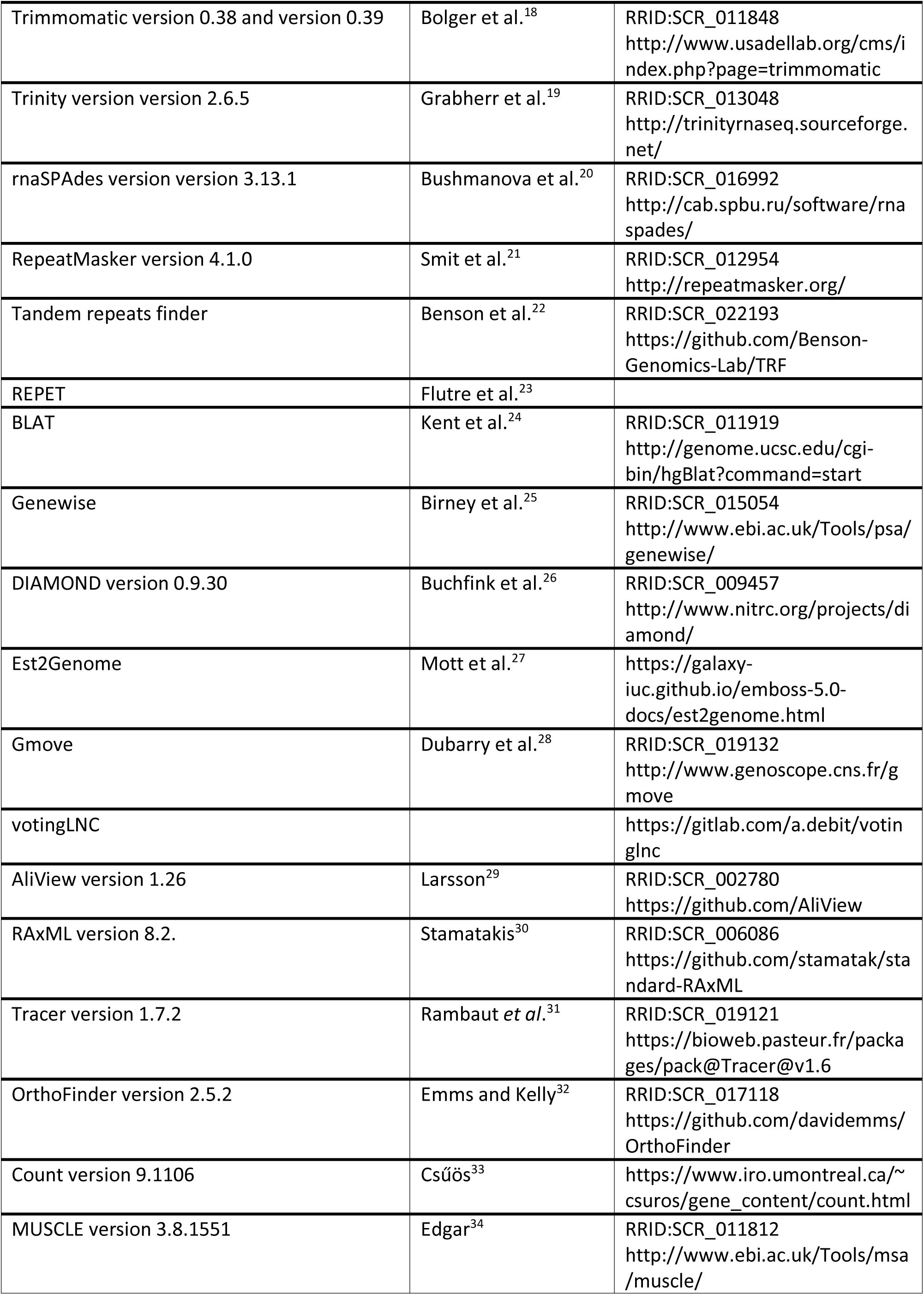
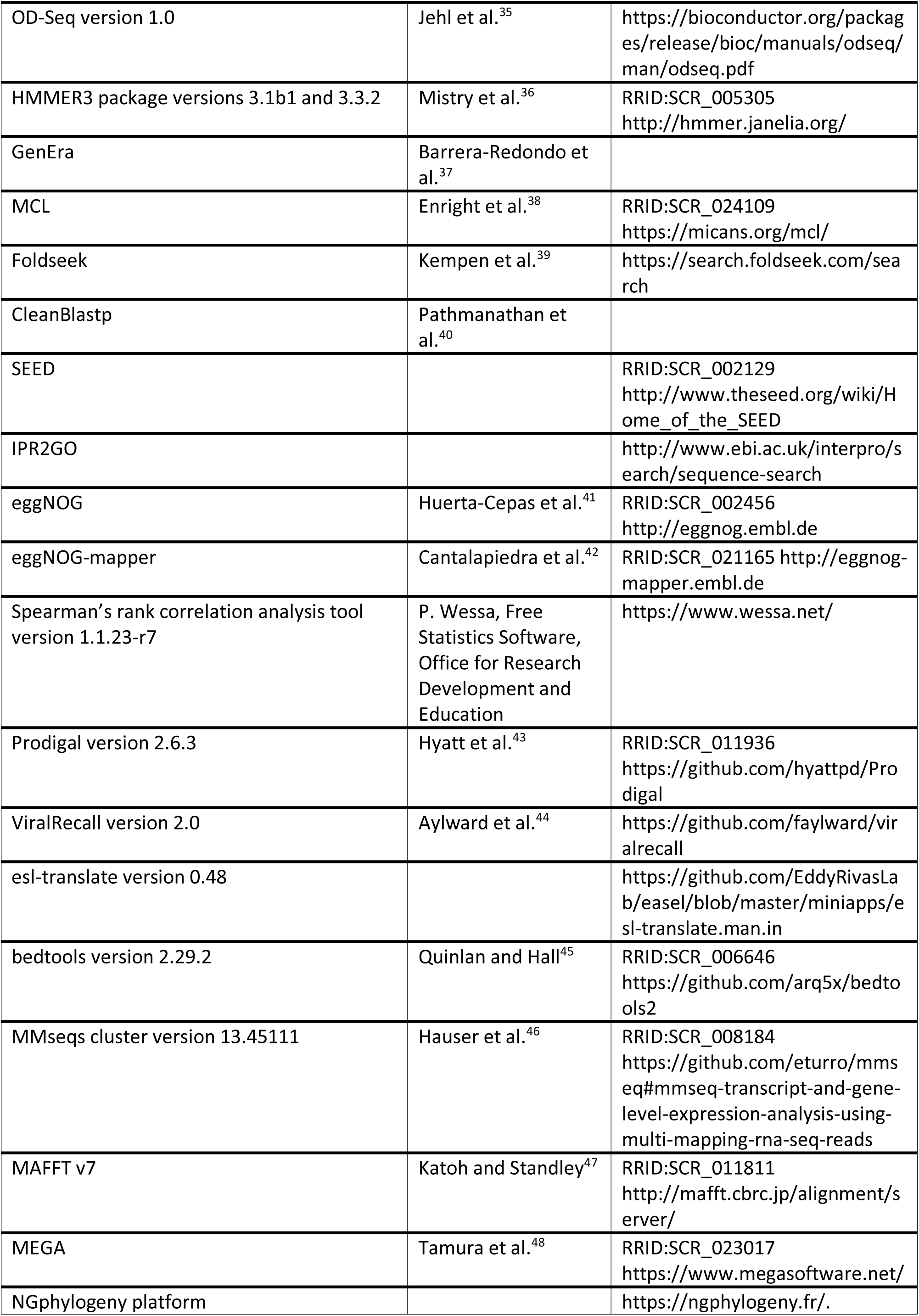
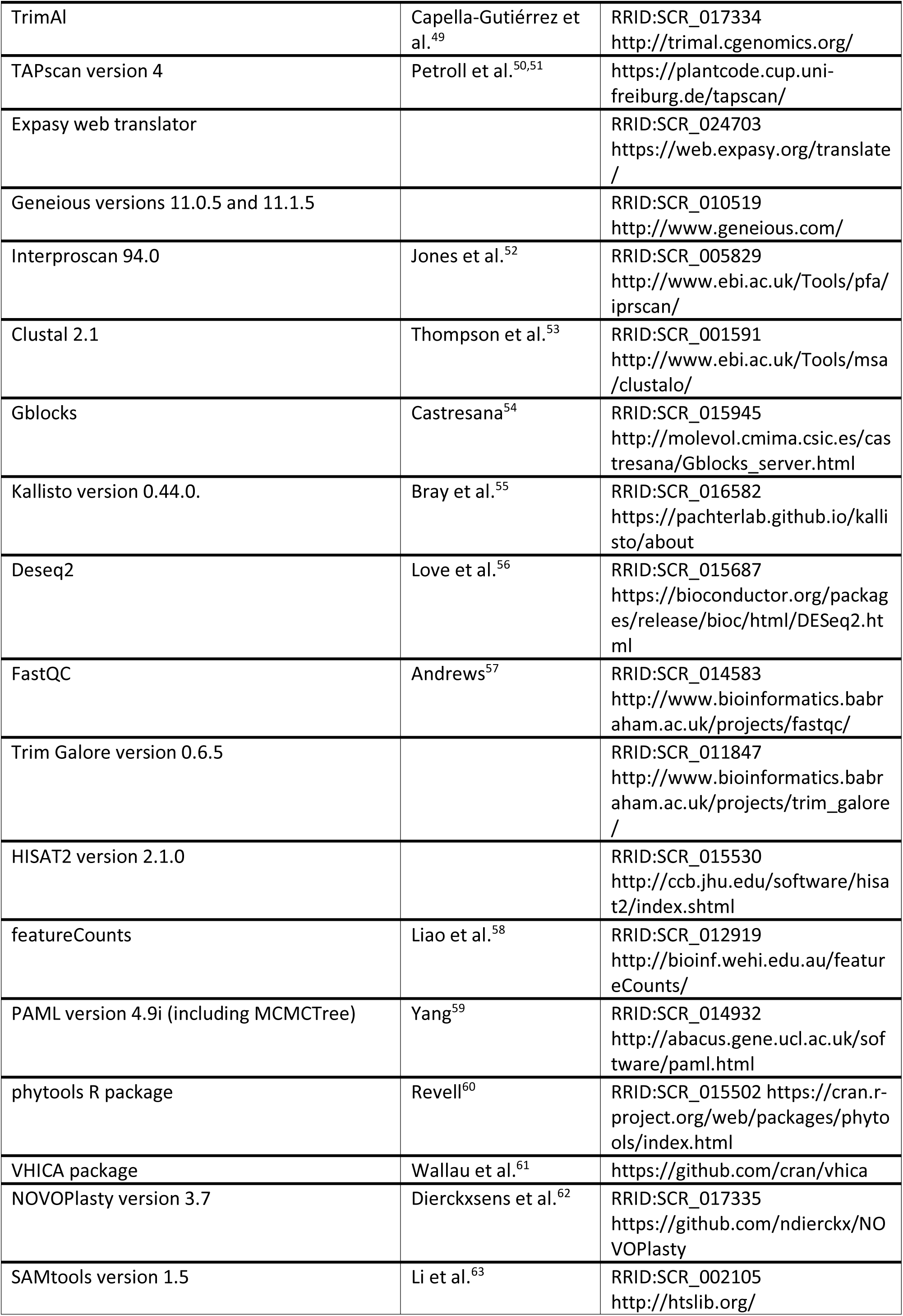
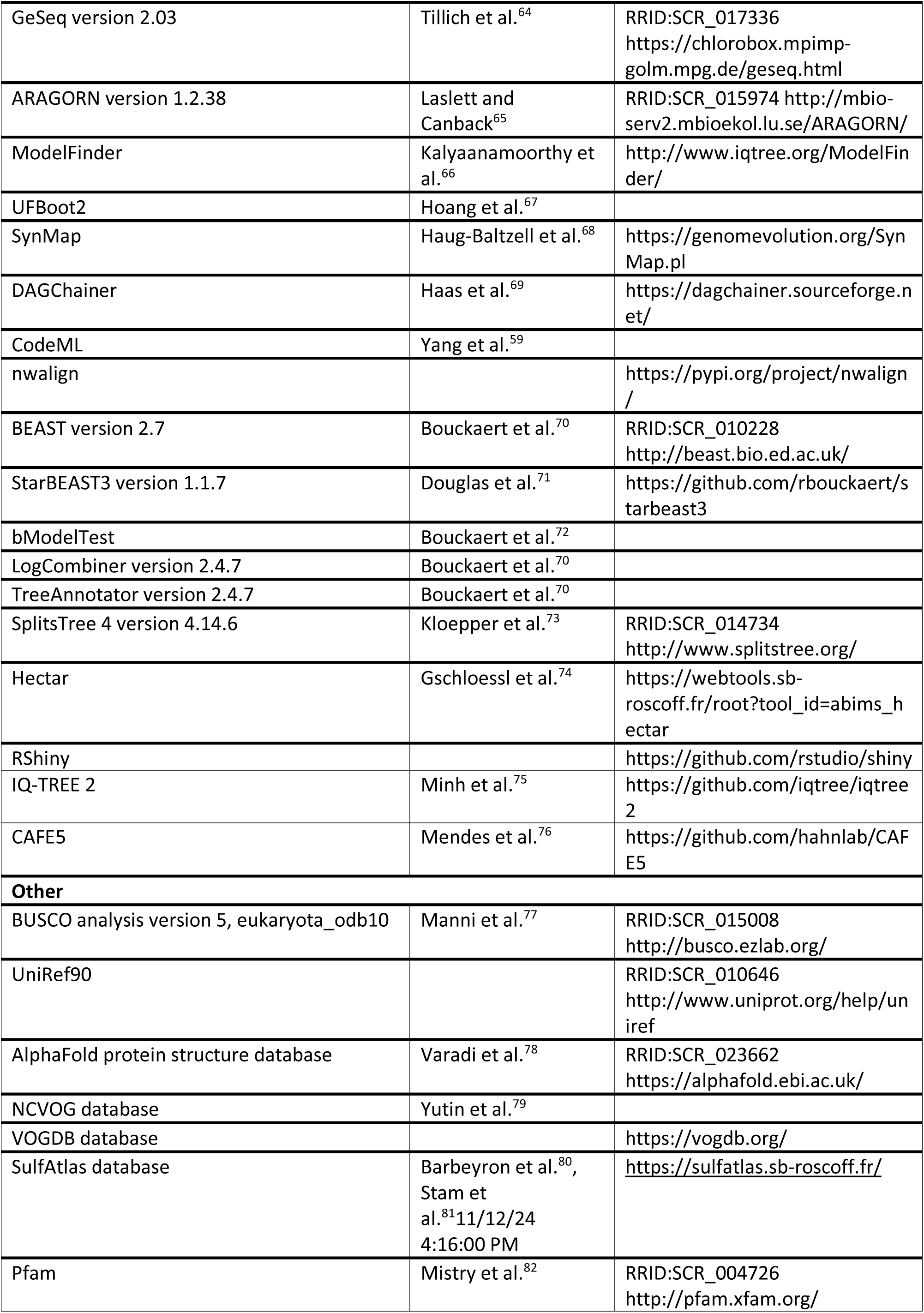
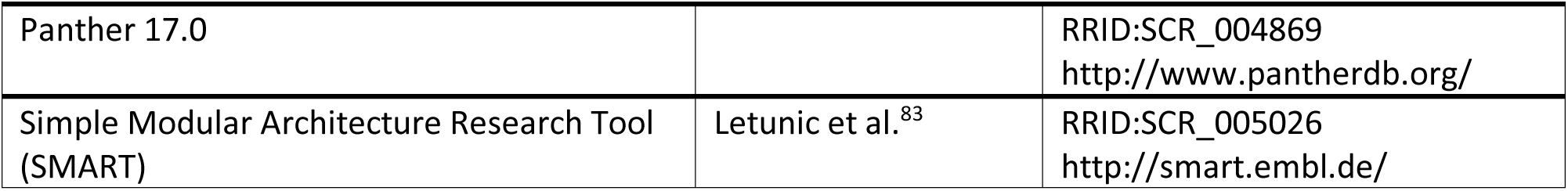

## RESOURCE AVAILABILITY

### Lead contact

Further information and requests for resources and reagents should be directed to and will be fulfilled by the lead contact, J. Mark Cock.

### Materials availability

All the laboratory-cultivated strains grown to provide material for genome sequencing can be accessed via the Roscoff Culture Collection (https://www.roscoff-culture-collection.org).

### Data and code availability

All sequence data, including DNA and RNA sequencing data, genome assemblies and annotations, have been deposited in the EBI/ENA database under the project accession PRJEB76691 and are publicly available. Additional data have been deposited in the CNRS Research Data depository under the title “Data for Phaeoexplorer publication: Evolutionary genomics of the emergence of brown algae as key components of coastal ecosystems” and are publicly available. The DOI is listed in the key resources table. Dataset description: “The Phaeoexplorer project sequenced 60 genomes corresponding to 44 brown algal and sister species. This dataset corresponds to supplementary information relating to the initial annotation of the Phaeoexplorer genomes and multiple analyses of the genome data. The dataset includes presubmission (v0) versions of the Phaeoexplorer genome annotation (GFF) files (GFF_v0.tar.gz) and genome-wide predicted proteomes as fasta files (Proteomes_v0.tar.gz), de novo transcriptome assemblies for the Phaeoexplorer species (RNA-seq data assembled with Trinity or rnaSPAdes; de-novo-transcriptomes.tar.gz), RepeatMasker analyses of repeat sequences (RepeatMasker.tar.gz), alignment files used to generate a phylogenetic tree for the Phaeoexplorer species (PhylogeneticTree.tar.gz), alignments used to build a densitree specifically for *Ectocarpus* species (Microevolution_Ectocarpus.tar.gz), an Orthofinder-based analysis of shared orthologues (Orthogroups.tar.gz) together with a Dollo-logic-based analysis of orthogroup gain and loss during evolution (Dollo_analysis.tar.gz), a Phylostratigraphy analysis of brown algal genes (Phylostratigraphy.tar.gz), an analysis of protein functional domain fissions and fusions (CompositeGenes.tar.gz), Interproscan analyses of protein domains (InterProScan.tar.gz), Hectar predictions of protein subcellular localisations (Hectar.tar.gz), eggNOG output providing information about predicted protein functions (eggNOG.tar.gz), RNA-seq-based data on gene expression levels (mRNAexpression.tar.gz), results of a search for genes acquired via horizontal gene transfer (HGT.tar.gz), analyses of intron conservation across genomes (Introns_conservation.tar.gz), an analysis of tandem gene duplications (Tandemely_duplicated_genes.tar.gz), comparisons of CDS size with the *Ectocarpus* reference genome that were used to evaluate gene model completeness (CDS_size.tar.gz), a DESeq2 analysis of differential gene expression between the sporophyte and gametophyte generations of several brown algal species (DEG_LifeCycle.tar.gz), information about orthogroups selected to analyse the effects of morphological complexity and life cycle structure on gene evolution (Genes_selection.tar.gz). Each individual dataset contains a README file explaining its content. Detailed information about the methodology used for each analysis can be found in the Methods section of the manuscript preprint (https://doi.org/10.1101/2024.02.19.579948). The majority of these analyses and datasets can also be accessed via the Phaeoexplorer website (https://phaeoexplorer.sb-roscoff.fr/).”

This paper does not report original code.

Any additional information required to reanalyse the data reported in this paper is available from the lead contact upon request.

## METHOD DETAILS

### Biological material

Sequencing brown algal genomes has been hampered by the significant challenges involved, including inherent problems with growing brown algae, the presence of molecules that interfere with sequencing reactions and complex associations with microbial symbionts. To address these problems, cultured, unialgal filamentous gametophyte material was used whenever possible (i.e. for species with haploid-diploid life cycles) and the extraction methodology was adapted for each species.

The algal strains analysed in this study are listed in Table S1, which provides information about the sampling site for each strain. The sampling sites are shown on a world map in Figure S2.

All strains except those belonging to the Fucales were grown under laboratory conditions. The latter cannot be maintained long-term in the laboratory so field material was harvested for extractions. The haploid gametophyte generation was grown in culture for species with characterised haploid-diploid life cycles, with the exception of *Ectocarpus* strains, for which haploid partheno-sporophytes or diploid sporophytes were cultivated. All cultures were grown either in 140 mm diameter Petri dishes or in 2– 10 L bottles, the latter aerated by bubbling with sterile air. Most cultures were grown in Provasoli-enriched^23^ natural seawater (PES medium) under fluorescent white light (10–30 μM photons/m2⋅s) at 13°C (or at 10°C for *Hapterophycus canaliculatus* and *Chordaria linearis* or 20°C for *Sphacelaria rigidula*, *Dictyota dichotoma, Schizocladia ischiensis* and *Chrysoparadoxa australica*). Exceptions included the freshwater species *Pleurocladia lacustris*, *Porterinema fluviatile* and *Heribaudiella fluviatilis*, which were grown in natural seawater that had been diluted to 5% with distilled water (i.e., 95% distilled water / 5% seawater) before addition of ES medium (http://sagdb.uni-goettingen.de/culture_media/01%20Basal%20Medium.pdf) micronutrients (at 20°C for *P. lacustris*) and *Phaeothamnion wetherbeei,* which was grown in MIEB12 (medium 7 in^84^). Whole thallus was extracted for all species except the Fucales, where either dissected meristematic regions or released male gametes were extracted. Tissue samples were frozen in liquid nitrogen and stored at −80°C before extraction.

### DNA extraction

DNA was extracted using either the OmniPrep Genomic DNA Purification Kit (G Biosciences, St. Louis, MO, USA) or the Nucleospin Plant II midi DNA Extraction Kit (Macherey-Nagel, Düren, Germany). DNA quality was assessed using a Qubit fluorometer (Themo Fisher Scientific, Waltham, MA, USA), and fragment length was assessed by migration on a 1% agarose gel for some of the samples.

### Illumina library preparation and sequencing

Libraries were prepared using the NEBNext DNA Modules Products (New England Biolabs, Ipswich, MA, USA) with an ‘on bead’ protocol developed by Genoscope, starting with 100 ng of genomic DNA. DNA was sonicated to a 100–800 bp size range using a Covaris E220 sonicator (Covaris, Woburn, MA, USA), end-repaired and 3ʹ-adenylated. Illumina adapters (Bioo Scientific, Austin, TX, USA) were then added using the NEBNext Sample Reagent Set (New England Biolabs, Ipswich, MA, USA) and the DNA purified using Ampure XP (Beckmann Coulter Genomics, Danvers, MA, USA). Adapted fragments were amplified with 12 cycles of PCR using the Kapa Hifi Hotstart NGS library Amplification kit (Roche, Basel, Switzerland), followed by 0.8x AMPure XP (Beckman Coulter Genomics, Danvers, MA, USA) purification. Libraries were sequenced with Illumina MiSeq, HiSeq 4000 or NovaSeq 6000 instruments (Illumina, San Diego, CA, USA) in paired-end mode, 150 base read-length.

### Oxford Nanopore library preparation and sequencing

Some samples were first purified using the Short Read Eliminator Kit (Pacific Biosciences, Menlo Park, CA, USA). All libraries were prepared using the protocol “1D Genomic DNA by Ligation” provided by Oxford Nanopore Technologies (Oxford Nanopore Technologies Ltd, Oxford, UK). Most of the libraries were prepared with the SQK-LSK109 kit (Oxford Nanopore Technologies), a few with the SQK-LSK108 or SQK-LSK110 kits (Oxford Nanopore Technologies). Three flow cells were loaded with barcoded samples. The samples were mainly sequenced on R9.4.1 MinION or PromethION flow cells.

### RNA extraction, Illumina RNA-seq library preparation and sequencing

RNA was extracted using either the Qiagen RNeasy kit or the Macherey Nagel RNAplus kit (Macherey-Nagel, Düren, Germany). RNA-seq libraries were prepared using the TruSeq Stranded mRNA Sample Prep (Illumina) according to the manufacturer’s protocol, starting with 500 ng to 1 µg of total RNA, or using the NEBNext Ultra II Directional RNA Library Prep for Illumina (New England BioLabs) according to the manufacturer’s protocol, starting with 100 ng of total RNA. The libraries were sequenced with Illumina HiSeq 2500, HiSeq 4000 or NovaSeq 6000 instruments (Illumina, San Diego, CA, USA), in paired-end mode, 150 base read-length.

### Assembly strategies

Two assembly strategies were employed (Figure S15): one was designed for genomes exclusively sequenced using short reads with Illumina technology, while the other was designed for genomes that underwent sequencing using a combination of long and short reads, using respectively the Nanopore and Illumina technologies.

### Short-read-based genome assembly

When sequencing was performed exclusively using short reads, reads corresponding to bacterial contaminants were filtered out early in the assembly process (Figure S15A) because, typically, the initial datasets were too large to run assemblers like SPAdes. To remove bacterial contaminants, an assembly based on the initial illumina dataset was first generated for each strain using a fast and non-greedy algorithm, MEGAHIT^1^ version 1.1.1 with the parameters --k-min 101--k-max 131 --k-step 10. Assigning taxonomy is easier when working with contigs than with reads. Contigs exceeding 500 bp in each preliminary assembly underwent taxonomic classification based on gene models predicted using the *ab initio* software MetaGene^2^ version 2008.8.19 with default parameters and then aligning proteins against UniprotKB using BLASTp (e-value <10e^-4^). A superkingdom (Eukaryota, Archaea or Bacteria) was assigned to each gene based on the best alignment (selected using the BLASTp score). Contigs that contained more than 50% of their genes assigned to Bacteria and with at least one gene every 10 Kbp were classified as bacterial sequences. For each strain, the initial Illumina sequencing reads were aligned against the corresponding bacterial sequences using latest version of the Burrows-Wheeler Aligner^4^ (BWA) with default parameters and mapped short-reads were labelled as contaminants, and assembled for the purpose of obtaining more contiguous contigs. These bacterial contigs were then used to build a contaminant sequence database. Finally, the clean subset of reads was obtained by aligning the whole Illumina dataset against this strain-specific bacterial contig database, using Bowtie2^5^ version 2.2.9 with default parameters. A final assembly was then generated for each strain using the contaminant-free read datasets and the SPAdes^6^ assembler version 3.8.1 with the parameters -k 21,57,71,99,127 -m 2000 --only-assembler -- careful. Genome assemblies based only on short-reads were more fragmented (N50 ranged from 3 Kbp to 31 Kbp) than assemblies that used long reads but the sizes of the former were consistent with expectations.

### Long-read-based genome assemblies

A subset of the strains produced DNA of both adequate quality and quantity, enabling successful long-read sequencing. In these cases, long reads were assembled directly and the detection of possible bacterial contigs was carried out after the assembly step (Figure S15B). To produce long-read-based genome assemblies we generated three samples of reads i) all reads, ii) 30X coverage of the longest reads and iii) 30X coverage of the filtlong (https://github.com/rrwick/Filtlong) highest-score reads. The three samples were used as input data for four different assemblers, Smartdenovo^7^, Redbean^8^, Flye^9^ and Necat^10^. Based on the cumulative size and contiguity, we selected the best assembly for each strain. This assembly was then polished three times using Racon^11^ with Nanopore reads, and twice with Hapo-G^12^ and Illumina PCR-free reads.

### Assembly decontamination

Contigs from the short- and long-read genome assemblies were inspected for potential bacterial sequences. This process was carried out using a combination of several analysis and tools: GC composition, read coverage, Metabat 2 (for tetramer composition and clustering)^13^ and Metagene (for gene prediction and taxonomic identification, as described previously). Contigs were manually removed based on their characteristics.

### Transcriptome assembly

Ribosomal-RNA-like reads were detected using SortMeRNA^14^ and filtered out. The Illumina RNA-Seq short reads from each strain were assembled using Velvet^15^ version 1.2.07 and Oases^16^ version 0.2.08 with kmer sizes of 61, 63 and 65 bp. BUSCO^77^ analysis (v5, eukaryota_odb10) was then performed on the three resulting assemblies for each strain in order to select the best assembly, i.e. the most complete at the gene level. Reads were mapped back to the contigs with BWA-mem, and only consistent paired-end reads were retained. Uncovered regions were detected and used to identify chimeric contigs. In addition, open reading frames (ORF) and domains were identified using TransDecoder (Haas, B.J., https://github.com/TransDecoder/TransDecoder) and CDDsearch^17^, respectively. Contigs were broken into uncovered regions outside ORFs and domains. In addition, read strand information was used to correctly orient RNA-Seq contigs.

### *De novo* transcriptomes

The RNA-seq data was also used to generate *de novo* transcriptomes. For each strain, all the RNA-seq data available was cleaned to remove poor quality sequence and adapter sequences using Trimmomatic^18^ v0.39 prior to being assembled using either Trinity^19^ version v2.6.5 or rnaSPAdes^20^ version v3.13.1. The strandness and Kmer-length parameters of the assemblers were adjusted to take into account RNA-seq read characteristics. The *de novo* transcriptomes represented an alternative source to identify and characterise genes if they were not detected in the genome assemblies. The *de novo* transcriptomes are available from the CNRS Research Data dataset (https://doi.org/10.57745/9U1J85) and from the Phaeoexplorer website (https://phaeoexplorer.sb-roscoff.fr/).

### Detection and masking of repeated sequences and transposons

Prior to gene annotation, each genome assembly was masked based on the repeat library from *Ectocarpus* species 7 (formerly *Ectocarpus siliculosus*)^85^ and using RepBase with RepeatMasker^21^ version v.4.1.0, default parameters. Tandem repeats finder (TRF)^22^ was also used to mask tandem repeat duplications. In addition, transposons were annotated in ten species using REPET^23^ and the transposons detected were used as a reference to mask all genomes with RepeatMasker^21^ version v4.1.0, default parameters.

### Gene prediction

For each strain, gene prediction was performed using both homologous proteins and RNA-seq data. Proteins from *Ectocarpus* species 7 (https://bioinformatics.psb.ugent.be/orcae/overview/EctsiV2)^86^ and UniRef90 (https://www.uniprot.org/uniref/) were aligned against each genome assembly. First, BLAT^24^ with default parameters was used to quickly localise putative genes corresponding to the *Ectocarpus* species 7 proteins. The best match and matches with a score ≥ 90% of the best match score were retained. Second, the alignments were refined using Genewise^25^ with default parameters, which is more precise for intron/exon boundary detection. Alignments were retained if more than 80% of the length of the protein was aligned to the genome. To detect conserved proteins and allow detection of horizontal gene transfer, UniRef90 proteins (without *E. siliculosus* sequences) were aligned with DIAMOND^26^ (v0.9.30 with parameters --evalue 0.001 --more-sensitive) to genomic regions lacking alignments with an *Ectocarpus* species 7 protein. Only the five best matches per locus were retained, based on their bitscore. Selected proteins from UniRef90 were aligned to the whole genome using Genewise as described previously, and alignments with at least 50% of the aligned protein length were retained. The assembled transcriptome for each strain was aligned to the strain’s genome assembly using BLAT^24^ with default parameters. For each transcript, the best match was selected based on the alignment score, with an identity greater or equal to 90%. Selected alignments were refined using Est2Genome^27^ in order to precisely detect intron boundaries. Alignments were retained if more than 80% of the length of the transcript was aligned to the genome with a minimal identity of 95%. Finally, the protein homologies and transcript mapping were integrated using a combiner called Gmove^28^. This tool can find coding sequences (CDSs) based on genome-located evidence without any calibration step. Briefly, putative exons and introns, extracted from the alignments, were used to build a simplified graph by removing redundancies. Then, Gmove extracted all paths from the graph and searched for open reading frames (ORFs) consistent with the protein evidence. Translated proteins of predicted genes were then aligned against NR prot (release 19/02/2019) and the *Ectocarpus* species 7 version v2 proteome^86^ (https://bioinformatics.psb.ugent.be/orcae/overview/EctsiV2) using DIAMOND BLASTp with parameters --evalue 10-5 --more-sensitive --unal 0. All predicted genes with significant matches (the smallest protein had to be aligned for at least 50% of its length) were retained. In addition to these genes, we also retained genes with CDS size greater than 300 bp and with a coding ratio (CDS size / mRNA size) greater or equal to 0.5.

### Annotation decontamination

After predicting the genes, an additional analysis was carried out to detect bacterial sequences. If a contig did not contain any genes, it was analysed with MetaGene and the predicted proteins added to the gene catalogue for the purpose of detecting bacterial sequences. Proteins generated from predicted genes (Gmove plus MetaGene) were then aligned against UniprotKB using BLASTp (e-value < 10e^-4^) and superkingdom (Eukaryota, Archaea or Bacteria) was assigned to each gene based on the best alignment (selected using the BLASTp score). Contigs that contained more than 80% of their genes assigned to bacteria, Archaea or viruses were classified as bacterial sequences and removed from the final assembly file. Genes belonging to these contigs were also removed from the final gene catalogue. Finally, completeness of each predicted gene catalogue was assessed using BUSCO^77^ (v5.0.0; eukaryota_odb10).

In addition, the quality of the annotations was assessed by comparing the length of coding regions in pairs of orthologous proteins (best reciprocal hits) between each genome and *Ectocarpus* species 7, which was used as a reference because its high-quality annotation has been extensively curated^86^. The correlation between orthologous CDS lengths was higher for genomes sequenced with long reads than for genomes only sequenced with short reads (Figure S1B). This difference was probably principally due to a higher proportion of underestimated protein lengths in the latter (Table S1B) which likely corresponded to fragmented genes. The qualities of Ectocarpales genome annotations were very high (BUSCO and length of predicted CDS) even when the genomes were sequenced using only short reads, probably because their phylogenetic proximity to *Ectocarpus* species 7 facilitated the building of good quality gene models.

### Analyses aimed at deducing functional characteristics of predicted proteins

Several different analyses of the predicted proteomes of each species were carried out to provide information about the cellular functions of the encoded proteins. These included eggNOG-mapper^42^ analyses (v2.1.8 or v2.0.1, with emapperDB v5.0.2 or v4.5.1) to provide multiple functional annotations (Gene Ontology, Kyoto Encyclopedia of genes and genomes, Clusters of Orthologous Genes, Pfam), Interproscan^52^ analyses (versions v5.55-88.0, v5.51-85.0 or v5.36-75.0) to detect functional domains, Hectar^74^ (v1.3) predictions of protein subcellular localisation and various DIAMOND^26^ (v2.0.15 vs UniRef90 2022_03, with parameter “evalue” set to 10e^-5^) sequence similarity searches aimed at identifying homologous proteins with functional annotations.

### Detection of tandemly duplicated genes

Starting with the protein alignments that had been constructed to build the orthogroups, matches between proteins within the same genome with an e-value of ≤10−20 and which covered at least 80% of the smallest protein were extracted. Two genes were considered to be tandemly duplicated if they were localised on the same genomic contig separated by five or less intervening genes, regardless of their orientation. The tandemly-duplicated genes were clustered using a single linkage clustering approach. A contingency test was applied to compare the proportion of tandemly-duplicated genes in each orthogroup with the global proportion of tandemly-duplicated genes (p=0.0532792). The *p*-values are shown in table S1.

### Relative orientation of adjacent genes and lengths of intergenic regions

For each species, the proportion of pairs of adjacent genes localized on opposite strands was compared to the expected proportion of 0.5 using a binomial test (with p=0.5). The *p*-values are shown in Table S1 (*p*-values of <0.05 correspond to cases where the proportion is significantly higher than 0.5).

The lengths of intergenic regions between pairs of adjacent genes located on opposite strands (i.e. divergently or convergently transcribed) were compared with the lengths of intergenic regions between genes located on the same strand (i.e. transcribed in the same direction). Contingency tables were constructed for each species using a threshold of 1000 bp for the intergenic length and the number of intergenic regions in each of four categories were counted: 1) same strand genes, intergenic <1000 bp, 2) opposite strand genes, intergenic <1000 bp, 3) same strand genes, intergenic ≥1000 bp, 4) opposite strand genes, intergenic ≥1000 bp. Fisher exact tests were applied to the contingency tables (alternative hypothesis: true odds ratio is greater than 1). The *p*-values are shown in Table S1. When *p*-values are <0.05, short intergenic lengths are significantly associated with pairs of genes on opposite strands. All calculations were performed with R^87^ (version 4.3.0).

### Detection of long non-coding RNAs

Transcriptome data for 11 species (Table S21), including nine brown algal strains and two outgroup taxa, was analysed to identify lncRNAs. Any transcripts with invalid nucleotide DNA symbols were discarded and sequences shorter than 200 nucleotides were removed to avoid the detection of small RNA transcripts. The transcriptome sequences in Fasta format were analysed with votingLNC (https://gitlab.com/a.debit/votinglnc) to detect lncRNA transcripts and assign a confidence level for each transcript. A similar approach was used to detect lncRNAs in the lncPlankton database^88^. VotingLNC is a meta-classifier combining the predictions of the ten most commonly used coding potential tools. Based on a majority voting ensemble procedure, the meta-tool assigns the final coding potential class to a transcript as the class label predicted most frequently by the ten classification models included in the ensemble. Alongside the majority voting class, a reliability score was calculated for each transcript. A cut-off non-coding reliability score of *p* > 0.5 was chosen to treat a transcript as lncRNA and to decrease false-positive identification. The set of transcripts predicted as lncRNA by the majority-voting procedure and having an ORF(s) encoding peptide(s) with length ≥ 100aa were discarded. lncRNA transcripts that had significant matches in either the Pfam^82^ (hmmscan e-value < 0.001) or SwissProt (BLASTp e-value < 1e^-5^ and similarity ≥ 90%) databases were removed from the dataset. Transcript length, GC content, and the length of the longest ORF were compared between lncRNAs and protein-coding RNAs. The comparison was carried out using a Wilcoxon test. R version V.4.1.2 was used for all the analyses and ggplot2 (version 3.4.0) for plotting.

### Intron conservation

Intron positions were compared in a set of single copy genes that are conserved across all the Phaeophyceae and the outgroup species. The analysis focused on the 21 reference genomes (Table S21) and on orthogroups that occurred exactly once in at least 20 of the 21 genomes, allowing the gene to be absent from only one of the 21 genomes. In addition, orthogroups were discarded if more than three copies had been annotated in the other Phaeophyceae genomes. These filters produced a set of 235 conserved (ancestral) orthogroups. Multiple alignments were carried out for each orthogroup using MUSCLE^34^ version 3.8.1551 with default parameters and conserved blocks were identified with Gblocks^54^ version 0.91b with the parameters -p=t -s=n -b5=a -b2=[nsp] -b1=[nsp] -b3=6, where “nsp” is equal to 90% of the number of proteins aligned. A shell script was then used to compare intron positions in the alignments. For each intron in the multiple sequence alignment, we obtained a corresponding conservation profile listing which species contains an intron at that position. The profiles obtained for the 949 introns that are in conserved blocks of the multiple alignments are shown in Figure S6B. Both phase and length of ancestral introns (e.g. that were conserved in most Phaeophyceae and at least two sister clades) were compared to the phase and length of *Ectocarpus* species 7 introns as a reference. The same approach was used to compare intron positions across 11 *Ectocarpus* species, with *Scytosiphon promiscuus* as an outgroup, by selecting 831 conserved monocopy orthogroups. The number of introns per gene in brown algae and in closely-related outgroup species were compared using a contingency test (table S1C).

### Phylogenomic tree of the Phaeophyceae

To provide a phylogenetic framework for the analyses of the Phaeoexplorer genome dataset, the 41-species phylogenomic tree reported by Akita *et al.*^89^ was updated by adding 15 additional species using the same methodology. Briefly, for the additional species, amino acid sequences were recovered for the 32 single-copy orthologous genes used to construct the published tree and these were aligned manually with the existing sequences using the alignment software AliView^29^ v.1.26. The aligned sequences of the final 56 species were concatenated and maximum likelihood analysis was carried out with 10,000 rapid bootstraps using RAxML^30^ v.8.2.9 and the gamma model. The best-fit evolutionary model for each gene was determined using AIC.

### Bayesian divergence time estimation for the brown algae

An estimation of brown algal divergence time was carried out using the 32 orthologous nuclear genes (see above and^89^) for 51 brown algae and five non-brown species (16,185 amino acids, 56 spp.) and MCMCTree (PAML package v4.9j) with the approximate likelihood method. The WAG protein model was selected based on the AIC and BIC criteria of ModelFinder^66^. The independent clock model was selected based on previous work on the brown algal timeline by Choi *et al*.^90^. One hundred million years was set to correspond to 1 in the MCMCTree calculation. A secondary calibration for the root was based on Choi *et al*.^90^ using a gamma distribution of 70.2 alpha and 10.22 beta. A kelp holdfast fossil^91^ was used to date the crown node of kelps with a minimum bound of 0.31, and a *Julescraneia* fossil^92^ for the *Macrocystis/Saccharina* clade with a minimum bound of 0.13 (Figure S3). MCMC chains were run 1.5 million generations, with the first 200,000 MCMC chains being discarded as burn-in, and the convergence of MCMC chains was checked with Tracer v1.7.2^31^. This analysis estimated that Schizocladiophyceae and brown algae diverged 457.88 Mya (95% HPD: 321.29-592.66 Ma), similar to (about 8 Mya older than) the previous estimate using plastid genes^90^ and that diversification of the major brown algal linages began about 220 million years later, after the origin of DFI clade (235.97 Mya, 95% HPD: 158.88-312.48 Mya), about 12 Ma earlier than the previous estimate^90^. The fossil-calibrated phylogenetic tree for 11 *Ectocarpus* species (Figure S13A) was extracted from the brown algal tree (Figure S3).

### Detection of orthologous groups

Predicted proteins from the 60 strains sequenced in Phaeoexplorer complemented with 16 public proteomes covering the Ochrophytina subphylum and the terrestrial oomycetes were clustered using OrthoFinder^32^ v2.5.2 with default parameters. This generated 56,340 orthogroups that contained 90.1% of the proteins (1,415,341 of the 1,571,648). Seventy-one of the 76 strains had more than 75% of their proteins in an orthogroup shared with at least one other strain. The orthogroups contain between 2 and 6,220 proteins with a mean of 25.1 proteins and a median of three.

### Dollo analysis of orthogroup gain and loss

An analysis of evolutionary events of gene family gain and loss was carried out on a selection of strains covering the brown algal phylogeny and sister groups as distant as the Raphidophyceae under the Dollo parsimony law using orthogroups as proxies for gene families. To limit possible problems due to the fragmentation of predicted proteins in some assemblies, we selected 24,410 orthogroups present in at least one of 17 strains that had both good quality genome assembly and good quality gene predictions. Dollo parsimony analysis was then run using Count^33^ version v9.1106 based on a cladogram of a subset of 24 species representative of the Phaeoexplorer project and excluding all public outgroups more distant than *Heterosigma akashiwo*. The cladogram was based on the topology of the brown algae phylogenetic tree published by Akita *et al*.^89^.

### Phylostratigraphy analysis

GenEra^37^ was used to estimate gene family founder events for each genome assembly by running DIAMOND^26^ in ultra-sensitive mode against the Phaeoexplorer protein dataset and the NCBI non-redundant database. All sequence matches with e-values < 10^-5^ were treated as being homologous with the query genes in the target genomes. The NCBI taxonomy was used as an initial template to infer the evolutionary relationships of each query gene with their matches in the sequence database but taxonomic assignments within the PX clade and Phaeophyceae were then modified to reflect the evolutionary relationships that were inferred in the maximum likelihood tree. Gene families were predicted based on a clustering analysis of the query proteins against themselves using an e-value cutoff of 10^-5^ in DIAMOND and an inflation parameter of 1.5 with MCL^38^. Estimated evolutionary distances were extracted for each pair of species from the maximum likelihood species tree (substitutions/site) to calculate homology detection failure probabilities^93^. Taxonomic sampling of the species tree enabled homology detection failure tests to be carried out within the PX clade. Gene families whose ages could not be explained by homology detection failure were analysed by inspecting the functional and domain annotations for *Ectocarpus* species 7^86^. Structural alignments were performed using Foldseek^39^ against the AlphaFold protein structure database^78^.

### Detection of gene family amplifications

A binomial test with a parameter of 17/21 was carried out to detect gene families (OGs) that had significantly expanded in 17 Phaeophyceae reference genomes compared with four closely-related outgroup species (*Schizocladia ischiensis*, *Tribonema minus*, *Chrysoparadoxa australica* and *Heterosigma akashiwo*; Table S21). Expanded gene families deviated significantly from the expected proportion (17/21 under the null hypothesis where there are equal gene numbers in all species). Benjamini–Hochberg FDR correction for multiple testing was then applied and 233 candidate OGs with corrected *p*-values of < 0.001 were retained. All calculations were performed with R (version 4.1.0).

The set of 233 candidate OGs was then filtered to limit counting errors due to annotation artefacts (e.g. genes missed or fragmented) using the following procedure:

1. A protein consensus was first deduced for each orthogroup. Protein sequences representative of all lineages were extracted and aligned using MUSCLE^34^ version 3.8.1551 with default parameters and the multiple alignments were filtered using OD-Seq^35^ version 1.0 to remove outlier sequences, with parameter –score set to 1.5. The consensus sequences were then extracted from the multiple alignments of non-outlier sequences using hmmemit in the HMMER3^36^ package version 3.1b1 with default parameters.
2. In order to estimate gene family copy number independently of the assembly and annotation processes, short read sequences for each genome were mapped onto the orthogroup consensus sequences using DIAMOND^26^. Unique matches were retained for each read and depth of coverage was calculated for each consensus orthogroup. The depth obtained for each orthogroup was normalised for each species by dividing by the depth obtained on a set of conserved single-copy genes, so that the final value obtained was representative of the gene copy number. Then, for each candidate amplified orthogroup, the average depth for the 17 Phaeophyceae species and the average depth for the four outgroup species was calculated and OGs where the depth for outgroups was more than half the depth for the Phaeophyceae were discarded. We retained 227 out of 233 orthogroups after this step.
3. Finally, functional annotations were used to remove orthogroups that were likely to correspond to transposable elements. A final list of 180 OGs was retained (Table S3).

The amplified gene families were manually categorised into functional classes based on the output of automatic functional annotation programs (Interproscan^52^, EggNOG^41^, nr BLASTp) and an amplification profile was assigned to each orthogroup by identifying the taxonomic group where the amplification of the family was most marked (Table S3).

In addition to the binomial tests, we also ran CAFE5^76^ to reconstruct the history of gene family amplifications. Such reconstructions rely on a species tree and require that all gene families are present at the root of the tree. However, of the 180 amplified OGs that were strongly amplified in Phaeophyceae (see above and listed in Table S3) only 19 were present at the ancestral node. The majority (161) of the 180 families were gained during the early evolution of the lineage, most (105) at the origin of the PX clade (i.e. a collapsed node corresponding to nodes n1 and n2 in Figure S4A) or of the Phaeophyceae/FDI clades (i.e. a collapsed node corresponding to nodes n5 and n6; Figure S4A). To determine whether the 180 amplified OGs were significantly enriched in genes that were gained early during Phaeophyceae evolution (i.e. at nodes n1/n2, n4, n5/n6 in Figure S4A), a Chi-squared test was carried out using the R *chisq.test* function on a contingency table containing the proportions of OGs gained at various periods during brown algal evolution for both the amplified OGs and for the entire set of OGs as a reference dataset (Table S1C). Twelve independent CAFE5 reconstructions were carried out on the OG subsets gained at 12 different nodes (n0, n1/2, n4, n5/n6, n8, n9, n10/n11, n13, n15, n18, n19, n20), using the subtrees rooted at these nodes so that the sets of OGs gained at each node would be placed at the root of the tree for one of the 12 analyses (Figure S5E). The analysis focused on the 19 highest quality genomes (Table S21), which is why some pairs of nodes were collapsed (e.g. nodes n1 and n2 to give n1/n2). Several parameters were tested for CAFE5: the –p option (Poisson distribution) resulted in better likelihood scores than default, but we observed a weak effect when increasing the value of lambda (–k). Consequently, all reconstructions were performed with –p (and no k, i.e. k=1) for efficiency purposes. As recommended by Mendes *et al*.^76^, very large gene families were discarded as these can cause the program to fail to initialize the parameters. The twelve reconstructions were then aggregated and the proportions of amplified and reduced gene families were calculated for each node (Table S6). Only results on internal nodes were considered, since leaves are more subject to artefactual amplifications/reductions due to genes being missed, fused or split in the annotations.

### Composite genes

The amino-acid sequences of all 530,598 genes present in the selected genomes were compared in an all-against-all pairwise alignment using DIAMOND BLASTp^26^ version 2.0.11; “very-sensitive” mode; e-value threshold 1e^-5^. This raw alignment was then filtered using CleanBlastp, from the CompositeSearch suite^40^, to remove sequence alignments with under 30% residue identity and produce the final sequence similarity network. CompositeSearch was then used on this network to identify putative composite gene families among the orthologous groups (OGs) previously computed by OrthoFinder^32^. Composite OGs containing two or more genes and having non-overlapping regions aligned to their component OGs were retained for further analysis, while singleton composite OGs and composites with overlapping component regions were discarded. A phylogeny-based approach^94^, which uses information from extant genomes to apply a Dollo parsimony model in Count^33^, was used to reconstruct the evolutionary events (domain fusions and fissions) that led to structural rearrangements of composite genes, allowing them to be labelled as fusion or fission events (or as complex events when sequentiality could not be clearly deduced).

### Horizontal gene transfer (HGT)

#### Dataset and experimental approach

Uneven data collection across taxa can impact HGT identification. The phylogeny-based HGT screening approach used here requires the establishment of a comprehensive and taxonomically diverse reference dataset. The analysis focused on the Phaeoexplorer genomes using a background database called REFAL and an automated bioinformatics tool called RoutineTree, which screens for HGTs using phylogenetics. The background database was built using a starting database, GNM1157, which includes a diverse set of 17,250,679 protein sequences from 1157 genomes spanning various prokaryotic and eukaryotic lineages (540 bacteria, 45 archaea, 431 Opisthokonta, 15 Rhodophyta, 83 Viridiplantae, and 43 genomes from CRASH lineages). Data from NCBI RefSeq (updated as of May 2020) and MMETSP were integrated into GNM1157 to form the background database REFAL. To enhance data quality and reduce redundancy, CD-HIT version 4.5.4 was used to remove highly similar sequences (with sequence identity ≥90%) within each taxonomic order. This curation process resulted in a protein database consisting of 39.9 million sequences, representing over 7,786 taxa and providing comprehensive coverage across the diverse branches of the tree of life. To obtain the best assembled genome within a genus, the latest version was selected if multiple versions were available. In addition, the dataset was expanded by searching for genomes in other repositories such as the Joint Genome Institute. Special attention was paid to achieving balanced representation of the Rhodophyta and Viridiplantae, which are particularly crucial for HGT analysis within the Chromalveolate group. To accomplish this, protein data from six red algal transcriptomes sourced from MMETSP was added. The HGT search was applied to 72 Stramenopile genomes, including 45 newly sequenced and 27 public genomes.

#### Phylogenetic Tree Reconstruction

The pipeline for constructing phylogenetic trees splits fasta files into individual sequence files and then carries out a search for homologous sequences, followed by multiple sequence alignment and tree-building. Nested positions within the trees were identified using artificial intelligence and hU and hBL methods were used for HGT verification. Instead of using all available sequences, sequences with the best BLAST hit scores from each kingdom, phylum, and class were used for tree construction to expedite tree-building and enhance clarity. Each gene, regardless of whether it was a copy or not, was used as a query for tree construction. To improve precision, four different methods were used for tree building: neighbour-joining, maximum parsimony, maximum likelihood and Bayesian. As a result, each node within a tree was associated with four support values. To create single-gene phylogenetic trees, a BLASTp^3^ search was carried out against the background database, employing an e-value cutoff of 1e^-05^. For each query, the top 1,000 significant matches were sorted by bit-score in descending order as the default criterion. Matching sequences were then retrieved from the database, with a constraint of no more than three sequences per genus and no more than 12 sequences per phylum. To further refine the selection, significant matches with a query-subject alignment length of at least 120 amino acids were re-sorted based on query-subject identity in descending order. A second set of homologous sequences was then retrieved from the database following the same procedure. These two sets of homologous sequences, along with the query, were merged and aligned using MUSCLE^34^ version 3.8.31 with default settings. The resulting alignments, trimmed to a minimum length of 50 amino acids using TrimAI^49^ version 1.2 in automated mode (-automated1), were used to construct phylogenetic trees with FastTree version 2.1.7, with the ‘WAG + CAT’ model and four rounds of minimum-evolution SPR moves (-spr 4) along with exhaustive ML nearest-neighbour interchanges (-mlacc 2 -slownni). Branch supports were estimated using the Shimodaira-Hasegawa (SH)-test.

#### Inferring HGT based on tree topology

Phylogenetic trees were examined to identify specific topologies where Phaeoexplorer query sequences were nested among other sequences, defined as a situation where two or more monophyletic clades consist of both queries and prokaryotic sequences, supported by distinct nodes within the tree. These monophyletic clades are considered to group together if they share the same set of prokaryotic sequences but differ in sequences from optional taxa. Singletons for both the donor and receptor genes were excluded to minimise contamination and recent HGT interference. To retain only robustly supported nested positions, positions were required to be multiply supported, with a minimum of ≥0.70 for the SH-test and aByes-test support from at least two Phaeoexplorer receptor nodes and three donor supporting nodes. Furthermore, queries that displayed significantly different amino acid compositions (P < 0.05) compared to the remaining sequences in the alignment were discarded. Queries from the CRASH category that nested among sequences from other kingdoms (supported by >70% UFBoot at one or more supporting nodes) were retained.

#### Enhancing accuracy and establishing the timing of HGTs

To enhance accuracy, a minimum requirement was imposed for all supporting nodes and for strongly supported nodes that indicate query-donor monophyly. To determine the timing of HGT events, temporal information, primarily derived from the timetree database, was incorporated into each node. We assigned the “smallest boundary” role to pinpoint the most recent common ancestor at the time of the HGT event. Essentially, if all descendants of a given query protein sequence can be traced back to the initial HGT event, a common ancestral node can be identified whose occurrence time can be inferred using a molecular clock approach based on archaeological and fossil evidence. The taxonomy boundaries of HGT descendants were determined by identifying the smallest ancestor shared by both the donor and receptor taxa from the monophyletic clades within the tree. By considering the emergence times of both taxa, the timing of the transfer of genes from earlier taxa to later taxa can be determined, as the reverse scenario is not considered plausible.

#### Verification of HGTs

Verification of HGT used the following contamination assessment criteria: i) HGT candidates were excluded if they were located in a contig where 50% of the genes had better matches with other kingdoms, ii) HGT candidates were excluded if they were located in a contig where 50% of the genes were primarily identified as HGT genes, iii) HGT candidates were excluded if one of their five closest flanking genes, both upstream and downstream, had a better match with other kingdoms. AI, hU and the hBL value were used to further validate HGT events. This process was supplemented with annotation and functional predictions for the identified HGTs.

Further validation was based on the following concepts:

*OUTGROUP*: This comprises all biological donors present in a tree, excluding the query species if it belongs to biological donors.

*SKIP*: This includes all biological receptors (species belonging to optional taxa) in a tree, again excluding the query species if it belongs to biological receptors.

*INGROUP*: This encompasses species from SKIP’s upper level, excluding SKIP itself and the query species (if it belongs to biological receptors).

*AI (Alien Index)*: computed for each query gene using e-values from BLAST hits:

*AI = (E-value of best BLAST hit in the INGROUP lineage) / (E-value of best BLAST hit in the OUTGROUP lineage)*

The AI score quantifies how similar queries are to their homologs in the OUTGROUP compared to homologs in the INGROUP. We apply a relatively lenient cut-off (AI > 0) for initial screening, which can be adjusted in the second screening as needed.

*hU (HGT Score Support Index)*: calculated for each query gene based on the best bit scores of INGROUP vs. OUTGROUP:

*hU = (Best-hit bitscore of OUTGROUP) - (Best-hit bitscore of INGROUP)*

A lenient cut-off (hU > 0) is used for initial screening, with flexibility for adjustment in the second screening.

*hBL (HGT Branch Length Support Index)*: calculated based on the minimum branch length to the query within INGROUP vs. OUTGROUP:

*hBL = (Minimum branch length to the query within INGROUP) - (Minimum branch length to the query within OUTGROUP)*

A lenient cut-off (hBL > 0) is applied initially, with the option for modification in the second screening. *CHE, CHS, CHBL (Consensus Hit Support)*: To mitigate the possibility that the best bit score for either INGROUP or OUTGROUP is influenced by contamination, we consider alternative matches. We introduce consensus hit support (CHE, CHS, and CHBL) to assess the reliability of AI, hU, and hBL, respectively.

For example, if AI > 0, CHE evaluates the likelihood that “AI remains greater than 0” when using the e-value of each sequence in OUTGROUP instead of bbhO. A similar approach applies to CHS for hU and CHBL for hBL. This additional layer of evaluation helps ensure the robustness of the HGT verification process.

#### Gene codon usage, functional annotation and expression

Indices of codon usage and GC content were calculated using Codonw 1.4.4 (http://codonw.sourceforge.net). Gene functions were assigned by searching against the Gene Ontology (GO) database using blast2GO (ref blast2GO 08) and the KEGG database using blastKOALA (http://www.kegg.jp/blastkoala/) with default parameters. The full gene sets of each species were set as the background for KEGG and GO enrichment analyses by applying Student’s t-test (*p*-value cutoff = 0.01). HGTs were also analysed with SEED (http://www.theseed.org/wiki/Home_of_the_SEED), IPR2GO (http://www.ebi.ac.uk/interpro/search/sequence-search), eggNOG^41^ (http://eggnogdb.embl.de/#/app/home) and Pfam^82^. For each species, the differences between mean gene expression levels for HGTs and non-HGT genes with common GO terms were accessed using Student’s t-test. Go terms with less than five genes in either gene category were ignored. The differences in expression dispersal (coefficient of variation: standard deviation across genes or samples / mean value) and expression specificity (frequencies of a gene to be detected as unexpressed, defined as TPM = 2, in any condition) were accessed in a similar manner. Given the variable experimental conditions associated with different transcriptome data for each species, gene expression values for a gene were used indiscriminately regardless of the conditions. Correlation tests between the codon adaptation index (CAI) and gene expression were carried out using the Spearman’s rank correlation analysis tool (P. Wessa, Free Statistics Software, Office for Research Development and Education, version 1.1.23-r7, https://www.wessa.net/).

### Comparative analysis of gene sets identified by genome-wide analyses of evolutionary history

Genes identified as belonging to orthogroups that were predicted to be gained at specific nodes of the phylogenetic tree based on the Dollo parsimony analysis, to belong to either significantly amplified gene families (binomial analysis) or to belong to gene families that have significantly changed in size over evolutionary time (CAFE5 analysis), to correspond to founder events (Phylostratigraphy analysis), to have been remodelled (composite gene analysis) or to have been derived from an HGT (HGT analysis) were extracted from the output of each of these analysis and aggregated in a single datatable. Correspondences were established manually between phylogenetic tree nodes and phylostrata and this information was integrated into the datatable. Counting and calculations of the frequency of events at specific time points were carried out using *ad hoc* R scripts (R version 4.4.1) and the tidyverse package (version 2.0.0). Graphs were generated using the ggplot2 package (version 3.5.1). For each gene, a COG functional category was retrieved from the eggNOG mapper output and the COG enrichment analysis was carried out in R using the clusterProfiler package (version 4.6.2) by comparing each set of gene families with the full set of gene families.

### Detection of viral genome insertions and viral regions in algal genomes

To reduce the dataset size for analysis, 64 Phaeoexplorer and eight public genomes were initially filtered to retain only contigs that were more than 10 kbp in length. Gene prediction was then carried out on all contigs using Prodigal^43^ (V2.6.3, settings: default, meta) and the resulting proteins were used as queries against the NCVOG^79^ and VOGDB^95^ databases using hmmscan (HMMER 3.3.2 with default settings). The contigs detected by hmmscan were then filtered to retain only sequences with at least one match to either viral database at a defined e-value cutoff (1e^-20^ for NCVOG, and 1e^-80^ for VOGDB). The resulting positive 4,951 contigs were then analysed using ViralRecall^44^ version 2.0 with settings -w 50 -g 1 -b -f -m 2 using the built-in Nucleocytoviricota (NCV) database GVOG and a window size of 50 kbp. To ensure that viral genes were not missed because they had not been annotated by Prodigal, six-frame translations of the contigs were generated using esl-translate (version 0.48 with default settings), and the resulting proteins queried against the same databases used by ViralRecall using hmmsearch (HMMER 3.3.2, settings: -E 1e-10). The ViralRecall results were then parsed using an in-house workflow. Six-frame translations were removed from the results if they overlapped (even partially) with any Prodigal gene prediction, as identified using bedtools^45^ (v2.29.2; intersect). Likewise, overlapping six-frame translations and gene predictions with the same NCVOG match were removed to reduce redundancy. Based on the distance between query sequences with the same GVOG hit, queries were flagged as frame-shifted (less than 100 bp gap), intron-containing (100-5,000 bp gap) or mono-exonic (greater than 5,000 bp gap). All queries were also checked for overlaps with multi-exonic genes that had been annotated by the Phaeoexplorer gene prediction procedure (using Gmove^28^), and flagged if they did. All queries were then filtered to retain only those that matched a set of key NCV marker genes, identified by NCVOG code (A32, D5 helicase, D5 DNA primase, MCP, DNA polymerase B, SFII and VLTF3) or some Phaeovirus integrase genes (integrase recombinase, integrase resolvase and RNR). The marker gene proteins were clustered with the protein sequences of NCVOGs using MMseqs cluster^46^ (version 13.45111 with settings --min-seq-id 0.3 -c 0.8). Finally, the parsed results of the NCV marker gene set identified by the ViralRecall screen were manually curated, retaining only those queries with varying combinations of the following properties: placement within a viral region as identified by ViralRecall, similar hmmsearch results (score and e-value) and gene length to that of known NCV genes, not part of a multi-exonic gene, lack of Pfam HMM matches to cellular domains sharing homology to the marker gene (specific to certain marker genes), and clustered with an NCVOG in the MMseqs analysis. We noted that the median number of viral regions found in genomes assembled with long reads was very similar to that for genomes assembled with short reads (9 and 10, respectively). The marker gene content of the viral regions was manually assessed to estimate the number of complete or partial inserted viruses in each genome. VRs were considered to be complete proviruses if they contained all seven of the key NCV marker genes listed above. VRs were classed as partial proviruses if they only contained a subset of the seven key NCV marker genes, the presence of the MCP and DNA polymerase B genes being particularly strong indicators of a partial provirus.

To classify genes in VRs (Figure 6B), viral sequences were removed for the NBCI RefSeq non-redundant protein database (NR) by removing proteins assigned to the “Viruses” category and by comparing the database with RVDB using BLASTp and removing any proteins that matched with an e-value cut-off of < 1-e40 to create a “virus-free NR” database. Deduced proteins were then compared with the RVDB and the virus-free NR databases using BLASTp and relative bitscores (rbitscores) were calculated by dividing the BLASTp bitscore for the best match in each database by the query protein’s self-hit bitscore^96^. Self-hit scores were acquired by comparing the complete deduced proteomes with themselves using BLASTp. Proteins with a RVDB rbitscore at least 20% greater than its virus-free NR rbitscore were designated as “viral”. Proteins with a virus-free NR rbitscore at least 20% greater than its RVDB rbitscore were designated as “cellular” (i.e. corresponding to a gene from a cellular organism). Ambiguous cases without a 20% differential were designated as “viral or cellular” and proteins with no significant matches were designated as ORFans (i.e. unknown proteins).

The presence of host regions flanking the viral regions was evaluated based on the ViralRecall output (Table S19C). The percentages of viral regions with two, one or zero flanking regions (longer than 2 kbp) were 25.8%, 15.0% and 59.2%, respectively (i.e. 40.8% of viral regions had at least one flanking region). Of the viral regions that had two flanking regions, 89.5%, 7.0% and 3.5% had flanking regions with a total length of >200 kbp, between 20 and 200 kbp or between 2 and 20 kbp, respectively. For the viral regions that had one flanking region, the corresponding percentages were 25.3%, 36.7% and 38.0%.

### Phylogenetic analysis of viral genes

Amino acid sequences of manually-curated collections of major capsid protein (MCP) and DNA polymerase B proteins were aligned using MAFFT (v7.520, settings: --adjustdirectionaccurately –auto --maxiterate 1000) and phylogenetic trees were generated using IQ-TREE (v 2.2.2.3, settings: -m MFP -B 1000).

### Metabolic networks

Genome-scale metabolic networks were reconstructed using AuCoMe^97^ version 0.5.1 using the MetaCyc^98^ version 26 database. A first dataset, consisting of the 60 species listed in Table S21 (column “Metabolic networks”) plus two public diatom genomes already used in the initial AuCoMe study (*Fragilariopsis cylindrus* and *Fistulifera solaris*) was processed to build the largest possible database (phaeogem) for exploratory comparisons (https://gem-aureme.genouest.org/phaeogem/). Then, a second comparison was performed on all long-read species plus outgroups. Based on Multidimensional-scaling (MDS) analyses, the most divergent long-read species (*Choristocarpus tenellus*, *Laminaria digitata, Phaeothamnion wetherbeei* and the public genome of *Sargassum fusiforme*) were excluded to construct a 16 species dataset, balancing assembly quality and phylogenetic coverage (https://gem-aureme.genouest.org/16bestgem/). The MDS plots shown in Fig S7 were build using the vegan package, version 2.6-4 (https://github.com/vegandevs/vegan) with R 4.1.2^87^, using Jaccard distances. A third stricter dataset (fwgem), enriched in high-quality long-read Ectocarpales, was built to address questions related to freshwater adaptation (https://gem-aureme.genouest.org/fwgem/). A set of reactions that were overrepresented in brown algae compared to the outgroup was created (Table S4) by taking reactions present in 100% of brown algae and less than 70% of outgroups. Reactions corresponding to genes lost in freshwater species were also extracted (Table S22). These reaction sets were extracted from all the networks using the Aucomana library (https://github.com/PaulineGHG/aucomana). Online wikis (phaeogem, 16bestgem and fwgem) were generated using AuReMe^99^.

### CAZymes

CAZyme genes were identified based on shared homology with biochemically characterised proteins, either individually or as hidden Markov model (HMM) profiles. For phylogenetic analyses, proteins were aligned using MAFFT^47^ with the iterative refinement method and the scoring matrix Blosum62. The alignments were manually refined and trees were constructed using the maximum likelihood approach. Alignment reliability was tested by a bootstrap analysis using 100 resamplings of the dataset. Only bootstrap values above 60% are shown. The phylogenetic trees were displayed with MEGA^48^. The annotated genes are listed in Table S6 with accession numbers.

### Sulfatases

The sulfatases encoded by each brown algal genome were identified and assigned to their respective family and subfamily using the SulfAtlas database^80,81^ (https://sulfatlas.sb-roscoff.fr/). Each predicted proteome was first submitted to the SulfAtlas HHM server (https://sulfatlas.sb-roscoff.fr/sulfatlashmm/), which allows rapid identification of sulfatase candidates and (sub)family assignment using hidden Markov model profiles for each SulfAtlas (sub)family. Each sulfatase candidate sequence was then used as a query in a BLASTp^3^ search against the SulfAtlas database (https://blast.sb-roscoff.fr/sulfatlas/). Sequences with at least 50% identity with sulfatases from marine bacteria or other marine microorganisms were considered to be contaminants. Below this threshold, additional examination of the predicted gene structure and genomic context of the candidate sequence was undertaken to identify possible horizontal gene transfers.

### Haloperoxidases

vHPO genes were identified based on sequence homology and active site conservation. Maximum likelihood phylogenetic analyses were carried out using the NGphylogeny platform at https://ngphylogeny.fr/. MAFFT was used to align vHPO sequences and alignments were automatically curated with TrimAl^49^, leading to the selection of 444 informative positions from the initial 1450 positions for the algal-type vHPOs and 402 informative positions from the initial 1078 positions for the bacterial-type vHPOs. Maximum likelihood trees were constructed using FastTree with the WAG+G gene model and 1000 bootstrap replicates. Maximum likelihood Newick files were formatted as circular representations using iTOL. Only bootstrap values between 0.7 and 1 were conserved. The lists of annotated vHPO genes are in Tables S7 and S8.

### Ion channels

A search was carried out for 12 classes of ion channel in the predicted proteomes of the 21 Phaeoexplorer reference genomes plus those of two diatoms, *Phaeodactylum tricornutum* and *Thalassiosira pseudonana* (Table S16). Predicted proteomes were screened using BLASTp^3^ and query sequences from *Ectocarpus* species 7 and seven other species from diverse eukaryotic taxa (Table S16).

### Membrane-localised proteins

Membrane protein family genes were identified either by carrying out BLASTp^3^ searches of the predicted Phaeoexplorer proteomes using *Ectocarpus* species 7 sequences as queries or by recovering orthogroups containing the relevant *Ectocarpus* species 7 sequences as members. The BLASTp approach was used for DEK1-like calpains, fasciclins, tetraspanins, CHASE, ethylene-binding-domain-like and MASE1 domain histidine kinases whereas the orthogroup approach was used to recover other members of the histidine kinase family. Both approaches were used to search for integrins and transmembrane receptor kinases. For integrins the two methods detected exactly the same set of proteins. For receptor kinases the BLASTp and orthogroup analyses detected 99.3% and 98.3% of the 269 genes, respectively. For these analyses, either the whole genome dataset was analysed or only the set of 21 reference genomes (Table S21), depending on the size of the gene family.

Manually-curated histidine kinase protein families were aligned with Muscle^34^ before phylogenetic tree construction using IQ-TREE 2^75^ (version 2.3.4) with automatic model selection and 1000 bootstraps.

### Transcription-associated proteins

TAPscan v4^51^ was used to analyse the transcription-associated protein (TAP) complements of 21 species. TAPscan^50^ is a comprehensive tool for annotating TAPs based on the detection of highly conserved protein domains using HMM profiles with specific thresholds and coverage cut-offs. Following detection, specialised rules are applied to assign protein sequences to TAP families based on the detected domains. TAPscan v4 can assign proteins to 138 different TAP (sub)families with high accuracy.

### EsV-1-7 domain proteins

EsV-1-7 domain proteins were identified in the 31 brown algal and sister taxa genomes (Table S21) by recovering the members of all orthogroups (with the exception of OG0000001, which is a very large OG that consisting principally of transposon sequences) that either contained one or more of a curated set of 101 EsV-1-7 domain proteins^100^ for *Ectocarpus* species 7 or contained an EsV-1-7 domain protein based on a match to the Pfam EsV-1-7 motif PF19114. The recovered proteins were screened manually for the presence of at least one EsV-1-7 domain and a total of 2018 were finally identified as members of the EsV-1-7 family.

To identify orthologues of the EsV-1-7 protein IMMEDIATE UPRIGHT^100^ (IMM), BLASTp searches of 25 brown algal and four sister taxa predicted proteomes were carried out with the amino-terminal domain of the IMM protein minus the five EsV-1-7 repeats as this domain is unique to IMM. Proteins were retained as IMM orthologues if they were more similar to IMM than to the most closely-related protein in *Ectocarpus* species 7, Ec-17_002150.

### Histones

Histone protein sequences were analysed in *Ascophyllum nodosum*, *Chordaria linearis*, *Chrysoparadoxa australica*, *Desmarestia herbacea*, *Dictyota dichotoma*, *Discosporangium mesarthrocarpum*, *Ectocarpus crouaniorum*, *Ectocarpus fasciculatus*, *Ectocarpus siliculosus*, *Fucus serratus*, *Heterosigma akashiwo, Pleurocladia lacustris*, *Porterinema fluviatile*, *Pylaiella littoralis*, *Saccharina latissima*, *Sargassum fusiform*, *Schizocladia ischiensis*, *Scytosiphon promiscuus*, *Sphacelaria rigidula*, *Tribonema minus* and *Undaria pinnatifida* using BLASTp against the complete predicted proteomes (https://blast.sb-roscoff.fr/phaeoexplorer/) with the histone protein sequences from the diatom *Phaeodactylum tricornutum* as queries. The genes and transcripts coding for the identified histones were then retrieved from the genomes and predicted transcripts using BLAST (https://blast.sb-roscoff.fr/phaeoexplorer/). The proteins encoded by the identified genes and transcripts were predicted with the Expasy web translator (https://web.expasy.org/translate/). All the identified histone protein sequences and the corresponding transcripts IDs are reported in Table S23. In order to identify truncated proteins or incorrect start codons, the following constraints were applied: H2A proteins must start with the SGKGKGGR sequence, H2B with AKTP, canonical H3.1 and variants H3.3 with ARTKQT and H4 with SGRGKGGKGLGKGG. For the linker histone H1, protein sequences had to be lysine-rich and sequences with incorrect start codons were determined by alignments of all identified H1 proteins. For proteins with incorrect start codons, the region upstream of the correct start codon was removed. For truncated proteins, *i.e.* proteins whose transcripts lacked either the start (no methionine) or stop codons, the protein sequence was completed based on alignment with the corresponding genomic region using the Geneious 11.0.5 software. When the sequence could not be completed, a BLAST was performed against the Phaeoexplorer *de novo* transcriptomes (https://blast.sb-roscoff.fr/phaeoexplorer_denovo/) when this data was available (this was not possible for the public genomes *T. minus*, *U. pinnatifida* and *S. fusiforme*). Based on the nomenclature established by^101^, H3 histones were classified as follows: canonical H3.1 proteins harbour AT residues at positions 31-32 while histone variants H3.3 harbour TA residues, H3 proteins with other residues at positions 31-32 were named H3.4 and so on. CenH3 variants of H3 were identified by analysis with Panther 17.0 (www.pantherdb.org/tools/sequenceSearchForm.jsp?) and/or Interproscan^52^ 94.0 (www.ebi.ac.uk/interpro/search/sequence/).

Species abbreviations used in histone phylogenetic trees are: Atr, *Amborella trichopoda*; At, *Arabidopsis thaliana*; Ce, *Caenorhabditis elegans*; Di, *Dictyostellium discoideum*; Dr, *Danio rerio*; Dm, *Drosophila melanogaster*; Hs, *Homo sapiens*; Mm, *Mus musculus*; Pp, *Physarum polycephalum*; Ppa *Physcomitrium patens*; Sc, *Saccharomyces cerevisiae*; Tm, *Tetrahymena thermophila*; Zm, *Zea mays*; Mp, *Marchantia polymorpha* subsp. *Ruderalis*; Bd, *Brachypodium distachyon*; Ccr, *Chondrus crispus*; Gs, *Galdieria sulphuraria*; Cm, *Cyanidioschyzon merolae*; Cr, *Chlamydomonas reinhardtii*; Ol, *Ostreococcus luciminarinus*; Ot, *Ostreococcus tauri*; To, Thalassiosira oceanica; Pt, *Phaeodactylum tricornutum*; An, *Ascophyllum nodosum*; Cl, *Chordaria linearis*; Ca, *Chrysoparadoxa australica*; Dh, *Desmarestia herbacea*; Ddi, *Dictyota dichotoma*; Dme, *Discosporangium mesarthrocarpum*; Ec, *Ectocarpus crouaniorum*; Ef, *Ectocarpus fasciculatus*; Es, *Ectocarpus siliculosus*; Fse, *Fucus serratus*; Ha, *Heterosigma akashiwo*; Pla, *Pleurocladia lacustris*; Pf, *Porterinema fluviatile*; Pli, *Pylaiella littoralis*; Sl, *Saccharina latissima*; Sf, *Sargassum fusiform*; Si, *Schizocladia ischiensis*; Sp, *Scytosiphon promiscuus*; Sri, *Sphacelaria rigidula*; Tm, *Tribonema minus*; Up, *Undaria pinnatifida*.

### DNA methyltransferases

Searches were carried out for methyltransferases and demethylases in the predicted proteomes of 20 of the high quality brown algal reference genome assemblies (based on Nanopore long-read sequence) plus the sister taxa *Chrysoparadoxa australica* and *Schizocladia ischiensis* using BLASTp (Table S21). A methyltransferase reference database was constructed by recovering sequences from NCBI, ENSEMBL and UniProtKB. Methyltransferase sequences were recovered for stramenopiles such as *Nannochloropsis gaditana*, the diatom *Phaeodactylum tricornutum*, the oomycete *Phytophthora infestans* and for species from more distant lineages including *Arabidopsis thaliana*, *Homo sapiens* and the fungus *Neurospora crassa*. The proteomes of the selected brown algal strains were then queried against this database using BLASTp and matches with an e-value of < 0.001, a bit score > 70, a maximum gap of 5 and percentage identity of >30% were retained. The retained matches were screened against the NCBI, UniProt and SwissProt databases to identify and remove contaminating bacterial or viral proteins. Methyltransferase domains were detected in the retained matches using the Simple Modular Architecture Research Tool (SMART)^83^ domain architecture analysis and InterPro searches (https://www.ebi.ac.uk/interpro/). Sequences with methyltransferase domains were retained for further analysis. Validated brown algal methyltransferases were aligned with reference methyltransferases using Clustal^53^ 2.1.

### Spliceosome

Components of the Major Spliceosome were identified using a reference set of 147 human components (https://www.genenames.org/data/genegroup/#!/group/1518), excluding the five small nuclear RNAs (snRNAs). Including isoforms, this query set consisted of 626 proteins. These proteins were used to screen the predicted proteomes of 54 genomes (Table S21) using BLASTp and matches were retained if they had an e-value of at most 1e^-40^ and coverage >30%. Searches were also carried out for components of LSm and Sm complexes which have roles as scaffolds in the formation of ribonucleoprotein particles (RNPs), in the maturation of mRNAs (including splicing, such as the cytoplasmic complex LSm1-7, LSm2-8 which is part of the core U6 snRNP and other complexes important for the formation of the 3’ ends of histone transcripts), in the assembly of P-Bodies and in the maintenance of telomeres.

### Flagella proteins

A previous proteomic analysis of anterior and posterior flagella of the brown alga *Colpomenia bullosa* identified a total of 592 proteins across the two proteomes^102^. Here the *Ectocarpus* species 7 orthologues of 70 of these proteins that had been detected with a very high level of confidence were used to identify the corresponding orthogroups and the presence or absence of these orthogroups was scored for seven representative species (Table S21).

### Detection of *Porterinema fluviatile* genes differentially expressed in freshwater and seawater

Six independent cultures of *Porterinema fluviatile* were cultivated for four weeks in 140 mm Petri dishes with Provasoli-enriched culture medium^103^, which was renewed every two weeks. For three Petri dishes, the culture medium was based on autoclaved natural seawater (high salinity treatment), for the other three Petri Dishes natural seawater was diluted 1:19 vol/vol with distilled water (low salinity treatment). Cultures were harvested with 40 µm nylon sieves, dried with a paper towel, and immediately frozen in liquid nitrogen. RNA extraction library construction and sequencing were carried out as described in section “RNA extraction, Illumina RNA-seq library preparation and sequencing”. RNA-seq reads were cleaned with Trimmomatic^18^ V0.38 and then mapped to the *P. fluviatile* genome using Kallisto^55^ version 0.44.0. Differentially expressed genes were identified using the Deseq2 package^56^ included in Bioconductor version 3.11, considering genes with an adjusted p < 0.05 and a log2 fold-change > 1 as differentially expressed. The complete list of differentially expressed genes is provided in Table S24. To compare the differentially expressed genes in *P. fluviatile* with an equivalent set previously identified for *Ectocarpus subulatus* in a microarray experiment using nearly identical growth conditions^104^, orthologues in the two species were detected using Orthofinder version 2.3.3. Of the 10,066 shared orthogroups, 6,606 had microarray expression data for *E. subulatus*. This information was used to classify differentially expressed genes for the two species as either shared orthologues or as lineage-specific.

### Identification of genes with generation-biased expression patterns

RNA-seq data (two to five replicates per condition) was recovered for gametophyte and sporophyte generations of ten species (Table S21). Data quality was assessed with FastQC^57^ version 0.11.9 and sequences were then trimmed with Trim Galore version 0.6.5 with the parameters --length 50, - quality 24, --stringency 6, --max_n 3. The cleaned reads were mapped onto the corresponding genome for each species using HISAT2 version 2.1.0 with default options. Counting was carried out with featureCounts^58^ from the subread package (version 2.0.1) on CDS features grouped by Parent. Transcript Per Kilobase Million (TPM) tables were generated for all conditions and differentially expressed genes were detected using DESeq2^56^ version 1.30.1. Genes were classified into six categories based on the differential expression analysis and the TPM values: gametophyte-biased, mean TPM ≥1 in gametophyte and sporophyte, log2(fold change) ≥1, adjusted *p*-value <0.05; sporophyte-biased: mean TPM ≥1 in gametophyte and sporophyte, log2(fold change) ≤-1, adjusted *p*-value <0.05; gametophyte-specific, mean TPM <1 in sporophyte and ≥1 in gametophyte, log2(fold change) ≥1, adjusted *p*-value <0.05; sporophyte-specific, mean TPM <1 in sporophyte and ≥1 in gametophyte, log2(fold change) ≤-1, adjusted p-value <0.05; unbiased genes: mean gametophyte and sporophyte TPMs ≥1, log2(fold change) <1 or >-1 and/or adjusted p-value ≥0.05; unexpressed genes, mean gametophyte and sporophyte TPM <1.

### Life cycle and thallus architecture

#### Genome dataset and traits

To study the impact of body architecture, the brown algae were divided into three categories: 22 filamentous species, eight simple parenchymatous species and 13 species with elaborate thalli (Table S21). For the life-cycle-based assessment, the groups were: 30 haploid-diploid species and six diploid species (Table S21). Body architecture information was available for 43 species, and life cycle information was available for 36 species; species without body plan or life cycle information were not used in subsequent analyses. Two approaches were used to estimate selection intensity across the phylogeny, (i) a model-based method, and (ii) by evaluating codon usage bias and nucleotide composition. Two evolutionary models were used, one based on architecture and the other based on life cycle. For model-based methods the phylogeny was categorised based on the above traits, and selection intensity parameters were estimated using PAML^59^ version 4.9i. Rate estimates were obtained for non-synonymous substitutions (dN), synonymous substitutions (dS) and omega (dN/dS) for the multiple sequence alignments of all genes within each orthogroup using the variable-ratio model of CODEML from PAML, which allows different omegas for different branch categories. The traits were assigned to the branches of the phylogeny using ancestral state estimation by stochastic mapping with the phytools R package^60,87^.

#### Evolutionary models to study impacts of body architecture

To study variation in selection intensity as a function of body architecture, we devised a model with the following trait categories: filamentous/pseudoparenchymatous (simple cell division and organisation on a single plane), parenchymatous (cell division and organisation on multiple planes) and elaborate thallus (tissue differentiation). To ensure that at least 50% of the species in each category were used in the analysis, we selected orthogroups (OGs) that contained at least 11 members for filamentous, at least four members for parenchymatous and at least six members for elaborate thallus algae. Using this filter, 1068 OGs were obtained, on which the model based on body architecture was fitted. Selection intensity parameters [rate of non-synonymous substitution (dN), rate of synonymous substitution (dS) and omega (dN/dS)] were estimated for the three trait categories for each gene alignment. We used the Wilcoxon signed-rank test to evaluate the statistical significance of differences between the selection intensity parameters (dN, dS and dN/dS) for each category.

#### Evolutionary models to study the impacts of life cycle

The impact of life cycle on molecular evolution was assessed using a model with two categories consisting of diplontic and haplodiplontic species. For this model we used 1,058 OGs that contained at least three members for diploid species and at least 15 members for haploid-diploid species. Using alignments of the gene within the OGs, we estimated the selection intensity parameters for the different categories and applied the Wilcoxon signed-rank test to assess the statistical significance of differences in selection intensity between the diploid and haploid-diploid life cycles.

#### Selection of intensity parameters

Omega (dN/dS) provides an estimate of the ratio of substitutions at sites under selection compared to neutral sites, and is generally used to infer the strength of purifying selection. Omega needs to be interpreted with caution because not all synonymous sites are neutral^105^ and also synonymous substitutions are often underestimated due to saturation of synonymous sites, which might in turn impact the omega ratios^106^. Omega values lower than one indicate substitutions are less frequent at sites under selection compared to neutral sites and are characteristic of highly conserved genes or genes evolving under strong purifying selection. As we used primarily low copy number genes in this study, the analysed genes were expected to evolve under strong purifying selection, with omega values much lower than one. Using omega for near neutral studies is challenging because near neutral sites are determined by effective population size, that is to say, sites under mild selection constraint in larger populations can behave as neutral sites in smaller populations. It is therefore difficult to infer the amount of mutation from relative values of omega. In order to obtain better insight into selection intensity, mutation accumulation was not only investigated using rates of synonymous (dS) and non-synonymous (dN) substitutions but also by estimating codon bias and nucleotide composition. Codon usage bias was used, in addition to omega, to infer selection intensity across species as the former reflects selection efficacy at synonymous sites^107–109^. We inferred codon usage bias by estimating the effective number of codons (ENC) for each species using the enc method from the VHICA package^61,89^. The effective number of codons (ENC) quantifies the extent of deviation of codon usage of a gene from equal usage of synonymous codons. For the standard genetic code, ENC values range from 20 (where a single codon is used per amino acid implying strong codon usage bias) to 61 (implies that all synonymous codons are equally used for each amino acid^110^). Low ENC indicates constrained use of codons, which potentially highlights stronger codon bias due to stronger selection at synonymous sites. As nucleotide composition can also influence codon bias, we calculated the overall GC composition, GC at the third codon position (GC3) and the theoretical expected ENC (EENC) based on GC3 using local R scripts. The lower the observed ENC (OENC, estimated from the gene sequence) relative to EENC, the stronger the influence of selection due to translation on codon usage. This was studied by estimating the difference (DENC = EENC - OENC) between the expected ENC and the observed ENC^111^. Positive DENC indicates a role for selection constraints on codon usage in addition to the influence of nucleotide composition. DENC values of zero or less indicate that codon bias is entirely driven by nucleotide composition. DENC values were used to study the influence of translation selection and nucleotide composition on codon usage bias.

### Assembly and analysis of organellar genomes

Plastid and mitochondrial genomes were assembled *de novo* using NOVOPlasty^62^ v3.7 and *rbcL* and *cox1* nucleotide sequences as seeds. Assembled genomes were checked by aligning reads using Bowtie2^5^ v2.3.5.1 and processed with SAMtools^63^ v1.5. Annotation of protein-coding genes was performed with GeSeq^64^ v2.03. Annotation of tRNAs, tmRNAs and rRNAs was performed with ARAGORN^65^ v1.2.38.

Maximum-likelihood (ML) phylogenetic trees were constructed using 92 plastid genomes (11 non-brown outgroup sequences) and 89 mitochondrial genomes (seven non-brown outgroup sequences). The conserved coding-region amino acid sequences of 139 plastid genes (31,159 amino acids) and 35 mitochondrial genes (7,461 amino acids) were used to construct these phylogenetic trees. The sequence for each gene was aligned individually using MAFFT^47^ v7 (--maxiterate 1000) and then concatenated. Alignment partitions were assigned based on genes. Each of the aligned gene sequences was trimmed with trimAl^49^ v1.2 (-automated1). ML phylogenetic trees were constructed with IQ-TREE 2^75^. The protein substitution models in each gene partition were selected using ModelFinder^66^. Statistical support for tree branches was assessed with 1,000 replicates of ultrafast bootstrap (UFBoot2)^67^.

### Analysis of *Ectocarpus* genome synteny

Global genome synteny analysis was performed using SynMap^68^ on the CoGe platform (https://genomevolution.org/coge/) with the following genomes: *Ectocarpus crouaniorum* male, *Ectocarpus fasciculatus* male, *Ectocarpus siliculosus* male, *Ectocarpus* species 7 male and *Ectocarpus subulatus*. SynMap identifies syntenic regions between two or more genomes using a combination of sequence similarity and collinearity algorithms. Last^112^ was used as the BLAST algorithm and syntenic gene pairs were identified using DAGChainer^69^ with settings “Relative Gene Order”, -D = 20, -A =5. Neighbouring syntenic blocks were merged into larger blocks. Substitution rates between the synthetic CDS pairs were calculated using CodeML^59^, which was also implemented in SynMap, CoGe. In detail, protein sequences were aligned using the Needleman-Wunsch algorithm implemented in nwalign (https://pypi.org/project/nwalign/) and then translated back to aligned codons. CodeML was run five times for each alignment using the default parameters and the lowest dS was retained, with the upper cutoff for dS values set at 2. *Ectocarpus* genes were grouped according to their age based on the phylostratigraphic analysis and by chromosomal location based on their chromosome position in *Ectocarpus* species 7. All plots and statistical analysis were carried out in R version v.4.3.1. Local synteny analysis was based on orthologous genes as identified by Orthofinder.

### Analysis of *Ectocarpus* gene evolution

Protein sequence alignments were used to remove gaps with trimAl^49^ and then translated back to DNA with backtranseq^113^. Only DNA fasta files with a minimum of 70 bp were retained (831 single-copy orthologs). PhyML trees were built with Geneious v11.1.5 (https://www.geneious.com). Maximum likelihood analysis was carried out to detect site specific, branch-site specific and branch specific positive selection as well as sites under negative selection, using PAML^114^.

### Phylogenetic analysis of *Ectocarpus* species

Phylogenetic analysis was carried out for 11 *Ectocarpus* species plus *Scytosiphon promiscuus* as an outgroup (Table S21). Of the 933 single-copy orthogroups identified for these 12 species, 261 high-confidence alignments were retained for gene tree and species tree inferences following the removal of low-quality alignments using BMGE^115^. Bayesian inference of the phylogeny of the *Ectocarpus* species complex was performed using BEAST^70^ v2.7. The analysis was conducted under the multi-species coalescent (MSC) model, implemented in StarBEAST3^71^ v1.1.7. The MSC model coestimates gene trees and the species tree within a multispecies coalescent framework, enabling the assessment of incongruences among genes with respect to the species tree. To account for substitution model uncertainty, bModelTest^72^ was employed to average over a set of substitution models for each alignment. StarBEAST3 was run under both the Yule model and the strict clock model. A total of 300,000,000 Markov Chain Monte Carlo (MCMC) generations were conducted, with tree states stored every 50,000 iterations. Posterior tree samples were combined, discarding the initial 10% burn-in, using LogCombiner v2.4.7. A maximum clade credibility tree was generated using TreeAnnotator^70^ v2.4.7.

### *Ectocarpus* introgression analysis

To distinguish introgression from shared ancestry, D estimates (i.e. ABBA-BABA tests) were generated from 36 four-taxon combinations^116^: four to test the level of introgression within clade 1 (i.e. *E. subulatus*, *E. crouaniorum*, *Ectocarpus* species 1, *Ectocarpus* species 2), 20 to test the level of introgression within clade 2 (i.e. *Ectocarpus* species 6, *Ectocarpus* species 7, *Ectocarpus* species 5, *Ectocarpus* species 9, *E. siliculosus*, *Ectocarpus* species 3) and 12 to test the level of introgression between these two clades. Tests were designed using a four-taxon fixed phylogeny (((P1,P2)P3)O), where P1 and P2 are closely related species from the same clade, P3 is a more divergent species that may have experienced admixture with one or both of the (P1,P2) taxa, and an out-group (O). *E. fasciculatus* was used as the out-group taxon for all ABBA-BABA tests. Details about how P1, P2 and P3 taxa were selected for each test are given in Table S25. Previous results of species tree inference were used to inform subsequent ABBA–BABA tests and to define the (((P1,P2)P3)O) phylogenies. ABBAs are sites at which the derived allele (called B) is shared between the taxa P2 and P3, whereas P1 carries the ancestral allele (called A), as defined by the outgroup while BABAs are sites at which the derived allele is shared between P1 and P3, whereas P2 carries the ancestral allele. Under incomplete lineage sorting, conflicting ABBA and BABA patterns should occur in equal frequencies, resulting in a D statistic equal to zero. Historical gene flow between P2 and P3 causes an excess of ABBA, generating positive values of D. Historical gene flow between P1 and P3 causes an excess of BABA, generating negative values of D. Patterson’s D-statistic was calculated for the concatenated alignments of the 261 orthologroups. Significance was detected using a block-jackknifing approach^116–118^, with a block size of 5 Kbp. For the jackknife procedure, one block of adjacent sites was removed n times. A Z-score was finally obtained by dividing the value of the D statistic by the standard error over n sequences of 5 Kbp. The ParimonySplits network was reconstructed for the genus *Ectocarpus* using SplitsTree 4^73^ (version 4.14.6) with 1000 bootstrap replicates.

## QUANTIFICATION AND STATISTICAL ANALYSIS

All statistical analyses are described in detail in the relevant sections of the “Method details” section and the results of statistical tests are shown in the tables and figures.

## ADDITIONAL RESOURCES

The Phaeoexplorer website (https://phaeoexplorer.sb-roscoff.fr) provides access to all the annotated genome assemblies described in this study as downloadable files. The output files from the Orthofinder^32^, Interproscan^52^, Hectar^74^ and eggNOG-mapper^42^ analyses, together with the results of the various DIAMOND^26^ sequence similarity analyses (see section “Analyses aimed at deducing functional characteristics of predicted proteins”), can also be downloaded. In addition, the site provides genome browser interfaces for the genomes and multiple additional tools and resources including BLAST interfaces for genomes, proteomes and *de novo* transcriptomes, various experimental protocols, an RShiny-based transcriptomic aggregator for the model brown alga *Ectocarpus* species 7 strain Ec32 and a link to genome-wide metabolic networks for the Phaeoexplorer species.

Additional data have been deposited in the CNRS Research Data depository under the title “Data for Phaeoexplorer publication: Evolutionary genomics of the emergence of brown algae as key components of coastal ecosystems” (DOI: https://doi.org/10.57745/9U1J85).

## SUPPLEMENTAL RESULTS

### Structural features of brown algal genomes

The brown algal genomes generated by this project exhibit differences in assembly size and GC content (Figure S16A, Table S1). The cumulative lengths of intergenic regions, the cumulative lengths of transposable elements and cumulative intron size were all correlated with genome size (Figures S6A and S16B, Table S1). An earlier analysis of the *Ectocarpus* species 7 (at that time classified as *Ectocarpus siliculosus*) genome found a strong tendency for adjacent genes to be located on opposite strands of the chromosome, a feature that is more typical of very small, compact genomes^85^. Analysis of the Phaeoexplorer genomes indicated that this characteristic is common to the whole FDI (Fucophycideae/Dictyotales/Ishigeales) clade but the tendency is strongest in the Ectocarpales, where the proportion of alternating gene pairs can be greater than 60% (Figure S6A, Table S1). Another feature that is shared by all brown algae, plus some of the closely-related sister taxa, is that intergenic intervals between divergently transcribed gene pairs tend to be shorter than those between tandem gene pairs (Figure S6A, Table S1). Again, this is a feature that was previously observed in *Ectocarpus* species 7 where it is associated with the presence of common nucleosome-depleted regions and histone modification peaks at the transcription start sites of the divergently transcribed genes^119^. Tandemly-repeated genes are more numerous in the Phaeophyceae than in closely-related outgroup clades (Figure S6A, Table S1) and, although they only represent modest proportions of the total gene sets (around 6% for the Ectocarpales and 10-12% for the Fucales, for example), they may have played important roles in brown algal adaptation and diversification.

### Composite genes

Composite genes (i.e. genes derived from domain fusions and fissions) tended to be retained at a higher frequency than non-composite genes during the most recent phase of brown algal evolution but a larger proportion of these genes had unknown functions than for non-composite genes (Figure S17A). Figure S17B shows an example of a novel domain association that is predicted to have arisen just prior to the emergence of the brown algal lineage.

### Horizontal gene transfer

Genes that are predicted to be derived from HGT exhibit different ranges of codon usage bias and gene feature lengths to non-HGT-derived genes (Figure S17C), and have diverse predicted functions, with carbohydrate transport and metabolism and cell wall/membrane/envelope biogenesis being the most prevalent (Figure S17D).

### Brown algal introns and the spliceosome

An analysis of intron conservation profiles using a set of single-copy orthologous proteins conserved across Phaeophyceae and close sister groups indicated that more than 90% of brown algal introns are shared with at least one of the sister clades, and therefore that most introns are ancient. This analysis also confirmed that a high proportion of *C. australica* introns are unique to that species, suggesting that a burst of intron gain occurred during the evolution of this taxa (Figures S6B, S18A, S18B and S18C). Analysis of the *Ectocarpus* genomes showed that intron sizes, phases and positions have been strongly conserved across the genus (Figure S18D).

The vast majority of splice sites in Phaeophyceae are of the canonical GT-AG type (Figure S18C) and no signatures of minor U12 introns were identified in intronic sequences, in accordance with a previous study^120^. Consistent with this observation, an analysis of spliceosome proteins using a recent reference dataset^121^ found that, with the exception of ZCRB1, none of the 12 minor spliceosome genes were present in brown algal genomes (Table S26), indicating that the minor intron spliceosomal associated machinery was lost in the Phaeophyceae. As far as the Sm/Lsm spliceosome is concerned, this analysis identified lineage-specific gene duplications and loss of some components^122^ (Figures S18E and S18F).

### Long non-coding RNAs

Long non-coding RNAs (lncRNAs) play key regulatory roles in diverse eukaryotic taxa^123,124^. This class of gene has been characterised in *Ectocarpus* species 7 and *Saccharina japonica* and there is evidence that some of these genes may play regulatory roles during development and life cycle progression in brown algae^86,125,126^. Eleven transcriptomes (Table S27) were searched for lncRNAs using a meta-classifier, votingLNC, that integrates information from ten lncRNA-detection programs. This analysis indicated that brown algal genomes contain about twice as many lncRNA genes as protein-coding genes (Figure S19A). The lncRNA transcripts tend to be shorter than mRNAs and to be less GC-rich (Figure S19B).

### Gene loss from plastid genomes

Comparison of the plastid genomes generated by the Phaeoexplorer project (Table S28) identified gene loss events during the diversification of the brown algal orders that affected five different loci (cp74, rpl32, rbcR, rpl9 and syfB). Two of these genes have been lost more than once within the lineage (Figure S20A, Table S29). Gene losses have previously been reported^127,128^ for three of these genes (rpl32, rbcR and syfB).

### Evidence for mitochondrial introgression events between species

Phylogenetic trees based on either the plastid or mitochondrial gene sets were congruent with the phylogeny based on nuclear gene data^89^ at the ordinal level (Figure S20B). For the plastid genome, this congruency was also observed at the sub-ordinal level but, interestingly, a marked discordance was observed between the mitochondrial tree and the nuclear and plastid trees for the Ectocarpales, indicating possible mitochondrial introgression events between species within this order (Figure S20B), as has been observed in other eukaryotic lineages, for example *Picea*^129^ and chipmunks^130^.

### Metabolic pathway analysis using AuCoMe

The robust set of 24 core reactions found mainly in brown algae by the AuCoMe analysis (see main text) included reactions catalysed by indole-3-pyruvate oxygenases (EC 1.14.13.168) and a PQQ-dependent sugar dehydrogenase (Table S4). Indole-3-pyruvate oxygenase was previously thought to be absent from brown algae and it has been proposed that its activity could be complemented by associated bacteria^131^ to enable auxin biosynthesis. The identification of indole-3-pyruvate oxygenases by the current study indicates that a full auxin biosynthetic pathway may be present in brown algae. Concerning PQQ-dependent sugar dehydrogenase, the most similar bacterial protein is a membrane-bound glucose sorbosone dehydrogenase involved in vitamin C biosynthesis but the closest eukaryotic match, outside stramenopiles, is an animal hedghog-interacting protein. This suggests this the brown algal protein could either have a metabolic and/or a signalling function.

### Polyketide metabolism

Polyketides are a group of secondary metabolites that exhibit a wide range of bioactivities. Brown algae possess three classes of type III polyketide synthase (Figures 3A, 4A and S8E, Table S9). The first class (PKS1) was acquired early in stramenopile evolution, after divergence from the oomycetes but before the divergence of Diatomista and Chrysista (Figure 4A). Acquisition of PKS1 probably involved horizontal transfer from a bacterium. Distance analysis of a maximum likelihood phylogenetic tree (Figure 4A) indicated that PKS1 was more likely to have been acquired from a cyanobacterium than from an actinobacterium, as was previously proposed^132^. The second class of type III polyketide synthase (PKS2) probably arose via a duplication of PKS1 just prior to the divergence of the brown algal lineage from the Phaeothamniophyceae (Figure 4A). *Undaria pinnatifida* is the only species where a duplication of PKS2 was observed. The PKS3 class appeared much more recently, within the Ectocarpales during the diversification of this brown algal order, again probably via a duplication of PKS1 (Figure 4A). PKS3 genes were only found in two families, the Ectocarpaceae and the Scytosiphonaceae. The enzymatic activities and functions of PKS2 and PKS3 remain to be determined but analyses of the functions of recombinant PKS1 proteins have indicated that different proteins may have different activities leading to the production of different metabolites^132–134^. For example, the *Ectocarpus* species 7 PKS1 enzyme produces phloroglucinol from malonyl-CoA alone or malonyl-CoA plus acetyl-CoA whereas the PKS1 enzymes of *Sargassum binderi* and *Sargassum fusiforme* generate 4-hydroxy-6-methyl-2-pyrone. It is therefore interesting that the PKS1 family has expanded in some members of the Sargassaceae, with a maximum of five PKS1 genes detected in *S. fusiforme* (Figures 3A and 4A). These expansions may be associated with diversification of PKS1 biochemical function.

### Sulphatases

Sulphatases remove sulphate groups from sulphated polysaccharides^81^. The brown algal enzymes appear to be of quite ancient origin, with homologues being present in other stramenopile lineages (Table S30).

### Carbon storage via the mannitol cycle

In brown algae, the output of the Calvin cycle is used to produce the carbon storage molecule mannitol via the mannitol cycle^135^. The genes that encode the four enzymes of the mannitol cycle are not only conserved in all brown algae but were also found in taxa as distant as *H. akashiwo* (Table S14), indicating that the mannitol cycle was acquired before brown algae diverged from closely-related stramenopile taxa. The mannitol cycle genes include a linked pair of *M1PDH3* and *M1Pase1* genes which may therefore represent the ancestral core of this metabolic cycle. These pairs of genes are head-to-head and adjacent in brown algae with an intergenic region ranging from about 500 bp (Ectocarpales) to 10 kbp (*Fucus serratus*). The organisation of this gene pair differs in sister groups with, for example, a tandem arrangement and an intervening gene in *S. ischiensis* and two *M1PDH3* / *M1Pase1* gene pairs in *H. akashiwo* with tail-to-head and head-to-tail arrangements and long (150-200 kbp) intergenic regions. Several species within the Ectocarpales and Laminariales have acquired an additional *M1PDH* gene perhaps allowing greater capacity or finer regulation of carbon storage through the mannitol cycle (Table S14).

### LHCX light-harvesting proteins

In diatoms, LHCX proteins play an important role in the photoprotection of the photosynthetic apparatus under conditions of excess light^136^. Chromosome six of the *Ectocarpus* species 7 genome is known to contain a cluster of 14 LHCX genes interspersed with genes coding for proteins unrelated to photosynthesis^85,137^. Analysis of the Phaeoexplorer genomes indicated that LHCX genes were present in this genomic context before the divergence of the Desmarestiales but that the expansion of the gene cluster only began in a common ancestor of the Laminariales and Ectocarpales (Figure S12A). The general features of the cluster are conserved in the genomes of *Ectocarpus* species 7, *Ectocarpus crouaniorum*, *Ectocarpus fasciculatus*, *Scytosiphon promiscuus*, *Porterinema fluviatile* and *C. linearis* but in *C. linearis* the cluster appears to have undergone a partial rearrangement and contains only three LHCX copies whereas the clusters in all other species include at least seven LHCX genes. In *S. latissima*, the genes of the cluster were found in three short contigs that may be adjacent on the same chromosome. LHCX gene clusters are also present in *D. dichotoma* (4 genes on contig 3024, 3 on contig 458 and 2 on contig 1592) of the early branching order Dictyotales and in *S. ischiensis* (4 plus 2 genes on contig 28 and 2 genes on contig 64), but the genomic context of these clusters differs from that of the cluster found in *Ectocarpus* spp., indicating independent expansions of the LHCX family in these taxa. LHCX sequences from *S. ischiensis*, *D. dichotoma* and the genus *Ectocarpus* grouped into independent, species/genus-specific clusters in a phylogenetic analysis (Figure S12B), further supporting independent expansions of the LHCX gene family in these taxa. The LHCX clusters probably evolved by gene duplication and ectopic gene conversion. The involvement of gene conversion is indicated by the higher intraspecific identity of LHCX sequences between paralogs than between putative orthologs (based on synteny) compared across species (Figures S12A, S12B). The large gene cluster that contains LHCX4 to LHCX7 commonly contains overlapping LHCX genes on opposite strands of the chromosome, with some or all of the exons of one gene being located within an intron of the other (Figure S12C). Changes in gene dosage in response to environmental selection (as observed, for example, in studies by^138,139^) may have been a driver of the LHCX gene family expansion.

### Transcription factors

An analysis of transcription-associated proteins (TAPs) indicated that the range of gene families and the relative sizes of each gene family are relatively constant across the brown algae and their close sister species (Figures 3A, S21A, S21B and S21C, Table S15), suggesting that the emergence of the brown algae did not involve major changes to the transcription machinery. For example, large families of C2H2, Zn_clus, GNAT and SWI_SNF_SNF2 genes are a common feature in both brown algal and sister taxa genomes.

The number of TAP genes has remained relatively stable during the diversification of the brown algae (Figures 3A, S21B and S21C). This contrasts with what has been observed for the green lineage, where the size of TAP complement has been shown to be broadly correlated with morphological complexity^140^. Although the overall TAP complement has been stable, variations were detected in the sizes of some specific TAP families, including variations that correlated with the degree of morphological complexity. A broadly positive correlation was observed for the three-amino-acid length extension homeodomain (TALE HD) transcription factor (TF) families for example (Figure S11A, Table S15). These observations suggest that increased multicellular complexity in the brown algae may have required different combinations of TFs rather than simply increasing numbers. Two TALE HD TFs (OUROBOROS and SAMSARA) have been shown to be necessary for the deployment of the sporophyte developmental program in *Ectocarpus* species 7^141^. The *Ectocarpus* species 7 TALE HD TF family consists of three genes which are conserved across the brown algae^141^ but we detected a modest expansion of the gene family in some members of the Fucales, which have acquired one or two additional paralogues of the third TALE HD TF (Figure S11B).

### Ion channels

The Phaeoexplorer dataset allowed the origins of the major families of brown algal ion channels to be traced (Table S16). Most classes of brown algal ion channel are also found in other stramenopile groups, such as IP_3_ receptors, which were earlier shown to mediate calcium signalling in *Fucus*^142^, and Nav/Cav four-domain voltage-gated calcium channels, which are associated with rapid signalling processes and the regulation of flagella motion^143^. Pennate diatoms and angiosperms, which both lack flagellated gametes, do not possess Nav/Cav channels^144^. Land plants and diatoms also lack IP_3_ receptors. All stramenopile lineages, including brown algae, have OSCAs, which are mechanosensitive channels that have been implicated in calcium signalling during osmoregulation^145,146^, and transient receptor potential (TRP) channels (although the latter were not detected in *H. akashiwo*). In brown algae the only ubiquitous ligand-gated ion channels are the glutamate receptors (GLR). P2X receptors are absent and the pentameric ligand-gated ion channel (pLGIC, related to acetylcholine and GABA receptors in animals) was only found in *D. mesarthrocarpum*. EukCat, a novel class of single domain voltage-gated calcium channel recently discovered in diatoms^147^, appears to be absent from brown algae and other stramenopiles.

The Mitochondrial Calcium Uniporter (MCU) plays an important role in the uptake of calcium into mitochondria and is highly conserved across the eukaryotes. It is therefore surprising that the Fucales, like diatoms^148^, have lost this transporter (Table S16) and it is unclear how Fucales and diatoms regulate mitochondrial calcium, as alternative mechanisms are not known. Cyclic nucleotide gated channels, which are only present at low copy number, also appear to have been lost from several brown algal lineages, including kelps. In contrast, TRP channels are found in all brown algal genomes and the gene family is massively expanded in some lineages (Table S16). In animals, TRP channels are primarily involved in sensory functions (e.g. touch, temperature) and ligand-based activation^149^, so the brown algal channels may play a role in sensing the environment. TRP channels have been lost in land plants, which show an expansion of CNGCs and GLRs^150^.

### Membrane-localised signalling proteins

Brown algae have integrin-like molecules^151^, and this feature was found to be shared with several sister stramenopile lineages including pelagophytes. A single integrin-like gene was found in close sister groups but this gene was duplicated early in brown algal evolution, with a further recent duplication in *Ectocarpus* species 7 and *U. pinnatifida* and gene loss in several other brown algae (Figures 3A and S9A, Table S31). Proteins with an FG-GAP domain (a component of integrins) were found in several other stramenopile lineages but these proteins did not have the characteristic integrin domain structure and their evolutionary relationship to brown algal integrin-like proteins is uncertain (Figure S9B).

Brown algae and closely-related sister groups also possess a single gene that is very similar to the land plant membrane-localised calpain domain protein DEK1 (Figures 3A and S9A, Table S32), which is thought to act as a mechanosensor in land plants by activating the RMA channel^152^.

Fasciclin (FAS) domain proteins are found throughout the stramenopiles but only brown algae have membrane-localised proteins with four extracellular FAS domains analogous to the membrane-localised fasciclin proteins in animals that interact with the extracellular matrix (Figure S9A, Table S33)^153^.

All stramenopiles, including brown algae, appear to have a single tetraspanin gene plus a gene that encodes a stramenopile-specific eight transmembrane domain tetraspanin-like protein (Figure S9A, Table S34).

### Flagellar proteins

Most stramenopiles possess motile cells with two heteromorphic flagella, although there are some exceptions such as the male gametes of pennate diatoms. Searches for the presence of flagellar proteins, based on proteomic inventories of *Colpomenia bullosa* anterior and posterior flagella^102^, indicated that highly-conserved proteins such as the mastigoneme proteins MAS1, MAS2 and MAS3, a flagellar titin protein, which is thought to be involved in connecting mastigonemes to the flagellar axoneme, and flagellar creatine kinase are present in all brown algae and sister taxa (Figures 3A and 4B, Table S13). In contrast, the HELMCHROME protein, which is thought to be involved in light reception and zoid phototaxis^102^, was only found in brown algae and some closely-related sister taxa, the most distant being *C. australica* (Figure 3A).

### Histone proteins

The emergence of the brown algae also appears to have been associated with genomic events that may have had an important impact on the regulation of chromatin function. Most brown algal genomes code for about 20 histone proteins but the number of histone genes was correlated with genome size. For example, *U. pinnatifida* and *A. nodosum*, which have quite large genomes (634 and 1,253 Mbp, respectively) have 36 and 173 histone genes, respectively (Table S23). As in animal genomes^154^, the majority of brown algal histone genes occur in clusters. For example, *Chordaria linearis* and *U. pinnatifida* both have three contigs containing 80-90% of their histone genes, whereas *Saccharina latissima* and *S. ischiensis* each have a single contig which contains ∼40% of their histone genes. As in other eukaryotic lineages, histone H4 is highly conserved due to its structural role in binding H3, H2A and H2B, and no variants were found in brown algae. All Phaeophyceae possess H3.3 and CenH3 but additional H3 variants were only found in the Discosporangiales and Dictyotales and in sister taxa as distant as the raphidophytes (Figures 3A and S22A). The centromeric variant CenH3 is highly divergent across the brown algae, particularly in the N-terminal tail (Figure S22B), as reported in other lineages^155^. In contrast to what has been observed in humans and land plants^156,157^, H3.3 is encoded by a single gene in brown algae. H3.1 is encoded by several genes as in the majority of organisms. The brown algal H3.1 and H3.3 clades are distinct from those of animals and land plants indicating independent amplifications (Figure S22C). Linker H1 variants are highly divergent across the eukaryotes and this is also true for brown algae, with *A. nodosum,* for example, possessing 11 different H1 proteins (Figure S22D, Table S23). Brown algae with a genome of less than 1 Gbp possess ∼10 H2B-coding genes but *F. serratus*, *A. nodosum* and the sister taxon *H. akashiwo* have at least three times more. Finally, brown algae not only possess H2A variants such as the H2A.X but also three novel H2A variants that only exist in Phaeophyceae and the sister taxon Schizocladiophyceae (Rousselot et al, unpublished).

### Histone post-translational modification

An analysis of eight histone post-translational modifications in *Ectocarpus* species 7 indicated that the majority had similar genomic and genic distributions to those observed in animals and land plants, indicating that they have similar functional roles^119^. However, dimethylation of lysine 79 of histone H3 (H3K79me2) has a very different distribution pattern in *Ectocarpus* compared to animals. In animals, H3K79me2 is associated with the 5’ ends of transcribed genes^158^ whereas in *Ectocarpus* it occurs as extended blocks, which can span several genes and appears to be associated, at least indirectly, with down-regulation of gene expression^119^. The H3K79me2 mark is written by DOT1-class DNA methylases, which are encoded by multiple genes in brown algae. Dollo analysis identified gain of a new DOT1 orthogroup (OG0008134) in the common ancestor of the Phaeophyceae and the two close sister taxa *S. ischiensis* and *Phaeothamnion wetherbeei*, indicating a possible correlation between this unusual H3K79me2 distribution pattern and the appearance of a novel DOT1 protein.

### Genome changes associated with adaptation to freshwater environments

A small number of brown algal species have adapted to freshwater habitats, with each adaptation event occurring independently. The Phaeoexplorer dataset includes genomes for four freshwater species in addition to the previously sequenced *Ectocarpus subulatus*^159^. Adaptation to a freshwater habitat was correlated with loss of orthogroups (Figure S4A) and clear reductions in the sizes of some key gene families such as haloperoxidases (Figures 3A and S8C). In terms of gene gain, 12 orthogroups were found to be consistently over-represented when the genomes of each of the five freshwater species were compared with those of related seawater species, including orthogroups with genes involved in membrane transport and signalling. A transcriptomic analysis of *P. fluviatile* grown in either seawater or freshwater conditions identified 1,328 differentially-expressed genes (DEGs), with the genes upregulated in freshwater being enriched in DNA replication regulation and transcription functions (Table S24). Comparison of the differentially regulated gene set with a similar set generated for *E. subulatus* indicated that both sets of DEGs were enriched in lineage-specific genes, suggesting that, to a large extent, each adaptation to freshwater involved lineage-specific innovations (Table S35). Furthermore, a focused comparative analysis of metabolic networks (https://gem-aureme.genouest.org/fwgem/; Figure S7G) indicated that both *P. fluviatile* and *Pleurocladia lacustris* have lost alginate lyases and a glycosyltransferase GT18, related to polysaccharide metabolism (Table S22). This is in line with previous work showing specific cell wall remodelling in *E. subulatus*^160^. *P. fluviatile* and *P. lacustris* have also lost genes encoding ubiquitin-conjugating enzymes and this may have impacted protein degradation and thus osmolarity, or be linked to the preferential activation of ubiquitin-independent regulatory mechanisms in situations of osmotic stress^145^. In addition to the shared gene losses, both species have independently lost genes. The strictly freshwater *P. lacustris* has lost several phosphatases, peptidases and aldolases, perhaps due to a reduced need to produce osmolytes, while the brackish water *P. fluviatile* has lost several enzymes related to phenol metabolism and a magnesium transporter. More surprisingly, the two species have lost different sets of enzymes involved in RNA modifications, which may either indicate convergent acquisition of different regulatory mechanisms, or may simply be markers of phylogenetic divergence.

### Comparative analysis of *Ectocarpus* genomes

The genus *Ectocarpus* includes 16 cryptic species^161^. Twenty-two new strains of *Ectocarpus*, representing 12 new species, were sequenced in this study. Comparison of high continuity *Ectocarpus* assemblies indicated that *Ectocarpus* spp. genomes share extensive synteny (Figure S23A, Table S36).

The genus *Ectocarpus* belongs to the order Ectocarpales, which emerged late in brown algal evolution but has become the most species-rich brown algal order, with over 750 species^162^. The Ectocarpales exhibited the same general phenomenon of orthogroup loss as the other Phaeophyceae orders during the diversification of the brown algae. However, this trend was not uniform within the Ectocarpales and a period of orthogroup gain is predicted to have occurred during the early emergence of the genus *Ectocarpus*, analogous to the burst of gene gain observed during the early emergence of the brown algal lineage (Figure S23B). The sets of both gained and conserved orthogroups were significantly enriched in transcription-related functions (Figure S23C). An analysis of the *Ectocarpus* species 7 reference genome orthologues indicated that rapidly-evolving, taxonomically-restricted genes (Group 5 based on the phylostratigraphy analysis: Table S2) are most abundant on the sex chromosome (Figure S23D), an observation that is consistent with other recent findings^163^. These *Ectocarpus* genus-specific genes exhibit higher dN and dS values than genes that the genus shares with other brown algae (Figure S23E, Table S18).

### Hormone biosynthesis pathways

Brown algal genomes contain homologues of components of several phytohormone biosynthetic pathways, including indole-3-acetic acid (the auxin IAA), cytokinins, brassinosteroids, ethylene and gibberellins (Figure S24, Table S37), with the exception of tryptophan aminotransferase (TAA; auxin biosynthesis) and copalyl diphosphate/ent-kaurene synthase and ent-kaurene oxidase (CPS/KO; gibberellin biosynthesis). The presence of these genes is consistent with the identification of the corresponding hormones^164,165^ and of brassinosteroid-like molecules^166^ in brown algae. As far as abscisic acid (ABA) is concerned, brown algae have the capacity to synthesise oxidised cleaved carotenoids, supporting the idea that they may generate ABA-like signalling molecules^167^. Similarly, OPDA, an intermediate of the jasmonic acid pathway, has been identified in brown algae, raising the possibility that this molecule itself may have a signalling role similar to OPDA-like molecules in hornworts^168^. The putative brown algal phytohormone biosynthetic genes are also present in the genomes of sister taxa and therefore appear to have been acquired well before the emergence of the brown algae. However, an expansion of the flavin monooxygenase (YUC) gene family (which mediates a rate limiting step in IAA production in land plants) appears to have occurred in the common ancestor of the Phaeophyceae, *S. ischiensis, C. australica* and *T. minus* (Figure S24). Several gene families that are putatively associated with the biosynthesis of phytohormones such as ABA, brassinolide, ethylene, gibberellins, jasmonic acid and strigolactones have expanded in the Fucales (Figure S24).

## Notes

https://phaeoexplorer.sb-roscoff.fr

https://doi.org/10.57745/9U1J85

https://www.ebi.ac.uk/ena/browser/view/PRJEB76691

### Competing Interest Statement

The authors have declared no competing interest.

### Summary of Updates

Author modification: "Seven T. LoDuca" should be "Steven T. LoDuca". Table S1 updated with accession numbers.

