## Supplementary material for "Evolutionary genomics of the emergence of brown algae as key components of coastal ecosystems": Table S25

| P1_P2_P3 | group | ABBA | BABA | D |
| --- | --- | --- | --- | --- |
| sp7_sp5_sp6 | clade_2 | 24069.63 | 24141.19 | -0.26 |
| sp7_sp6_sp9 | clade_2 | 24648.30 | 24646.74 | 0.01 |
| sp5_sp6_sp9 | clade_2 | 24641.30 | 24657.63 | -0.09 |
| sp6_sp7_sp9 | clade_2 | 24646.74 | 24648.30 | -0.01 |
| sp7_sp5_sp9 | clade_2 | 24572.33 | 24554.44 | 0.13 |
| sp5_sp9_Esil | clade_2 | 24774.81 | 24746.81 | 0.07 |
| sp7_sp9_Esil | clade_2 | 24755.11 | 24717.00 | 0.10 |
| sp6_sp9_Esil | clade_2 | 24866.07 | 24821.74 | 0.11 |
| sp7_sp5_Esil | clade_2 | 25032.00 | 25021.89 | 0.07 |
| sp6_sp5_Esil | clade_2 | 25121.44 | 25105.11 | 0.09 |
| sp6_sp7_Esil | clade_2 | 25113.15 | 25106.93 | 0.04 |
| sp9_Esil_sp3 | clade_2 | 24675.00 | 24760.56 | -0.18 |
| sp5_Esil_sp3 | clade_2 | 24795.30 | 24868.41 | -0.14 |
| sp7_Esil_sp3 | clade_2 | 24780.26 | 24840.15 | -0.12 |
| sp6_Esil_sp3 | clade_2 | 24882.93 | 24948.26 | -0.12 |
| sp5_sp9_sp3 | clade_2 | 24773.78 | 24761.33 | 0.04 |
| sp7_sp9_sp3 | clade_2 | 24756.15 | 24730.48 | 0.08 |
| sp6_sp9_sp3 | clade_2 | 24860.89 | 24840.67 | 0.06 |
| sp7_sp5_sp3 | clade_2 | 25039.78 | 25026.56 | 0.12 |
| sp6_sp5_sp3 | clade_2 | 25119.37 | 25111.59 | 0.05 |
| sp6_sp7_sp3 | clade_2 | 25114.70 | 25120.15 | -0.04 |
| sub_Ecro_sp2 | clade_1 | 26007.07 | 26066.96 | -0.09 |
| sub_Ecro_sp1 | clade_1 | 25804.85 | 25873.30 | -0.12 |
| sp2_sp1_Ecro | clade_1 | 26033.78 | 26033.00 | 0.00 |
| sp1_sp2_sub | clade_1 | 26213.44 | 26222.78 | -0.04 |
| sp2_sp1_sp3 | clade_1_2 | 25976.48 | 25962.48 | 0.06 |
| sub_Ecro_sp3 | clade_1_2 | 25749.37 | 25785.15 | -0.08 |
| sp2_sp1_Esil | clade_1_2 | 26139.81 | 26123.48 | 0.07 |
| sub_Ecro_Esil | clade_1_2 | 25954.70 | 25913.48 | 0.09 |
| sp2_sp1_sp9 | clade_1_2 | 26016.15 | 25995.93 | 0.10 |
| sub_Ecro_sp9 | clade_1_2 | 25807.96 | 25814.19 | -0.01 |
| sp2_sp1_sp5 | clade_1_2 | 26142.41 | 26103.52 | 0.16 |
| sub_Ecro_sp5 | clade_1_2 | 25912.44 | 25936.56 | -0.05 |
| sp2_sp1_sp7 | clade_1_2 | 26115.44 | 26091.33 | 0.11 |
| sub_Ecro_sp7 | clade_1_2 | 25898.70 | 25910.37 | -0.03 |
| sp2_sp1_sp6 | clade_1_2 | 26214.48 | 26196.59 | 0.08 |
| sub_Ecro_sp6 | clade_1_2 | 25995.93 | 26019.26 | -0.05 |
