## Supplementary material for "Evolutionary genomics of the emergence of brown algae as key components of coastal ecosystems": Table S35

|  | **Upregulated in seawater** | | | **Upregulated in freshwater** | | |
| --- | --- | --- | --- | --- | --- | --- |
|  | ***E. subulatus*** | ***P. fluviatile*** | **common** | ***E. subulatus*** | ***P. fluviatile*** | **common** |
| **Lineage-specific genes** | 20% (411/2021) | 6.7% (292/4344) |  | 10% (210/2021) | 8.0% (346/4344) |  |
| **Shared orthogroups** | 12% (791/6606) | 3.8% (254/6606) | 0.54% (36/6606) | 18% (1169/6606) | 3.0% (195/6606) | 0.35% (23/6606) |
